# SEQing: web-based visualization of iCLIP and RNA-seq data in an interactive python framework

**DOI:** 10.1101/2019.12.17.865873

**Authors:** Martin Lewinski, Yannik Bramkamp, Tino Köster, Dorothee Staiger

## Abstract

**Background:** RNA-binding proteins interact with their target RNAs at specific sites. These binding sites can be determined genome-wide through individual nucleotide resolution crosslinking immunoprecipitation (iCLIP). Subsequently, the binding sites have to be visualized. So far, no visualization tool exists that is easily accessible but also supports restricted access so that data can be shared among collaborators.

**Results:** Here we present SEQing, a customizable interactive dashboard to visualize crosslink sites on target genes of RNA-binding proteins that have been obtained by iCLIP. Moreover, SEQing supports RNA-seq data that can be displayed in a different window tab. This allows, e.g. crossreferencing the iCLIP data with genes differentially expressed in mutants of the RBP and thus obtain some insights into a potential functional relevance of the binding sites. Additionally, detailed information on the target genes can be incorporated in another tab.

**Conclusion:** SEQing is written in Python3 and runs on Linux. The web-based access makes iCLIP data easily accessible, even with mobile devices. SEQing is customizable in many ways and has also the option to be secured by a password. The source code is available at https://github.com/malewins/SEQing.

## Background

RNA-binding proteins (RBPs) play a key role in orchestrating the transcriptome at the posttranscriptional level. To unravel posttranscriptional networks controlled by RBPs, RNAs associated with RBPs *in vivo* are recovered by immunoprecipitation of the RBP and high throughput sequencing of the co-precipitated RNAs. Recently, we have adapted individual nucleotide resolution crosslinking immunoprecipitation (iCLIP) originally developed for mammals [1–3] for the use in the reference plant *Arabidopsis thaliana* [4].

After processing sequencing data, a common bioinformatician’s task is the visualization of genomic windows or genes of interest, i.e. targets of RBPs in the case of iCLIP data. Genomic windows are frequently visualized as stacked charts alongside the gene model. To accomplish this, tools have been developed to support scientists without a programming background to visualize their datasets. The visualization tools either produce static or dynamic (interactive) visualizations. Each of these tools serves a particular purpose, e.g. exploration, documentation or the production of customized figures. Local (installable) tools run on the user’s machine and can import sequencing files which are stored in a secure location. Some of the local tools aim to produce static visualizations by using a set of genomic coordinates and commonly available programming languages like R or Shell script. Gviz [5] and ggbio [6] are both R packages to produce images of genomic windows and can be customized to answer particular scientific questions. These images can then be shared with co-workers or used in publications. Svist4get [7] is a Python package to create figures of genomic windows simply by using the command line. This is convenient for automated use, for example as part of a pipeline, and also does not require knowledge of the Python language. But images lack the ability to explore the data interactively, and these tools often cannot be handled by users without a programming background. IGV [8] and IGB [9] are interactive genome browsers which can be installed on a local computer and require no programming knowledge. Both support the most common file formats from high-throughput datasets and offer a variety of customization options. The downside of locally installed tools is the lack of the ability to share the interactive views of genomic windows. Each user needs to download the identical, and in most cases fairly large files, to import them into the tool. An alternative to local genome browsers are web-based genome browsers like the UCSC Genome Browser [10] and Zenbu [11]. These genome browsers also allow exploration on an interactive level, but can be accessed via a web-browser. The limitation of both online tools is the number of available genomes, as neither of them provides e.g. plant genomes. The existing gene annotations can be arranged online by the user, together with data tracks of various origins. Users can also upload custom tracks to inspect published or ongoing results and share them with colleagues. However, reservations may exist to upload data of ongoing research to remote servers which do not allow restricted access. Therefore we propose a tool to visualize iCLIP data which combines the accessibility of web-based solutions and the data safety of local tools, designated SEQing.

## Implementation

SEQing is implemented in Python 3 (3.5 and above) and publicly available on github (https://github.com/malewins/SEQing). It requires few pypi packages (dash, plotly and pandas). For other dependencies please see the requirements file inside the repository. The source code and sample data are provided in **Additional file 1**. Installation and usage instructions are supplied in **Additional file 2**, as well as on the github webpage. Pre-processing of the input files is not part of SEQing and can be accomplished beforehand with freely available toolkits, e.g. bedtools [12].

SEQing utilizes one of the recently developed Python packages for interactive data visualization named Dash [13]. This framework generates javascript from python code and hosts interactive dashboards on the local machine which are remotely accessible via a link in the browser, e.g. https://192.168.0.1:8060. The port number in the link can be set during startup of a Dash application and the IP address is dependent on the network setting of the hosting machine. Dash provides different types of basic components (e.g. buttons, dropdowns and tables), whereas the graphs are generated using the Plotly library [13]. The current viewport can be exported in common figure formats (PNG and SVG) using the native image export button. The interplay between different components of the dashboard (buttons and plots) is realized using callback functions delivered by Dash.

A schematic overview of the SEQing data flow, as well as the visualization goals in each tab, are displayed in Figure 1. Upon initialization, all input files must pass through a validator to check the consistency of the files before running the application. After passing the validation step, large files (e.g. annotation or data tracks) are saved locally in binary format via the python pickle library to vastly speed up future restarts of the dashboard. Optionally, a login window can be activated to restrict access to specific users.

**Figure 1.**
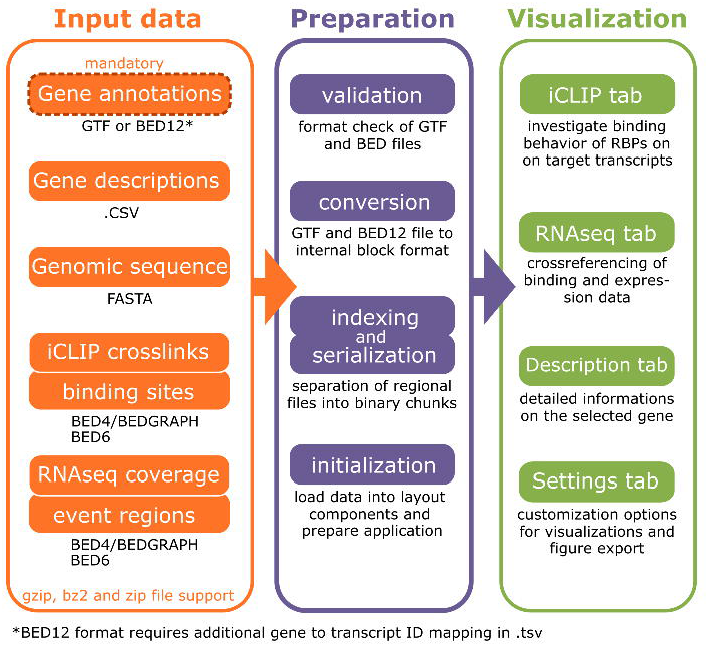
Schematic overview of data flow in SEQing. The displayed columns represent the three domains, input data (orange), preparation (purple) and visualization (green) of the SEQing data flow, respectively. Arrows between the columns indicate the direction of the data flow.

The minimal startup requirement for SEQing is a gene regional annotation file in `BED12` or `GTF` format. In the case of `BED12` an additional key-value table in `.tsv` is required to provide gene to transcript-id mapping. Running SEQing with the full feature set requires, in addition to the gene regional annotation, iCLIP crosslink sites in `BEDGRAPH` format, significant crosslink sites in `BED6` format, short text annotations of each gene in Tab-separated values (`.tsv`), extended gene annotations in `.tsv`, the genomic sequence for each transcript in `FASTA` format, RNA-seq coverage in `BEDGRAPH` format, and supplementary regional annotations for the coverage in BED6 format. If necessary, multiple gene annotation files can be loaded into SEQing, together with the corresponding sequence files in the described formats. BED, GTF or FASTA files can be provided in common compressed file formats, like .zip, .gz or .bz2, to save local disk space.

## Results and discussion

The features of SEQing are presented by running the tool with sample data from [4] on a local linux machine. We show common application cases of SEQing: browsing genes and their corresponding genomic windows. The source code and sample data can be downloaded from github or cloned directly into a local directory by using git clone https://github.com/malewins/SEQing.git. All dependencies to run SEQing can be installed by executing `pip3 install -r requirements.txt` in the SEQing directory. After startup, the application is accessible via a web-browser by entering the IP address of the host with a specified port number (defaults is 8060), e.g. https://192.168.0.1:8060. This is possible on the same machine or from remote machines inside the network.

### Visualization of Protein-RNA interactions

To showcase SEQing we utilize datasets with binding targets and crosslink sites of the RBP *Arabidopsis thaliana* glycine-rich RNA-binding protein 7 (*At*GRP7) [4]. Because *At*GRP7 is controlled by the circadian clock [14, 15], iCLIP was performed on plants harvested at the circadian maximum of *At*GRP7 expression, 36 h after transfer of the plants to continuous light (LL36) and at the circadian minimum of *At*GRP7 expression, 24 h after transfer of the plants to continuous light (LL24) [4]. The uniquely mapped reads were reduced to the position 1 nucleotide upstream of the read start (crosslink site) and piled up at each nucleotide position. We refer to transcripts with significant crosslink sites as *targets* of the RBP. The resulting value represents the number of crosslinks captured by iCLIP. A screenshot of SEQing displaying crosslink sites and significant crosslink sites from this dataset (GSE99427) is shown in Figure 2. It shows *KIN1* (AT5G15960), one of the target transcripts of *At*GRP7. We refer to transcripts with significant crosslink sites as *targets* of the RBP. Significant crosslink sites in this dataset were determined as described in König et al. [1] with the modifications that the threshold of the FDR was set to < 0.01 instead of < 0.05 and that crosslink sites had to be present at the same nucleotide in all but one biological replicate [4].

**Figure 2.**
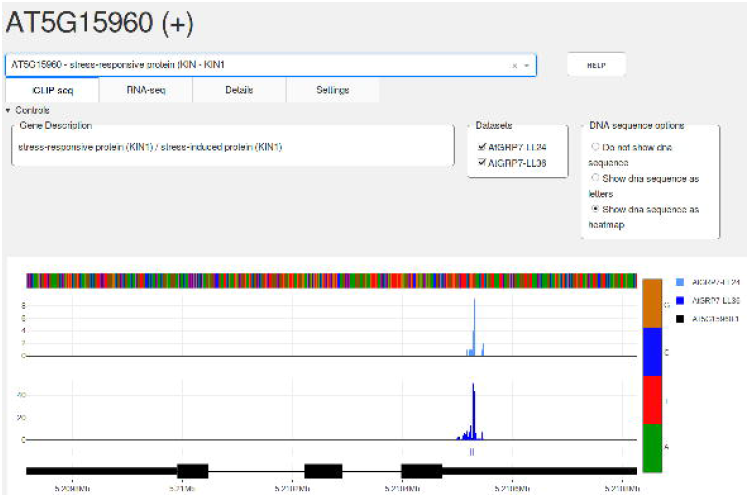
Screenshot of the SEQing iCLIP tab. The currently selected gene identifier with corresponding strand information in brackets is displayed at the top. Genes of interest can be selected in the dropdown menu, which also supports free text search. The four tabs below contain the main features of SEQing, the display of iCLIP (currently active) and RNA-seq data, as well as additional details on the selected gene and settings to customize plot appearances. The help button provides basic instructions to the user. The data shown originates from the included example data set, which is a subset of the GSE99427 dataset, visualized using the sources from Additional File 1.

The accession-id and strand information of the currently selected gene reside at the top of the dashboard. Genes of interest can be selected via a dropdown menu, also positioned at the top. The dropdown supports a free text search for gene identifiers and descriptions which have been supplied at the start (`-desc` parameter). After a gene is selected, the corresponding gene description is displayed inside the *Gene Description* field as well as the matching data tracks in the plot area below the control panel. The imported datasets are selectable in the adjacent *Datasets* area and the display modes of genomic sequences in the field *DNA sequence options*. By default, the sequences are displayed in heatmap mode, which can be changed to text mode or disabled (hidden). The legend at the right side of the plots shows the names and colors for each data track and gene isoform. Gene models on the same strand are colored black, whereas genes on the opposite strand are colored grey. The gene models are displayed in the 5’ → 3’ direction, i.e. if the selected gene resides on the forward strand (+ in description on top) a gene in opposite direction overlapping the selected gene will be shown in grey, and vice versa. The gene annotation track displays the annotated gene models, where thin lines represent introns and thick lines exons. If the selected gene is a protein coding gene with annotated untranslated regions, thinner bars represent untranslated regions and thicker bars represent coding regions. The dashboard will also provide the user with additional information about the selected gene in the corresponding *Details* tab (Supplemental Figure 1). This place is reserved for tables provided with the `-adv_descr` parameter containing e.g. complete or extended gene descriptions, synonyms, gene ontologies or known interaction partners of the selected gene. The appearance of the plots can be customized in the *Settings* tab (Supplemental Figure 2).

As an additional example we visualize crosslink sites from a public human dataset (GSE99700) using SEQing. This dataset contains, among others, *in vitro* iCLIP data with crosslink sites of the human splicing factor U2 Auxiliary Factor 2 (U2AF2) in the presence (GSM2650339) or absence (GSM2650359) of Far Upstream Element Binding Protein 1 (FUBP1) [16]. BED files from this dataset were imported into SEQing conjoined with the corresponding gene annotation (downloaded from ftp://ftp.ensembl.org/pub/). Supplemental Figure 3 depicts a genomic window from one of the U2AF2 targets, Polypyrimidine Tract Binding Protein 2 (PTBP2), similar to Figure 6 from Sutandy et al. [16]. The area marked by the arrow points to a binding site that is only detected when FUBP1 is present.

### Display of coverage tracks and splice events from RNA-seq

Additional information about iCLIP targets can be inferred from functional data, e.g. RNA-seq data from loss-of-function or gain-of-function mutants of the corresponding RBP (Figure 3). Here, we use RNA-seq data from *atgrp7* mutants and plants constitutively overexpressing *At*GRP7 (*At*GRP7-ox plants) compared to wild type plants. The samples were again harvested either at LL36 or LL24. The read coverage for each sample is presented as an area graph, combined with annotations below, in this case significant splice events. The splice events shown were determined with SUPPA2 [17] and transformed into BED6 format. The colors represent different event classes of SUPPA2, the alternative 3’ splice site (A3), alternative 5’ splice site (A5), and intron retention (RI) events. We refer to a significant splice event if a change of percent spliced-in (PSI) ratio was greater than 0.1 and the p-value of this event was < 0.01 when comparing the mutants to the wild type. Comparisons of the sample graphs yield information on genes differentially expressed or alternatively spliced in response to reduced or elevated *At*GRP7 levels. Crosslink sites in the vicinity of alternative splicing events may hint at a regulation of the splicing event by the RBP. The annotation bars below the coverage have three display modes. In the default setting, bars are plotted in blue below the corresponding coverage track. The second option paints the bars corresponding to the supplied text in the name field of the BED6 file, and the last option offers a score-dependent color gradient to display e.g. a score given in floating point numbers. The gene annotation track at the bottom is coherent with the iCLIP tab.

**Figure 3.**
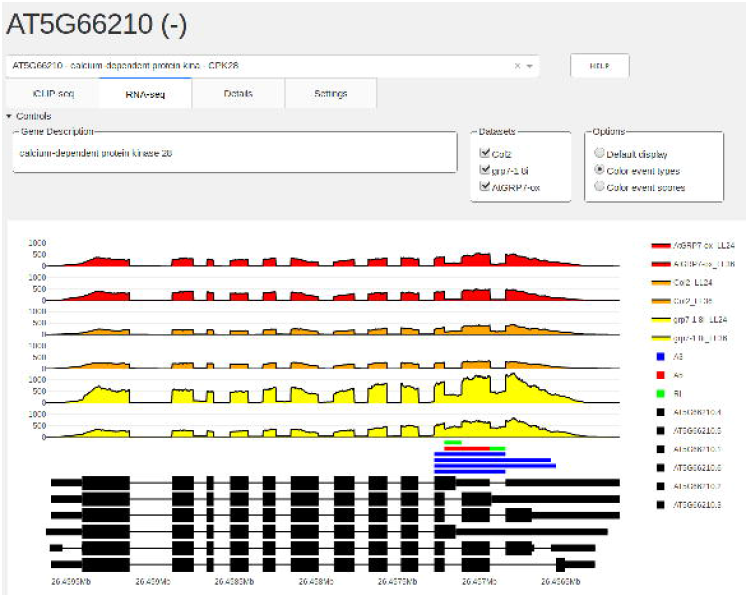
Screenshot of the SEQing RNA-seq tab. The RNA-seq tab shows coverage tracks of difierent samples derived from the dataset GSE99615 and significant splice events determined with SUPPA2 on the *CPK28* transcripts. The event types shown correspond to A3 (alternative 3’ splice sites) in blue, A5 (alternative 5’ splice sites) in red and RI (intron retentions) in green. The transcript isoforms are positioned below the coverage tracks. Besides displaying a brief description, the control panel allows the user to select which samples to plot and set the display mode of splice events. This screenshot was produced using the SEQing version provided in Additional File 1.

## Conclusions

Results from high-throughput sequencing need to be easily accessible for wet-lab scientists and safely shareable with co-workers. Especially the latter is important for ongoing research. The purpose of SEQing is to visualize iCLIP data sets and to make them accessible for wet-lab scientists. Upon start, it creates an interactive visualization platform whilst running on a local Linux machine. Group members can access iCLIP data, browse genes, and create figures by entering the link of the dashboard into their browser. The dashboard is multi-user capable, supports access over mobile devices (Android and iOS tested) and can be password protected (`-pwd` parameter). This combination of features is unique for SEQing and supports data safety as well as a convenient web-access.

## Availability and requirements

**Project name:** SEQing

**Project home page:** http://github.com/malewins/SEQing

**Operating system:** Linux (start), Any (browsing)

**Programming language:** Python 3

**Other requirements:**Python modules: plotly, dash, pandas (others see requirements.txt in github repository)

**License:**MIT

## Author’s contributions

ML provided the concept and contributed to parts of the source code. YB implemented most of the source code and documentation. TK and ML tested the software. ML and DS wrote the manuscript. All authors read and approved the manuscript.

## Acknowledgments

We thank Lena Raupach for implementing the initial version of the RNA-seq tab as part of a programming course.

## Ethics approval and consent to participate

Not applicable.

## Consent for publication

Not applicable.

## Availability of data and materials

The current study generated no datasets. A sample dataset containing a full sample track set and gene annotations is stored within the package’s subfolder `example_set`. The sample data originates from published *Arabidopsis thaliana* iCLIP (GSE99427) and RNA-seq (GSE99615) experiments.

## Funding of data and materials

This project was supported by the German Research Foundation (grants STA 653|9-1 and STA 653|14-1) to DS.

## Additional Files

Additional file 1 — Source code and sample data for SEQing

The Python code and samples of *Arabidopsis thaliana* iCLIP (GSE99427) and RNA-seq (GSE99615) data used to start the sample dataset dashboard. (.zip)

Additional file 2 — Installation and usage instructions

SEQing installation instructions and command line options with use cases and descriptions. (.pdf)

Additional file 3 — Supplemental figures containing screenshots of the Details tab, Settings tab and a visualization example of a public human dataset (GSE99700)

Screenshots of SEQing’s Details tab to deliver additional information on the selected gene to the browsing user (Supplemental Figure 1) and customization options in the Settings tab (Supplemental Figure 2). The screenshots were taken using the sample data incorporated in the github repository. Supplemental Figure 3 shows crosslink sites from a public human dataset (GSE99700) visualized using SEQing. (.pdf)

## References

1. König, J., Zarnack, K., Rot, G., Curk, T., Kayikci, M., Zupan, B., Turner, D.J., Luscombe, N.M., Ule, J.: iCLIP reveals the function of hnRNP particles in splicing at individual nucleotide resolution. Nature structural & molecular biology 17(7), 909–915 (2010)

2. Rossbach, O., Hung, L.-H., Khrameeva, E., Schreiner, S., König, J., Curk, T., Zupan, B., Ule, J., Gelfand, M.S., Bindereif, A.: Crosslinking-immunoprecipitation (iCLIP) analysis reveals global regulatory roles of hnRNP L. RNA Biology 11(2), 146–155 (2014)

3. Brugiolo, M., Botti, V., Liu, N., Müller-McNicoll, M., Neugebauer, K.M.: Fractionation iCLIP detects persistent SR protein binding to conserved, retained introns in chromatin, nucleoplasm and cytoplasm. Nucleic acids research 45(18), 10452–10465 (2017)

4. Meyer, K., Köster, T., Nolte, C., Weinholdt, C., Lewinski, M., Grosse, I., Staiger, D.: Adaptation of iCLIP to plants determines the binding landscape of the clock-regulated RNA-binding protein AtGRP7. Genome biology 18(1), 204 (2017)

5. Hahne, F., Ivanek, R.: Visualizing Genomic Data Using Gviz and Bioconductor. Methods in molecular biology (Clifton, N.J.) 1418, 335–351 (2016)

6. Yin, T., Cook, D., Lawrence, M.: ggbio: an R package for extending the grammar of graphics for genomic data. Genome biology 13(8), 77 (2012)

7. Egorov, A.A., Sakharova, E.A., Anisimova, A.S., Dmitriev, S.E., Gladyshev, V.N., Kulakovskiy, I.V.: svist4get: a simple visualization tool for genomic tracks from sequencing experiments. BMC bioinformatics 20(1), 113 (2019)

8. Thorvaldsdóottir, H., Robinson, J.T., Mesirov, J.P.: Integrative Genomics Viewer (IGV): high-performance genomics data visualization and exploration. Briefings in bioinformatics 14(2), 178–192 (2013)

9. Freese, N.H., Norris, D.C., Loraine, A.E.: Integrated genome browser: visual analytics platform for genomics Bioinformatics Oxford (England) 32(14), 2089–2095 (2016)

10. Fujita, P.A., Rhead, B., Zweig, A.S., Hinrichs, A.S., Karolchik, D., Cline, M.S., Goldman, M., Barber, G.P., Clawson, H., Coelho, A., Diekhans, M., Dreszer, T.R., Giardine, B.M., Harte, R.A., Hillman-Jackson, J., Hsu, F., Kirkup, V., Kuhn, R.M., Learned, K., Li, C.H., Meyer, L.R., Pohl, A., Raney, B.J., Rosenbloom, K.R., Smith, K.E., Haussler, D., Kent, W.J.: The UCSC Genome Browser database: update 2011. Nucleic acids research 39(Database issue), 876–82 (2011)

11. Severin, J., Lizio, M., Harshbarger, J., Kawaji, H., Daub, C.O., Hayashizaki, Y., Bertin, N., Forrest, A.R.R.: Interactive visualization and analysis of large-scale sequencing datasets using ZENBU. Nature biotechnology 32(3), 217–219 (2014)

12. Quinlan, A.R., Hall, I.M.: BEDTools: a flexible suite of utilities for comparing genomic features. Bioinformatics Oxford (England) 26(6), 841–842 (2010)

13. Inc., P.T.: Collaborative Data Science. https://plot.ly

14. Staiger, D.: RNA-binding proteins and circadian rhythms in Arabidopsis thaliana. Philosophical transactions of the Royal Society of London. Series B, Biological sciences 356(1415), 1755–1759 (2001)

15. Heintzen, C., Nater, M., Apel, K., Staiger, D.: AtGRP7, a nuclear RNA-binding protein as a component of a circadian-regulated negative feedback loop in Arabidopsis thaliana. Proceedings of the National Academy of Sciences of the United States of America 94(16), 8515–8520 (1997)

16. Sutandy, F.X.R., Ebersberger, S., Huang, L., Busch, A., Bach, M., Kang, H.-S., Fallmann, J., Maticzka, D., Backofen, R., Stadler, P.F., Zarnack, K., Sattler, M., Legewie, S., König, J.: In vitro iCLIP-based modeling uncovers how the splicing factor U2AF2 relies on regulation by cofactors. Genome research 28(5), 699–713 (2018)

17. Trincado, J.L., Entizne, J.C., Hysenaj, G., Singh, B., Skalic, M., Elliott, D.J., Eyras, E.: SUPPA2: fast, accurate, and uncertainty-aware differential splicing analysis across multiple conditions. Genome biology 19(1), 40 (2018)

