## Additional File 1 for "SEQing: web-based visualization of iCLIP and RNA-seq data in an interactive python framework": app_layout.py

```
#!/usr/bin/env python
# -*- coding: utf-8 -*-

""" Interactive visualizaton for iClIP-Seq and RNA-Seq data"""
import dash
import dash_core_components as dcc
import dash_html_components as html
import dash_auth
from textwrap import dedent
from app import app
import rna_tab
import iclip_tab
import settings_tab
import description_tab
import cfg

__author__ = "Yannik Bramkamp"


if __name__ == '__main__':
    if 'dropList' not in globals():
        print('Please start the program via validator.py')
        exit()
    # Properly define all global variables that are handed to this module by validator.py
    cfg.init(globals())

    try:
        myfile = open ("../help_text.md", "r")
        helpText = myfile.read()  
    except FileNotFoundError:
        helpText = '''
                ##### General
                
                Welcome to SEQing, an interactive, web based visualisation and exploration tool for iCLIP-seq and RNA-seq data.
                Use the drop down menu at the top to select your gene of interest. You can search by gene identifier,
                and depending on provided data, also by gene name and partial descriptions. 
                For more detailed information on the different tabs please consult the following paragraphs.
                
                ##### iCLIP-seq
                
                In this tab you can explore raw iCLIP-seq data and, if available, also predicted binding sites. Besides the basic
                interactive plot controls provided by the Dash framework, which you can access by hovering over the graphs,
                there are two main control elements:
                  -    On the left side you have checkboxes to select which datasets you wish to display, if more than one was provided to the tool.
                  -    On the right side, if dna sequence data was provided, you can select the display mode for said sequences. You can choose from
                       heatmap, letters, and no display at all. heatmap is strongly recommended for interactive use, as letters has a signifficantly
                       higher performance impact and is recommended only for the creation of static images.
                    
                ##### RNA-seq
                
                In this tab you can view RNA-seq coverage plots as well as splice events, if the necessary data was provided.
                Use the checkboxes in the Datasets panel to select which plots you want to view. The options panel allows you to select 
                between different display variants: Default, event type based cooring and score based coloring.
                
                ##### Details
                
                In this tab you can view further information on your selected gene. Which information is available depends on what your administrator has provided
                when setting up the tool.
                
                ##### Settings
                
                Here you can select colors for the graphs in the iCLIP-seq tab. Select a dataset from the dropdown, choose your color using
                the sliders and hit confirm. Should the plot legend elements overlap you can change the
                 distance between the colorbar and the trace legend with the colorbar margin slider. You can also select your desired format for
                 image export, currently png and svg are supported.
                '''
    
    
    

    # Set defaults if no advanced descriptions are available
    if cfg.advancedDesc is None:
        print('Advanced description DF is None')
        advList = []
        advStart = None
        advDisabled = True
    else:
        advList = list(cfg.advancedDesc.columns.values)
        advList.remove('gene_ids')
        advDisabled = False
    
    imgFormat = 'svg' # Default format for image export
    
    # Hide sequence related controls if no sequence data is available
    if len(cfg.sequences) == 0:
        seqDispStyle = {'display': 'none', 'height' : '100%', 'width' : '20vw'}
    else:
        seqDispStyle = {'height' : '100%', 'width' : '20vw'}
    if len(cfg.dataSetNames) == 0:
        dataSetStyle = {'display': 'none', 'height' : '100%', 'width' : '15vw'}
    else:
        dataSetStyle = {'height' : '100%', 'width' : '15vw'}
    if len(cfg.spliceSetNames[1]) == 0:
        rnaDataStyle = {'display': 'none', 'height' : '100%', 'width' : '15vw'}
    else:
        rnaDataStyle = {'height' : '100%', 'width' : '15vw'}
    
    
    # Try to setup color picker for iCLIP-seq tracks
    try:
        initialColor = cfg.dataSetNames[0]
        disableSettings = False
    except IndexError:
        initialColor = None
        disableSettings = False
    # Try to setup color picker for coverage tracks
    try:
        initialColorCoverage = cfg.spliceSetNames[1][0]
        disableSettings = False
    except IndexError:
        initialColorCoverage = None
        disableSettings = False
    # Try to setup color picker for coverage tracks
    try:
        initialColorEvents = cfg.eventTypes[0]
        disableSettings = False
    except IndexError:
        initialColorEvents = None
        disableSettings = False
        
    def help_popup():
        """ Defines the help popup that can be opened in the dashboard. Code is
        based on https://community.plot.ly/t/any-way-to-create-an-instructions-popout/18828/2
        by mbkupfer.
        """
        return html.Div(
            id='help',
            className="model",
            style={'display': 'none'},
            children=(
                html.Div(
                    className="help-container",
                    children=[
                        html.Div(
                            className='close-container',
                            children=html.Button(
                                "Close",
                                id="help_close",
                                n_clicks=0,
                                className="closeButton",
                                style={'border': 'none', 'height': '100%'}
                            )
                        ),
                        html.Div(
                            className='help-text',
                            children=[dcc.Markdown(
                                children= dedent(helpText)
                                )
                            ]
                        )
                    ]
                )
            )
    )
    
    # Default tab style
    tabStyle = {'padding' : '0', 'line-height' : '5vh'}
    # Colors for the alternating coloring in Details

    # For plots with multiple columns in the legend, basically anything using a color scale or heatmap,
    # this value determines the margin between colorbar and legend items. For very wide or very narrow
    # screens this value might need to be adjusted a bit, this can unfortunately not be done
    # automatically right now.
    legendColumnOffset = 1.05
    
    

app.config['suppress_callback_exceptions']=True
if __name__ == '__main__':
    if cfg.authentication != '': # Enable authentication if a password was provided
        auth = dash_auth.BasicAuth(
                app,
                {'u' : cfg.authentication})

    
    app.layout = html.Div(
        children=[
            html.Div(
                children = [
                    html.H1(id='headline', children='Report')
                ],
                style={'width': '90vw', 'display': 'table-cell', 'verticalalign': 'middle'}
            ),
            html.Div(
                children=[
                    html.Div(
                        children=[
                            html.Div(
                                children=[
                                    dcc.Dropdown(
                                        id='geneDrop',
                                        options=[{'label': i[0], 'value': i[1]} for i in cfg.dropList],
                                        value=cfg.dropList[0][1]
                                    )
                                ],
                                style={'width': '70vw', 'display': 'table-cell', 'verticalalign': 'middle'}
                            ),
                            html.Div(
                                style={'width': '1vw', 'display': 'table-cell', 'verticalalign': 'middle'}
                            ),
                            html.Div(
                                style={'width': '3vw', 'display': 'table-cell', 'verticalalign': 'middle'}
                            ),
                            html.Div(
                                children = [
                                    html.Button(id='helpButton', n_clicks=0, n_clicks_timestamp=0, children='help',
                                            style = {'backgroundColor' : 'rgb(255,255,255)'}
                                    )
                                ],
                                style={'width': '8vw', 'display': 'table-cell', 'verticalalign': 'middle'}
                            )
                        ]
                    ),
                    dcc.Tabs(
                        id='tabs',
                        style={
                            'width': '50%',
                            'height': '5vh'
                        },
                        children=[
                            dcc.Tab(
                                label='iCLIP-seq',
                                style=tabStyle,
                                selected_style=tabStyle,
                                id='clipTab',
                                children=[
                                    html.Div(children = [
                                        html.Div(className = 'table-cont',
                                            children=[
                                                    html.Div(className = 'table-row',
                                                        children = [
                                                        html.Details(open = 'open', children = [
                                                        html.Summary("Controls"),
                                                        html.Div(
                                                            className = 'table-cell column-1',
                                                            children = [
                                                                html.Fieldset(
                                                                    className = 'field-set',
                                                                    children = [ 
                                                                        html.Legend('Gene Description'),
                                                                        html.Div(id='descDiv',
                                                                            children = html.P(html.B(''))
                                                                        )
                                                                    ],
                                                                )
                                                            ]
                                                        ),
                                                        html.Div(style = dataSetStyle,
                                                            className = 'table-cell column-2',
                                                            children = [
                                                                html.Fieldset(
                                                                    className = 'field-set',
                                                                    children = [
                                                                        html.Legend('Datasets'),                                                     
                                                                        dcc.Checklist(
                                                                            id='paramList',
                                                                            options=[{'label': i, 'value': i} for i in cfg.dataSetNames],
                                                                            values=[i for i in cfg.dataSetNames]
                                                                        )
                                                                    ]
                                                                
                                                                )
                                                            ]
                                                        ),
                                                        html.Div(style = seqDispStyle,
                                                            className = 'table-cell column-3',
                                                            children = [
                                                                html.Fieldset(
                                                                    className = 'field-set',
                                                                    children = [
                                                                        html.Legend('DNA sequence options'),
                                                                        dcc.RadioItems(
                                                                            id='sequenceRadio',
                                                                            options=[
                                                                                {'label': 'Do not show dna sequence', 'value': 'noSeq'},
                                                                                {'label': 'Show dna sequence as letters', 'value': 'letterSeq'},
                                                                                {'label': 'Show dna sequence as heatmap', 'value': 'heatSeq'}
                                                                            ],
                                                                            value='heatSeq'
                                                                        )
                                                                    ]
                                                                )
                                                            ]
                                                        )
                                                    ]
                                                ),
                                                ])
                                            ],
                                        ),
                                        html.Div(style = {'height' : '25px'}),
                                        html.Div(
                                            children = [dcc.Store(id = 'bsGraphMem'),
                                                        dcc.Loading(
                                                        id ="iCLIP_loading",
                                                        type = 'dot',
                                                        children = [                                                               
                                                        dcc.Graph(id='bsGraph',
                                                            style = {'padding' : '3px'},
                                                            config = {'toImageButtonOptions' : 
                                                                {'filename' : 'iCLIP', 'width' : None,
                                                                'scale' : 1.0, 'height' : None, 'format' : 'svg'} }
                                                        )])                                              
                                            ]         
                                        )
                                    ]
                                )
                                ]
                            ),
                            dcc.Tab(
                                label='RNA-seq',
                                style=tabStyle,
                                selected_style=tabStyle,
                                id='rnaTab',
                                children=[
                                    html.Div(children=[
                                        html.Div(className='table-cont',
                                                 children=[
                                                     html.Div(className='table-row',
                                                              children=[
                                                              html.Details(open = 'open', children = [
                                                              html.Summary("Controls"),
                                                                  html.Div(
                                                                      className='table-cell column-1',
                                                                      children=[
                                                                          html.Fieldset(
                                                                              className='field-set',
                                                                              children=[
                                                                                  html.Legend('Gene Description'),
                                                                                  html.Div(id='rnaDescDiv',
                                                                                           children=html.P(html.B(''))
                                                                                           )
                                                                              ],
                                                                          )
                                                                      ]
                                                                  ),
                                                                  html.Div(style=rnaDataStyle,
                                                                           className='table-cell column-2',
                                                                           children=[
                                                                               html.Fieldset(
                                                                                   className='field-set',
                                                                                   children=[
                                                                                       html.Legend('Datasets'),
                                                                                       dcc.Checklist(
                                                                                           id='rnaParamList',
                                                                                           options=[{'label':
                                                                                                         cfg.spliceSetNames[1][
                                                                                                             i], 'value':
                                                                                                         cfg.spliceSetNames[1][
                                                                                                             i]} for
                                                                                                    i in range(
                                                                                                   len(cfg.spliceSetNames[1]))],
                                                                                           values=[cfg.spliceSetNames[1][i] for
                                                                                                   i in range(
                                                                                                   len(cfg.spliceSetNames[1]))],
                                                                                       ),
                                                                                   ]
    
                                                                               )
                                                                           ]
                                                                           ),
                                                                  html.Div(style={'height': '100%', 'width': '20vw'},
                                                                           className='table-cell column-3',
                                                                           children=[
                                                                               html.Fieldset(
                                                                                   className='field-set',
                                                                                   children=[
                                                                                       html.Legend('Options'),
                                                                                       dcc.RadioItems(
                                                                                           id='rnaRadio',
                                                                                           options=[
                                                                                               {'label': 'Default display',
                                                                                                'value': 'one'},
                                                                                               {'label': 'Color event types',
                                                                                                'value': 'two'},
                                                                                               {'label': 'Color event scores',
                                                                                                'value': 'three'}
                                                                                           ],
                                                                                           value='one'
                                                                                       )
                                                                                   ]
                                                                               )
                                                                           ]
                                                                           )
                                                              ]
                                                              ),
                                                              ])
                                                            
                                                        ],
    
                                    ),
                                    html.Div(style = {'height' : '25px'}),
                                    dcc.Loading(
                                        id ="RNAseq_loading",
                                        type = 'circle',
                                        children = [
                                            dcc.Store(id = 'spliceMem'),
                                            dcc.Graph(id='spliceGraph',
                                            style = {'padding' : '3px'},
                                            config = {'toImageButtonOptions' : 
                                            {'filename' : 'iCLIP', 'width' : None, 'scale' : 1.0, 'height' : None, 'format' : 'svg'} }
                                            )
                                        ]
                                    )
                                ])
                            ]
                            ),
                            dcc.Tab(
                                label = 'Details',
                                id = 'deTab',
                                disabled = advDisabled,
                                style = tabStyle,
                                selected_style = tabStyle,
                                disabled_style = tabStyle,
                                children = [
                                    html.Div(
                                        id='detailMainDiv',
                                        children=[]
                                    )
                                ]
                            ),
                            dcc.Tab(
                                label='Settings',
                                id='settings',
                                style=tabStyle,
                                selected_style=tabStyle,
                                disabled=disableSettings,
                                disabled_style=tabStyle,
                                children=[
                                    html.Div(style = {'display' : 'table'},
                                    children =[
                                    html.Div(style={'width': '100vw', 'display': 'table-row', 'verticalalign': 'middle'},
                                        children=[
                                            html.Div(className = 'table-cell', 
                                            children = [
                                            html.Fieldset(title = 'iCLIP Settings', 
                                                className = 'field-set',
                                                children = [ 
                                                    html.Legend('iCLIP Settings'),
                                                    dcc.Store(
                                                        id='colorDiv',
                                                        data=cfg.colorMap
                                                    ),
                                                    dcc.Store(
                                                        id='colorFinal',
                                                        data=cfg.colorMap
                                                    ),
                                                    dcc.Dropdown(
                                                        id='colorDrop',
                                                        options=[{'label': i, 'value': i} for i in cfg.dataSetNames],
                                                        value=initialColor
                                                    ),
                                                    html.Div(
                                                        id='rDisp',
                                                        children=html.P(html.B('R: ')),
                                                        style={'width': '10vw', 'display': 'table-cell'}
                                                    ),
                                                    dcc.Slider(
                                                        id='rInput',
                                                        min=0,
                                                        max=255,
                                                        step=1,
                                                        updatemode='drag'
                                                    ),
                                                    html.Div(
                                                        id='gDisp',
                                                        children=html.P(html.B('G: ')),
                                                        style={'width': '10vw', 'display': 'table-cell'}
                                                    ),
                                                    dcc.Slider(
                                                        id='gInput',
                                                        min=0,
                                                        max=255,
                                                        step=1,
                                                        updatemode='drag'
                                                    ),
                                                    html.Div(
                                                        id='bDisp',
                                                        children=html.P(html.B('B: ')),
                                                        style={'width': '10vw', 'display': 'table-cell'}
                                                    ),
                                                    dcc.Slider(
                                                        id='bInput',
                                                        min=0,
                                                        max=255,
                                                        step=1,
                                                        updatemode='drag'
                                                    ),
                                                    html.Div(
                                                        children=[
                                                            html.Div(id='preview',
                                                                     children=html.P(html.B('Preview')),
                                                                     style={'width': '30vw', 'display': 'table-cell',
                                                                            'verticalalign': 'middle'}
                                                                     ),
                                                            html.Div(
                                                                children=[
                                                                    html.Button(id='colorConfirm', n_clicks_timestamp=0,
                                                                                children='confirm')
                                                                ],
                                                                style={'width': '10vw', 'display': 'table-cell',
                                                                       'verticalalign': 'middle'}
                                                            )
                                                        ],
                                                        style={'width': '10vw', 'display': 'table-cell'}
                                                    )
                                                ]
                                            )]),
                                            html.Div(className = 'table-cell', 
                                            children = [
                                            html.Fieldset(title = 'coverage settings', 
                                                className = 'field-set',
                                                children = [ 
                                                    html.Legend('Coverage plot settings'),
                                                    dcc.Store(
                                                        id='covColorDiv',
                                                        data=cfg.coverageColors
                                                    ),
                                                    dcc.Store(
                                                        id='covColorFinal',
                                                        data=cfg.coverageColors
                                                    ),
                                                    dcc.Dropdown(
                                                        id='covColorDrop',
                                                        options=[{'label': i, 'value': i} for i in cfg.spliceSetNames[1]],
                                                        value=initialColorCoverage
                                                    ),
                                                    html.Div(
                                                        id='covRDisp',
                                                        children=html.P(html.B('R: ')),
                                                        style={'width': '10vw', 'display': 'table-cell'}
                                                    ),
                                                    dcc.Slider(
                                                        id='covRInput',
                                                        min=0,
                                                        max=255,
                                                        step=1,
                                                        updatemode='drag'
                                                    ),
                                                    html.Div(
                                                        id='covGDisp',
                                                        children=html.P(html.B('G: ')),
                                                        style={'width': '10vw', 'display': 'table-cell'}
                                                    ),
                                                    dcc.Slider(
                                                        id='covGInput',
                                                        min=0,
                                                        max=255,
                                                        step=1,
                                                        updatemode='drag'
                                                    ),
                                                    html.Div(
                                                        id='covBDisp',
                                                        children=html.P(html.B('B: ')),
                                                        style={'width': '10vw', 'display': 'table-cell'}
                                                    ),
                                                    dcc.Slider(
                                                        id='covBInput',
                                                        min=0,
                                                        max=255,
                                                        step=1,
                                                        updatemode='drag'
                                                    ),
                                                    html.Div(
                                                        children=[
                                                            html.Div(id='covPreview',
                                                                     children=html.P(html.B('Preview')),
                                                                     style={'width': '30vw', 'display': 'table-cell',
                                                                            'verticalalign': 'middle'}
                                                                     ),
                                                            html.Div(
                                                                children=[
                                                                    html.Button(id='covColorConfirm', n_clicks_timestamp=0,
                                                                                children='confirm')
                                                                ],
                                                                style={'width': '10vw', 'display': 'table-cell',
                                                                       'verticalalign': 'middle'}
                                                            )
                                                        ],
                                                        style={'width': '10vw', 'display': 'table-cell'}
                                                    )
                                                ]
                                            )]),
                                            html.Div(className = 'table-cell', 
                                            children = [
                                            html.Fieldset(title = 'event settings', 
                                                className = 'field-set',
                                                children = [ 
                                                    html.Legend('Splice event plot settings'),
                                                    dcc.Store(
                                                        id='eventColorDiv',
                                                        data=cfg.eventColors
                                                    ),
                                                    dcc.Store(
                                                        id='eventColorFinal',
                                                        data=cfg.eventColors
                                                    ),
                                                    dcc.Dropdown(
                                                        id='eventColorDrop',
                                                        options=[{'label': i, 'value': i} for i in cfg.eventTypes],
                                                        value=initialColorEvents
                                                    ),
                                                    html.Div(
                                                        id='eventRDisp',
                                                        children=html.P(html.B('R: ')),
                                                        style={'width': '10vw', 'display': 'table-cell'}
                                                    ),
                                                    dcc.Slider(
                                                        id='eventRInput',
                                                        min=0,
                                                        max=255,
                                                        step=1,
                                                        updatemode='drag'
                                                    ),
                                                    html.Div(
                                                        id='eventGDisp',
                                                        children=html.P(html.B('G: ')),
                                                        style={'width': '10vw', 'display': 'table-cell'}
                                                    ),
                                                    dcc.Slider(
                                                        id='eventGInput',
                                                        min=0,
                                                        max=255,
                                                        step=1,
                                                        updatemode='drag'
                                                    ),
                                                    html.Div(
                                                        id='eventBDisp',
                                                        children=html.P(html.B('B: ')),
                                                        style={'width': '10vw', 'display': 'table-cell'}
                                                    ),
                                                    dcc.Slider(
                                                        id='eventBInput',
                                                        min=0,
                                                        max=255,
                                                        step=1,
                                                        updatemode='drag'
                                                    ),
                                                    html.Div(
                                                        children=[
                                                            html.Div(id='eventPreview',
                                                                     children=html.P(html.B('Preview')),
                                                                     style={'width': '30vw', 'display': 'table-cell',
                                                                            'verticalalign': 'middle'}
                                                                     ),
                                                            html.Div(
                                                                children=[
                                                                    html.Button(id='eventColorConfirm', n_clicks_timestamp=0,
                                                                                children='confirm')
                                                                ],
                                                                style={'width': '10vw', 'display': 'table-cell',
                                                                       'verticalalign': 'middle'}
                                                            )
                                                        ],
                                                        style={'width': '10vw', 'display': 'table-cell'}
                                                    )
                                                ]
                                            )]),
 html.Div(className = 'table-cell', 
                                                children = [
                                                    html.Fieldset(title = 'iCLIP-seq plot settings', 
                                                        className = 'field-set',
                                                        children = [
                                                            html.Legend('iCLIP-seq plot settings'),
                                                            html.Div(children = 'iCLIP plot scale'),
                                                            dcc.Slider(
                                                                id = 'iCLIPScale',
                                                                min = 0.25,
                                                                max = 2.5,
                                                                step = 0.05,
                                                                value = 1.0,
                                                                marks = {
                                                                    0.25: '0.25',
                                                                    1.0: '1.0',
                                                                    1.5: '1.5',
                                                                    2.0: '2.0',
                                                                    2.5: '2.5',
                                                                    }
                                                            ),
                                                            html.Div(
                                                                style = {'height' : '25px'}),
                                                            html.Div(children = 'Binding site plot scale'),
                                                            dcc.Slider(
                                                                id = 'bsScale',
                                                                min = 0.25,
                                                                max = 2.5,
                                                                step = 0.05,
                                                                value = 1.0,
                                                                marks = {
                                                                    0.25: '0.25',
                                                                    1.0: '1.0',
                                                                    1.5: '1.5',
                                                                    2.0: '2.0',
                                                                    2.5: '2.5',
                                                                    }
                                                            ),
                                                            html.Div(
                                                                style = {'height' : '15px'})
                                                        ]
                                                    )
                                                ]
                                            ),                                                                
                                            html.Div(className = 'table-cell', 
                                                children = [
                                                    html.Fieldset(title = 'RNA-seq plot settings', 
                                                        className = 'field-set',
                                                        children = [
                                                            html.Legend('RNA-seq plot settings'),
                                                            html.Div(children = 'Coverage plot scale'),
                                                            dcc.Slider(
                                                                id = 'coverageScale',
                                                                min = 0.25,
                                                                max = 2.5,
                                                                step = 0.05,
                                                                value = 0.5,
                                                                marks = {
                                                                    0.25: '0.25',
                                                                    1.0: '1.0',
                                                                    1.5: '1.5',
                                                                    2.0: '2.0',
                                                                    2.5: '2.5',
                                                                    }
                                                            ),
                                                            html.Div(
                                                                style = {'height' : '25px'}),
                                                            html.Div(children = 'Event plot scale'),
                                                            dcc.Slider(
                                                                id = 'eventScale',
                                                                min = 0.25,
                                                                max = 2.5,
                                                                step = 0.05,
                                                                value = 0.5,
                                                                marks = {
                                                                    0.25: '0.25',
                                                                    1.0: '1.0',
                                                                    1.5: '1.5',
                                                                    2.0: '2.0',
                                                                    2.5: '2.5',
                                                                    }
                                                            ),
                                                            html.Div(
                                                                style = {'height' : '15px'})
                                                        ]
                                                    )
                                                ]
                                            ),
                                            html.Div(className = 'table-cell', 
                                                children = [
                                                    html.Fieldset(title = 'Legend Settings', 
                                                        className = 'field-set',
                                                        children = [
                                                            html.Legend('Legend Colorbar Margin'),
                                                            dcc.Store(
                                                                id = 'legendSpacingDiv',
                                                                data = legendColumnOffset
                                                                ),
                                                            dcc.Slider(
                                                                id = 'legendSpacingSlider',
                                                                min = 1.02,
                                                                max = 1.08,
                                                                step = 0.005,
                                                                value = legendColumnOffset,
                                                                marks = {
                                                                    1.02: '.02',
                                                                    1.03: '',
                                                                    1.04: '',
                                                                    1.05: '.05',
                                                                    1.06: '',
                                                                    1.07: '',
                                                                    1.08: '.08',
                                                                    }
                                                            ),
                                                            html.Div(
                                                                style = {'height' : '15px'})
                                                        ]
                                                    )
                                                ]
                                            ),
                                            html.Div(className = 'table-cell', 
                                                children = [
                                                    html.Fieldset(title = 'Image Export', 
                                                        className = 'field-set',
                                                        children = [
                                                            html.Legend('Image export format'),
                                                            dcc.Dropdown(
                                                                id = 'imgFormatDrop',
                                                                options = [{'label': i, 'value': i} for i in ['png', 'svg']],
                                                                value = imgFormat
                                                            ),
                                                            html.Div(
                                                                style = {'height' : '15px'})
                                                        ]
                                                    )
                                                ]
                                            )
                                        ],
                                    ),
                                ])
                                ]
                            )
                        ]
                    )
    
                ]
            ),
            help_popup()
        ],
        style = {'backgroundColor' : 'rgb(240,240,240)'}
    )

@app.callback(
    dash.dependencies.Output('help', 'style'),
    [dash.dependencies.Input("helpButton", "n_clicks"),
     dash.dependencies.Input("help_close", "n_clicks")]
)
def showHelpPopup(open_click, close_click):
    """ Display and hide the help popup depending on the clicked button. Code is
    based on https://community.plot.ly/t/any-way-to-create-an-instructions-popout/18828/3
    by mbkupfer
    
    Positional arguments:
    open_click -- Open the popup
    close_click -- Close the popup
    """
    if open_click > close_click:
        return {"display": "block"}
    else:
        return {"display": "none"}

if __name__ == '__main__':
    app.run_server(debug=True, host='0.0.0.0', port=cfg.port, use_reloader=False)
```
