## Additional File 1 for "SEQing: web-based visualization of iCLIP and RNA-seq data in an interactive python framework": cfg.py

```
#!/usr/bin/env python3
# -*- coding: utf-8 -*-

def init(globs):
    """ Takes globals from validator and stores them in variables for the other modules.
    
    Positional arguments:
    globs -- Globals from validator.
    """
    global colorMap
    colorMap = globs['colorMap'] 
    global descAvail
    descAvail = globs['descAvail']
    global colorA, colorC, colorG, colorT
    colorA = globs['colorA']
    colorC = globs['colorC']
    colorG = globs['colorG']
    colorT = globs['colorT']
    global port
    port = globs['port']
    global procAvail
    procAvail = globs['procAvail']
    global dsElements
    dsElements = globs['dsElements'] 
    global spliceElements
    spliceElements = globs['spliceElements']
    global bsProcDFs
    bsProcDFs = globs['bsProcDFs']
    global bsRawDFs
    bsRawDFs = globs['bsRawDFs'] 
    global dataSetNames
    dataSetNames = globs['dataSetNames']
    global spliceSetNames
    spliceSetNames = globs['spliceSetNames'] 
    global rawAvail
    rawAvail = globs['rawAvail']
    global spliceAvail
    spliceAvail = globs['spliceAvail']
    global dropList
    dropList = globs['dropList']
    global geneDescriptions
    geneDescriptions = globs['geneDescriptions']
    global sequences
    sequences = globs['sequences']
    global geneAnnotations
    geneAnnotations = globs['geneAnnotations']
    global sortKeys
    sortKeys = globs['sortKeys']
    global advancedDesc
    advancedDesc = globs['advancedDesc']
    global subTables
    subTables = globs['subTables']
    global spliceEventElements
    spliceEventElements = globs['spliceEventElements']
    global spliceEventDFs
    spliceEventDFs = globs['spliceEventDFs']
    global spliceEventNames
    spliceEventNames = globs['spliceEventNames']
    global spliceEventAvail
    spliceEventAvail= globs['spliceEventAvail']
    global eventColors
    eventColors = globs['eventColors']
    global coverageColors
    coverageColors = globs['coverageColors']
    global eventTypes
    eventTypes = globs['eventTypes']
    global authentication
    authentication = globs['authentication']
    global coverageData
    coverageData = globs['coverageData']
```
