## Additional File 1 for "SEQing: web-based visualization of iCLIP and RNA-seq data in an interactive python framework": converter.py

```
#!/usr/bin/env python
# -*- coding: utf-8 -*-

import pandas as pd
import filetype as ft
import numpy as np

__author__ = "Yannik Bramkamp"

if __name__ == '__main__':
    print('Please start the program via validator.py')
    exit()

gtfHeader = ['chrom','chromStart','chromEnd','geneID', 'transID','score','strand','thickStart','thickEnd','itemRGB','blockCount','blockSizes','blockStarts']
dtypes = {'chrom' : 'category', 'chromStart' : 'uint32','chromEnd': 'uint32','geneID' : 'object', 'transID' : 'object','score' : 'int16','strand' : 'category','thickStart' : 'uint64',
                                  'thickEnd' : 'uint64', 'itemRGB' : 'int16', 'blockCount' : 'uint32','blockSizes' : 'object','blockStarts' : 'object'}

def convertGTFToBed(df):
    """ Convert a GTF dataframe to a BED12 dataframe for internal use. Only relevant
    lines will be taken into account, in this case exon and cds can be used 
    to reconstruct all needed information.
    
    Positional arguments:
    df -- GTF dataframe.
    """
    bedFile = []
    current = ''
    currentGene = ''
    chrom = ''
    strand = ''
    itemRGB = 0
    score = 0
    blockCount = 0
    blockStarts = []
    blockEnds = []
    blockSizes = []
    thickStart = -1
    thickEnd = -1
    
    rowIndex = 1
    percent = 10
    for i in df.iterrows():
        if rowIndex/len(df)*100 >= percent:
            print(str(percent) + '%')
            percent += 10
        rowIndex +=1
        # Split attributes for relevant lines and extract needed ones
        if i[1]['feature'] == 'exon' or i[1]['feature'] == 'CDS':
            attributes = i[1]['attribute'].split(';')
            geneID = [s for s in attributes if 'gene_id' in s]
            transID = [s for s in attributes if 'transcript_id' in s]
            # check if gene and transcript id are specified only once
            if len(geneID) == 1:
                geneID = geneID[0].split(' ')[-1]
                if geneID[0] == '"':
                    geneID = geneID[1:-1]
            else:
                geneID = ''
            if len(transID) == 1:
                transID = transID[0].split(' ')[-1]
                if transID[0] == '"':
                    transID = transID[1:-1]
            else:
                transID = ''
#            # handle ids that consist of multiple, point seperated values
#            geneParts = geneID.split('.')
#            transParts = transID.split('.')
#            finalID = '' # this will be the final id used for the gene
#            if geneID != '':
#                if transID != '':
#                    if len(geneParts) == 1 and len(transParts) == 1:
#                        finalID = geneParts[0] + '.' + transParts[0]
#                    else:
#                        if len(geneParts) == 1 and len(transParts) == 2:
#                            if geneParts[0] == transParts[0]:
#                                finalID = geneParts[0] + '.' + transParts[1]
#                            else:
#                                finalID = geneParts[0] + '.' + '_'.join(transParts)
#                        else:
#                            finalID = '_'.join(geneParts) + '.' + '_'.join(transParts)
#                else:
#                    finalID = geneID
#            else:
#                continue
            # convert the data into bed 12
            if current == '':
                current = transID
                currentGene = geneID
                chrom = i[1]['seqname']
                strand = i[1]['strand']
                blockCount = 0
                blockStarts = []
                blockEnds = []
                blockSizes = []
                thickStart = -1
                thickEnd = -1
                if i[1]['feature'] == 'exon':
                    blockCount += 1
                    blockStarts.append(int(i[1]['start']))
                    blockEnds.append(int(i[1]['end']))
                    blockSizes.append(int(i[1]['end'])-int(i[1]['start'])+1)
                if i[1]['feature'] == 'CDS':
                    if thickStart == -1 or thickStart > i[1]['start']:
                        thickStart = int(i[1]['start'])-1
                    if thickEnd == -1 or thickEnd < i[1]['end']:
                        thickEnd = i[1]['end']
            else:
                if transID == current: # line still belongs to the same gene
                    if i[1]['feature'] == 'exon':
                        blockCount += 1
                        blockStarts.append(int(i[1]['start']))
                        blockEnds.append(int(i[1]['end']))
                        blockSizes.append(int(i[1]['end'])-int(i[1]['start'])+1)
                    if i[1]['feature'] == 'CDS':
                        if thickStart == -1 or thickStart > i[1]['start']:
                            thickStart = int(i[1]['start'])-1
                        if thickEnd == -1 or thickEnd < i[1]['end']:
                            thickEnd = i[1]['end']
                else:   # line belongs to a new gene with a different final id
                    chromStart = min(blockStarts)
                    chromEnd = max(blockEnds)
                    if thickEnd == -1:
                        thickEnd = chromEnd
                    if thickStart == -1:
                        thickStart = chromStart-1
                    bBlockStarts = [int(i)-int(chromStart) for i in blockStarts]
                    # append data of previous gene before resetting values for
                    # the new one
                    bedFile.append([chrom, chromStart-1, chromEnd, currentGene, current, score,
                                    strand, thickStart, thickEnd, itemRGB,
                                    blockCount, ','.join(map(str, blockSizes)), ','.join(map(str, bBlockStarts))])
                    current = transID
                    currentGene = geneID
                    chrom = i[1]['seqname']
                    strand = i[1]['strand']
                    blockCount = 0
                    blockStarts = []
                    blockEnds = []
                    blockSizes = []
                    thickStart = -1
                    thickEnd = -1    
                    if i[1]['feature'] == 'exon':
                        blockCount += 1
                        blockStarts.append(int(i[1]['start']))
                        blockEnds.append(int(i[1]['end']))
                        blockSizes.append(int(i[1]['end'])-int(i[1]['start'])+1)
                    if i[1]['feature'] == 'CDS':
                        if thickStart == -1 or thickStart > i[1]['start']:
                            thickStart = int(i[1]['start'])-1
                        if thickEnd == -1 or thickEnd < i[1]['end']:
                            thickEnd = i[1]['end']
    chromStart = min(blockStarts)
    chromEnd = max(blockEnds)
    if thickEnd == -1:
        thickEnd = chromEnd
    if thickStart == -1:
        thickStart = chromStart-1
    bBlockStarts = [int(i)-int(chromStart) for i in blockStarts]
    # append data of previous gene before resetting values for
    # the new one
    bedFile.append([chrom, chromStart-1, chromEnd, geneID, transID, score,
                                    strand, thickStart, thickEnd, itemRGB,
                                    blockCount, ','.join(map(str, blockSizes)), ','.join(map(str, bBlockStarts))])
    finDF = pd.DataFrame(data = bedFile, columns = gtfHeader)
    for key, dtype in dtypes.items():
        finDF[key] = finDF[key].astype(dtype)
    return finDF


class FileInput:
    """Decorator for file checking"""
    def __init__(self, file_path, file_type, zipped, zip_type, header_present):
        self.file_path = file_path
        self.file_type = file_type
        self.zipped = zipped
        self.zip_type = zip_type
        self.header_present = header_present


def check_input_file(file_path):
    """A function to test an input file and classifies it by content"""
    # try to guess file type by analysing the head
    guessed_type = ft.guess(file_path)
    file_zipped = False

    if guessed_type == None:
        # read uncompressed head
        file_head = pd.read_csv(file_path, sep='\t', header=None, nrows=5, error_bad_lines = False)
        zip_type = None
    elif guessed_type.mime in ['application/gzip', 'application/x-bzip2', 'application/zip']:
        zip_type = guessed_type.mime.split('/')[1]
        # read compressed head
        file_head = pd.read_csv(file_path, compression='infer', sep='\t', header=None, nrows=5, error_bad_lines = False, warn_bad_lines = False)
        file_zipped = True
    else:
        # return unsupported file type
        return FileInput(file_path, 'unsupported', False, False)
        zip_type == None
    head_dtypes = np.array(file_head.dtypes)
    # check for header (no numbers in first row)
    header_present = not any(cell == np.int for cell in head_dtypes)
    header = pd.Series()
    if header_present:
        header = file_head.iloc[0]
        if file_zipped:
            file_head = pd.read_csv(file_path, compression='infer', sep='\t', header=None, nrows=5, skiprows=1, comment='#')
        else:
            file_head = pd.read_csv(file_path, sep='\t', header=None, nrows=5, skiprows=1, comment='#')

    # assign file type by shape of table
    head_dim = file_head.shape
    # check for BED4
    if head_dim[1] == 4:
        return FileInput(file_path, 'BED4', file_zipped, zip_type, header_present)
    # check for BED6
    elif head_dim[1] == 6:
        return FileInput(file_path, 'BED4', file_zipped, zip_type, header_present)
    # check for GFF or GTF
    elif head_dim[1] == 9:
        if not header.empty:
            for col in header:
                if 'gff-version 3' in col:
                    return FileInput(file_path, 'GFF3', file_zipped, zip_type, header_present)
                else:
                    return FileInput(file_path, 'GTF', file_zipped, zip_type, header_present)
        else:
            if '"' in file_head.iloc[0, 8]:
                return FileInput(file_path, 'GTF', file_zipped, zip_type, header_present)
    elif head_dim[1] == 12:
        return FileInput(file_path, 'BED12', file_zipped, zip_type, header_present)
    else:
        # unsupported format
        return FileInput(file_path, 'unsupported', False, zip_type, header_present)
```
