## Additional File 1 for "SEQing: web-based visualization of iCLIP and RNA-seq data in an interactive python framework": description_tab.py

```
#!/usr/bin/env python3
# -*- coding: utf-8 -*-

import dash
import pandas
import dash_html_components as html
from app import app
import cfg

tableColors = ['rgb(255, 255 ,255)', 'rgb(220, 220, 220)']

@app.callback(
    dash.dependencies.Output('detailMainDiv', component_property = 'children'),
    [dash.dependencies.Input('geneDrop', 'value')]
)
def showDetails(name):
    """ Create tabular view of additional data

    Positional arguments:
    name -- Gene name for initialization.
    """
    if cfg.advancedDesc is not None:
        try:
            df = cfg.advancedDesc[cfg.advancedDesc['gene_ids'].str.contains(name)]
        except TypeError:
            df = pandas.DataFrame()
    else:
        df = pandas.DataFrame()
    columns = list(df.columns.values)
    rowCounter = 1  # Keep track of the row number to alternate coloring
    usedColumns = []  # Keeps track of preset columns already added, needed later
    usedColumns.append('gene_ids')
    content = []  # Table content to be displayed
    generalColumns = ['symbol', 'brief_description', 'is_obsolete', 'computational_description',
                      'curator_summary', 'name']
    tableRows = []  # Will contain the table rows
    for i in generalColumns:
        if i in columns:
            if str(df.iloc[0][i]) not in ['nan', ';""']:
                tableRows.append(html.Tr(children=[html.Td(html.B(i.replace('_', ' ').title())),
                                               html.Td(str(df.iloc[0][i]).strip())],
                                     style={'background-color': tableColors[rowCounter % 2]}))
                usedColumns.append(i)
                rowCounter += 1
    # Go through a number of predefined columns
    if 'synonyms' in columns:
        synonyms = str(df.iloc[0]['synonyms'])
        usedColumns.append('synonyms')
        if synonyms not in ['nan', ';""']:
            tableRows.append(createDetailRow(synonyms, 'synonyms', rowCounter))
            rowCounter += 1
    if 'publications' in columns:
        usedColumns.append('publications')
        publications = str(df.iloc[0]['publications'])
        if publications not in ['nan', ';""']:
            tableRows.append(createDetailRow(publications, 'publications', rowCounter))
            rowCounter += 1
    if 'proteins' in columns:
        usedColumns.append('proteins')
        proteins = str(df.iloc[0]['proteins'])
        if publications not in ['nan', ';""']:
            tableRows.append(createDetailRow(proteins, 'proteins', rowCounter))
            rowCounter += 1
    if 'gene_ontology' in columns:
        usedColumns.append('gene_ontology')
        geneOntology = str(df.iloc[0]['gene_ontology'])
        if geneOntology not in ['nan', ';""']:
            tableRows.append(createDetailRow(geneOntology, 'gene_ontology', rowCounter))
            rowCounter += 1
    if 'pathways' in columns:
        usedColumns.append('pathways')
        pathways = str(df.iloc[0]['pathways'])
        if pathways not in ['nan', ';""']:
            tableRows.append(createDetailRow(pathways, 'pathways', rowCounter))
            rowCounter += 1
    if 'plant_ontology' in columns:
        usedColumns.append('plant_ontology')
        plantOntology = str(df.iloc[0]['plant_ontology'])
        if plantOntology not in ['nan', ';""']:
            tableRows.append(createDetailRow(plantOntology, 'plant_ontology', rowCounter))
            rowCounter += 1
    # Go through all remaining columns using formatting standard
    remainingColumns = [x for x in columns if x not in usedColumns]
    for i in remainingColumns:
        value = str(df.iloc[0][i])
        if value not in ['nan', ';""']:
            tableRows.append(createDetailRow(value, i, rowCounter))
            rowCounter += 1

    if len(tableRows) >= 1:
        content.append(html.Table(tableRows))
    else:
        content.append(html.B('No additional information available for the currently selected gene'))
    return content


def createDetailRow(content, name, rowNumber):
    """ Returns a single row for the details table

    Positional arguments:
    content -- The attribute data as String.
    name -- Name for the attribute.
    rowNumber -- Used for odd/even coloring.
    """
    # Check subtable information
    try:
        headerLine = [cfg.subTables['column_id'].str.contains(name)]
    except (TypeError, AttributeError, KeyError):
        headerLine = None
    try:
        headers = str(headerLine.iloc[0]['columns']).split(';')
    except (TypeError, AttributeError, KeyError):
        headers = None
    if content == None or name == None:
        return None
    subRows = [] # Holds elements for multivalue attributes
    subTable = [] # Holds elements for subtables
    if headers != None: # We have subtable information, so try and create one
        headerRow = []
        for k in headers: # Build table header line
            headerRow.append(html.Th(k))
        subTable.append(html.Tr(children = headerRow))
        tableError = False
        for i in content.split(';'): # Build subtable rows dictated by ; delimitation
            subSubRow = []
            if len(i.split(',')) == len(headers):
                for j in i.split(','): # Build subtable columns dictated by , delimitation
                    if j != '':
                        if j[0] == '?':
                            subSubRow.append(html.Td(html.A(j[1:], href=j[1:].strip(), target='_blank')))
                        else:
                            subSubRow.append(html.Td(j.strip()))    
                subTable.append(html.Tr(children = subSubRow))
            else: 
                tableError = True
        if tableError == True: # Column numbers didn't match, default display
            print('Warning: Number of columns specified in subtable file do not match number of columns in description file')
            subTable = []
            for l in content.split(';'):
                if l != '' :
                    if l[0] == '?': # Create hyperlinks
                        subRows.append(html.Tr(html.Td(html.A(l[1:], href = l[1:].strip(), target = '_blank'))))
                    else:
                        subRows.append(html.Tr(html.Td(l.strip())))
    else: # No subtable information
        for i in content.split(';'):
            if i != '' :
                    if i[0] == '?':
                        subRows.append(html.Tr(html.Td(html.A(i[1:], href = i[1:].strip(), target = '_blank'))))
                    else:
                        subRows.append(html.Tr(html.Td(i.strip())))
    if len(subRows) > 5: # Hide values in details element if more than 5 values
        tableRow = html.Tr(children = [html.Td(html.B(name.replace('_',' ').title())),
                               html.Td(html.Details(title = str(len(subRows)) + ' values', children = [html.Summary(str(len(subRows)) + ' values'), html.Table(children = 
                                       subRows)]))], style = {'background-color' : tableColors[rowNumber%2]})
    else:
        tableRow = html.Tr(children=[html.Td(html.B(name.replace('_', ' ').title())),
                                     html.Td(html.Table(children=
                                                        subRows))],
                           style={'background-color': tableColors[rowNumber % 2]})
    if len(subTable) > 0:
        return html.Tr(
            children=[html.Td(html.B(name.replace('_', ' ').title())), html.Td(html.Table(children=subTable))],
            style={'background-color': tableColors[rowNumber % 2]})
    else:
        return tableRow
```
