## Additional File 1 for "SEQing: web-based visualization of iCLIP and RNA-seq data in an interactive python framework": start_sample.sh

```
#!/bin/bash

if command -v python3 &>/dev/null; then
	echo Python3 is installed, starting dashboard ...
	mkdir -p ../bin_data
	python3 ../validator.py Araport_sample.bed -seq Araport_sample.fa -bsraw AtGRP7-LL24_crosslinks.bedgraph AtGRP7-LL36_crosslinks.bedgraph -bsdata AtGRP7-LL24_significant.bed AtGRP7-LL36_significant.bed -desc descriptions_sample.tsv -port 8066 -adv_desc advanced_details_sample.tsv -splice_data Col2_LL24.bedgraph Col2_LL36.bedgraph grp7-1\ 8i_LL24.bedgraph grp7-1\ 8i_LL36.bedgraph AtGRP7-ox_LL24.bedgraph AtGRP7-ox_LL36.bedgraph -splice_events grp7-1\ 8i_LL24.dpsi.bed grp7-1\ 8i_LL36.dpsi.bed AtGRP7-ox_LL24.dpsi.bed AtGRP7-ox_LL36.dpsi.bed.gz -name sample -colors 'rgb(88, 151, 255)' 'rgb(0, 0, 255)' -geneindex gene_index.csv
else
	echo Python3 is not installed, quitting
fi
```
