## Additional File 1 for "SEQing: web-based visualization of iCLIP and RNA-seq data in an interactive python framework": iclip_tab.py

```
#!/usr/bin/env python3
# -*- coding: utf-8 -*-

import dash
import pandas
from app import app
import cfg
import dash_html_components as html
from plotly import tools
import plotly.graph_objs as go
import plotly.utils as pu

@app.callback(
    dash.dependencies.Output('descDiv', component_property='children'),
    [dash.dependencies.Input('geneDrop', 'value')]
)
def setDesc(name):
    """Callback to update gene description.

    Positional arguments:
    name -- Name of the currently selected gene
    """
    if cfg.descAvail:
        try:
            return [
                html.P(    
                    cfg.geneDescriptions.loc[
                        cfg.geneDescriptions['ensembl_gene_id'] == name,
                        ['description']
                    ].iloc[0]
                )
            ]
        except IndexError:
            return ['No description available']
        except KeyError:
            return ['No description available']        
    else:
        return ['No description available']

@app.callback(
    dash.dependencies.Output('bsGraph', 'figure'),
    [dash.dependencies.Input('bsGraphMem', 'data'),
     dash.dependencies.Input('paramList', 'values'),
     dash.dependencies.Input('sequenceRadio', 'value'),
     dash.dependencies.Input('colorFinal', 'data'),
     dash.dependencies.Input('legendSpacingDiv', 'data'),
     dash.dependencies.Input('iCLIPScale', 'value'),
     dash.dependencies.Input('bsScale', 'value')]
)
def showICLIP(figData, dataSets, seqDisp, colorF, legendSpacing,  iCLIPScale, bsScale):
    """ Update callbacks that selects traces to be displayed based on user input.
    
    Positional arguments:
    figData -- Trace data from the data callback.
    datasets -- List of selected datasets.
    seqDisp -- Sytle for the reference sequence.
    colorF -- Colors for the traces.
    legendSpacing -- Spacing between legend and colorbar.
    """
    figData = figData
    traces = []
    rowHeights = []
    legendColumnSpacing = legendSpacing
    numRows = 1
    try:
        seqTrace = figData[seqDisp]
    except:
        seqTrace = []
    colorDict = colorF
    numIsoforms = len(figData['geneModels'])
    numParams = 0
    for index, elem in enumerate(figData['iCLIPTraces']):
            name = elem[0]['meta']
            newColor = colorDict[name]
            if name in dataSets:
                numParams += 1
                element = figData['iCLIPTraces'][index][0]
                element['marker'] = go.bar.Marker(
                        color = newColor 
                        )
                traces.append(element)
                
                numRows += 1
                if len(elem) == 2:
                    if elem[1] != [] or cfg.procAvail:
                        elements = elem[1]
                        for i in elements:
                            i['marker'] = {'color' : newColor}
                        traces.append(elements)
                        numRows += 1
    for i in figData['geneModels']:
        traces.append(i)
        numRows += 1
    plotSpace = 0.8  # Space taken up by data tracks
    spacingSpace = 1.0 - plotSpace  # Space left for spacer tracks
    rowHeight = plotSpace / numRows
    if numRows > 1:
        vSpace = spacingSpace / (numRows - 1)
    else:
        vSpace = spacingSpace

    # Final height values for rows respecting type, has to be in bottom-up order
    dataSetHeights = [] # Base unit to build height list from contains entries for one dataset
    # Set correct graph height based on row number and type
    rowOffset = 4  # Relative size of data tracks compared to gene model tracks
    baseHeight = 30  # Size of gene model row, for plot scaling
    if cfg.rawAvail == True:
        rawDataRows = numParams * rowOffset
    else:
        rawDataRows = 0
    if cfg.procAvail == True and rawDataRows > 0:
        procDataRows = numParams * 0.5
    else:
        procDataRows = 0
    if cfg.procAvail == True:
        dataSetHeights.append(rowHeight / 2 * bsScale)
    if cfg.rawAvail == True:
        dataSetHeights.append(rowHeight * rowOffset * iCLIPScale)
        
    rowHeights = [rowHeight] * numIsoforms + dataSetHeights * numParams + [rowHeight]
    blockHeight = 0.4
    fig = tools.make_subplots(print_grid=False, rows=numRows, cols=1, shared_xaxes=True, vertical_spacing=vSpace, row_width=rowHeights)
    for i in seqTrace:
        fig.append_trace(i, 1, 1)

    counter = 2
    
    for i in traces:
        if isinstance(i, list):
            for j in i:
                if isinstance(j, list):
                    for k in j:
                        fig.append_trace(k, counter, 1)
                else:
                    fig.append_trace(j, counter, 1)
        else:
            fig.append_trace(i, counter, 1)
        counter += 1    

    strand = figData['strand']
    fig['layout']['xaxis'].update(nticks=6)
    fig['layout']['xaxis'].update(tickmode='array')
    fig['layout']['xaxis'].update(showgrid=True)
    fig['layout']['xaxis'].update(ticks='outside')
    fig['layout']['xaxis'].update(ticksuffix='b')
    fig['layout'].update(hovermode='x')
    if strand == '-':
        fig['layout']['xaxis'].update(autorange='reversed')
    # The trailing ',' actually matters for some reason, don't remove
    fig['layout'].update(
        barmode='relative',
        margin=go.layout.Margin(l=30, r=40, t=25, b=60)
    )
    fig['layout']['yaxis'].update(visible=False, showticklabels=False, showgrid=False, zeroline=False)
    if cfg.procAvail:
        for i in range(0, numParams * cfg.dsElements, 2):
            fig['layout']['yaxis' + str(i + 3)].update(showticklabels=False, showgrid=False, zeroline=False)
    for i in range(numIsoforms):  # Edit all y axis in gene model plots
        fig['layout']['yaxis' + str(i + numParams * cfg.dsElements + 2)].update(showticklabels=False, showgrid=False,
                                                                            zeroline=False, range =[-blockHeight, blockHeight])
    for i in range(1,numRows + 1):  # Prevent zoom on y axis
        fig['layout']['yaxis' + str(i)].update(fixedrange=True)

    fig['layout']['height'] = (baseHeight * rawDataRows * iCLIPScale
                               + baseHeight * procDataRows *bsScale
                               + baseHeight * (numIsoforms + 1)
                               + 80)
    fig['layout']['legend'].update(x = legendColumnSpacing)            
    return fig


@app.callback(
    dash.dependencies.Output('bsGraphMem', 'data'),
    [dash.dependencies.Input('geneDrop', 'value')],
    [dash.dependencies.State('paramList', 'values'),
     dash.dependencies.State('sequenceRadio', 'value'),
     dash.dependencies.State('colorFinal', 'data'),
     dash.dependencies.State('legendSpacingDiv', 'data')]
)
def iCLIPCallback(geneName, dataSets, seqDisp, colorsFinal, legendSpacing):
    """Data callback that handles the selection of data and creates all possible traces.

    Positional arguments:
    geneName -- Name of the selected gene in order to filter the data.
    dataSets -- Selected data tracks with raw binding site data.
    seqDisp -- Display mode for dna sequence trace.
    colorsFinal -- Last confirmed color.
    legendSpacing -- Specifies margin between colorbar and other legend items.
    """
    dataSets = cfg.dataSetNames
    # Check which of the two triggering buttons was pressed last
    colors = colorsFinal
    # Dict that will store plot data, to be serialized later
    figData = {}
    # Sort the list of selected data tracks to keep consistent order
    for i in cfg.sortKeys:
        try:
            dataSets.sort(key=eval(i[0], {'__builtins__': None}, {}), reverse=eval(i[1], {'__builtins__': None}, {}))
        except (TypeError, SyntaxError):
            print(
                'Please check your keys. Each key should be added similar to this: -k \'lambda x : x[-2:]\' \'False\'	. For multiple keys use multiple instances of -k')
    # Select appropriate data from either the coding or non-coding set
    currentGene = pandas.DataFrame()
    for elem in cfg.geneAnnotations:
        currentGene = elem[elem['geneID'].str.contains(geneName)]
        if not currentGene.empty:
            break
    # Setup some variables for plot creation 
    xAxisMin = currentGene['chromStart'].min() # Left border of the plot region
    xAxisMax = currentGene['chromEnd'].max() # Rigt border of the plot region
    strand = currentGene['strand'].iloc[0] # Strand the selected gene is on
    chrom = currentGene['chrom'].iloc[0] # Chromosome the selected gene is on
   
    figData.update({'strand' : strand})
    overlappingGenes = []
    for i in cfg.geneAnnotations: # Select data for gene model from all annotaion files
        bcrit11 = i['chrom'] == chrom
        bcrit21 = i['chromStart'] >= xAxisMin
        bcrit22 = i['chromStart'] <= xAxisMax
        bcrit31 = i['chromEnd'] >= xAxisMin
        bcrit32 = i['chromEnd'] <= xAxisMax
        bcrit41 = i['chromStart'] <= xAxisMin
        bcrit42 = i['chromEnd'] >= xAxisMax
        preDF = i.loc[bcrit11 & ((bcrit21 & bcrit22) | (bcrit31 & bcrit32) | (bcrit41 & bcrit42))]
        result = preDF[~preDF['geneID'].str.contains(geneName)]
        overlappingGenes.append(result)
        
    overlaps = pandas.concat(overlappingGenes)
    isoformList = pandas.concat([currentGene, overlaps]) 

    # Create list of 3-tupels containing start, end, name for each isoform.
    isoformRanges = []
    for elem in currentGene.itertuples():
        name = elem.transID
        isoformRanges.append((elem.chromStart, elem.chromEnd, name))
    # Create master sequence for sequence display
    try:
        combinedSeq = generateMasterSequence(cfg.sequences, isoformRanges, xAxisMin, xAxisMax)
    except TypeError:
        combinedSeq = ''
    try:  # Create traces for sequence display, either scatter or heatmap
        figData.update({'heatSeq' : createSequenceTrace('heatSeq', strand, combinedSeq, xAxisMin, xAxisMax)})
        figData.update({'letterSeq' : createSequenceTrace('letterSeq', strand, combinedSeq, xAxisMin, xAxisMax)})
    except IndexError:
        pass
    except TypeError:
        pass

    iCLIPTraces = []
    for i in range(len(dataSets)):
        bsTraces = createICLIPTrace(dataSets[i], xAxisMax, xAxisMin, chrom, strand, colors)  # Plot binding site data
        iCLIPTraces.append(bsTraces)
    figData.update({'iCLIPTraces' : iCLIPTraces})
    # Calculate gene models. We have to distinguish between coding region and non-coding region
    blockHeight = 0.4
    geneModels = createGeneModelPlot(isoformList, xAxisMin, xAxisMax, blockHeight, strand)
    figData.update({'geneModels' : geneModels})
    return figData

def generateMasterSequence(sequences, isoforms, xAxisMin, xAxisMax):
    """Helper function that creates a master sequence given a dataframe with sequences and a list containing
        start and end points as well as names for the relevant isoforms.
    
    Positional arguments:
    sequences -- The list of sequence dicts.
    isoforms -- List containing three-tuples(start, end, name) for each isoform.
    xAxisMin -- Start point of the gene on the x-axis.
    xAxisMax -- End point of the gene on the x-axis, used for potential early termination.
    """  
    if xAxisMax <= 0:
        return ''
    try:
        if len(isoforms) == 0:
            return ''
    except TypeError:
        return ''
    # Sort tuples by start point. this ensures that the algorithm will cover the whole
    # gene sequence, if possible, albeit not necessarily with the least amount of subsequences
    isoforms.sort()
    # Initialize using the leftmost sequence
    seqDict = None
    try:
        for i in sequences: # Determine which dict contains the relevant sequences, all have to be in the same dict
            if isoforms[0][2] in i:
                seqDict = i
    except TypeError:
        return ''
    if seqDict == None:
        return ''
    combinedSeq = ''
    for elem in isoforms:
        if len(str(seqDict[elem[2]].seq)) == elem[1]- elem[0]:
            if elem[0] > xAxisMin:
                combinedSeq = ' '*(elem[0]-xAxisMin-1) + str(seqDict[elem[2]].seq)
            else:
                combinedSeq = str(seqDict[elem[2]].seq)
            currentEnd = elem[1]
            break
    if combinedSeq == '':
        return ''
    # Loop through elements and try to append sequences
    for elem in isoforms:
        provEnd = -1 # Used to fill blanks resulting from bad isoform names
        if elem[1] > currentEnd: 
            if elem[0] <= currentEnd: # current element overlaps and adds to the sequence
                if provEnd != -1: # Check if sequence before triggered a KeyError and fill gaps
                    if elem[0] >= provEnd:
                        combinedSeq += ' '*(elem[0]-currentEnd)
                        provEnd = -1
                    if elem[0] < provEnd:
                        combinedSeq += ' ' * (elem[0]-provEnd)
                        provEnd = -1
                try:
                    if len(str(seqDict[elem[2]].seq)) == (elem[1]- elem[0]):
                        combinedSeq += str(seqDict[elem[2]].seq)[(currentEnd - elem[0]):]
                        currentEnd = elem[1]           
                except KeyError:
                    provEnd = elem[1]
            else: # Current element does not overlap but will add to the sequence, fill with gaps
                try:
                    if len(seqDict[elem[2]].seq) == (elem[1]-elem[0]):
                        if provEnd != -1: # Check if sequence before triggered a KeyError and fill gaps
                            if elem[0] >= provEnd:
                                combinedSeq += ' '*(elem[0]-currentEnd)
                                provEnd = -1
                            if elem[0] < provEnd:
                                combinedSeq += ' ' * (elem[0]-provEnd)
                                provEnd = -1
                        fillerDist = elem[0] - currentEnd
                        combinedSeq += ' ' * fillerDist
                        combinedSeq += str(seqDict[elem[2]].seq)
                        currentEnd = elem[1]
                except KeyError:
                    provEnd = elem[1]
        if currentEnd >= xAxisMax: # The master sequence is complete, the entire region is covered
            break
    if provEnd != -1 or currentEnd < xAxisMax:
        combinedSeq += ' '*(xAxisMax-currentEnd)
    return combinedSeq

def createICLIPTrace(name, xMax, xMin, chrom, strand, colors):
    """Helper function to plot the subplots containing iCLIP data.
    
    Positional arguments:
    name -- Name of the subplot to create a title.
    xMax -- Maximum x-axis value, used to select relevant data.
    xMin -- Minimum x-axis value.
    chrom -- The chromosome we are looking for data on.
    strand -- Strand the gene is on, to look for correct binding sites.
    colors -- Colors for the traces.
    """
    colors = colors
    # Selection criteria
    crit1 = cfg.bsRawDFs[name]['chrom'] == chrom
    crit21 = cfg.bsRawDFs[name]['chromStart'] >= xMin
    crit22 = cfg.bsRawDFs[name]['chromStart'] <= xMax
    crit31 = cfg.bsRawDFs[name]['chromEnd'] >= xMin
    crit32 = cfg.bsRawDFs[name]['chromEnd'] <= xMax
    rawSites = cfg.bsRawDFs[name].loc[crit1 & ((crit21 & crit22) | (crit31 & crit32))]
    # Setup arrays to hold the values that will be needed for plotting
    countsX = []
    countsY = []
    countsW = []
    for i in rawSites.itertuples():
        countsX.append(i.chromStart)
        countsY.append(i.count)
        countsW.append(i.chromEnd - i.chromStart)
    # Plot data
    rawTrace = go.Bar(
        x=countsX,
        y=countsY,
        width=countsW,
        hoverinfo='x+y',
        name=name,
        meta = name,
        legendgroup=name,
        marker=go.bar.Marker(
            color=colors[name]
        ),
        showlegend=True
    )

    # Setup criteria to select binding sites that are within the current region of the genome
    procSitesList = []
    try:
        bcrit11 = cfg.bsProcDFs[name]['chrom'] == chrom
        bcrit12 = cfg.bsProcDFs[name]['strand'] == strand
        bcrit21 = cfg.bsProcDFs[name]['chromStart'] >= xMin
        bcrit22 = cfg.bsProcDFs[name]['chromStart'] <= xMax
        bcrit31 = cfg.bsProcDFs[name]['chromEnd'] >= xMin
        bcrit32 = cfg.bsProcDFs[name]['chromEnd'] <= xMax
        bindingSites = cfg.bsProcDFs[name].loc[bcrit11 & bcrit12 & ((bcrit21 & bcrit22) | (bcrit31 & bcrit32))]
        # Plot binding sites
        for k in bindingSites.itertuples():
                procSitesList.append(
                    go.Bar(
                        opacity = 0.5,
                        x = [k.chromStart + (k.chromEnd - k.chromStart) // 2],
                        y = [0.1],
                        hoverinfo = 'name',
                        legendgroup = name,
                        width = k.chromEnd - k.chromStart,
                        name = name + '_bs',
                        meta = name,
                        marker = go.bar.Marker(
                            color = colors[name]
                        ), showlegend = False
                    )
                )
    except KeyError:
        pass
    except Exception as e:
        print('Error in binding plot: ' + str(type(e).__name__) + str(e.args))
    return [rawTrace, procSitesList]


def setUpBlockConstraints(chromStart, chromEnd, blockStarts, blockSizes, xAxisMin, xAxisMax):
    """ This function will calculate the correct block start and block size values.
        It takes partial overlapping of blocks with the target area into account.
        Any block that is outside (xAxisMin, xAxisMax) will have a blockStart of -1,
        indicating that it i to be ignored when drawing the gene model.
        
        Positional arguments:
        chromStart -- Isoform start point.
        chromEnd -- Isoform end point.
        blockStarts -- Block startpoints.
        blockSizes -- Sizes for the blocks.
        xAxisMin -- Start of the relevant genomic region.
        xAxisMax -- End of the relevant genomic region.
        
        Returns:
        Tupel containing finished block starts and block sizes as lists:
        (blockStartsF, blockSizesF)
    """
    if chromStart >= xAxisMin:
        if chromEnd <= xAxisMax: # Gene contained entirely in region, no cutting necessary
            loc = 'cont'
            blockStartsF = [int(x) + chromStart for x in blockStarts.rstrip(',').split(',')]
            blockSizesF = [int(x) for x in blockSizes.rstrip(',').split(',')]
        else: # Gene overlaps region on right, cut end
            loc = 'left'
            blockStartsF = [int(x) + chromStart for x in blockStarts.rstrip(',').split(',')]
            blockSizesF = [int(x) for x in blockSizes.rstrip(',').split(',')]
            for index, elem in enumerate(blockStartsF):
                if elem >= xAxisMax: # The current block starts right of the relevant region, disregard
                    blockStartsF[index] = -1
                else:
                    blockEnd = elem + blockSizesF[index]
                    if  blockEnd > xAxisMax:
                        blockSizesF[index] = blockSizesF[index] - (blockEnd-xAxisMax)
    else: 
        if chromEnd <= xAxisMax: # Gene overlaps on the left, cut start
            loc = 'right'
            blockStartsF = [int(x) + chromStart for x in blockStarts.rstrip(',').split(',')]
            blockSizesF = [int(x) for x in blockSizes.rstrip(',').split(',')]
            for index, elem in enumerate(blockStartsF):
                if elem + blockSizesF[index] < xAxisMin: # Block ends left of relevant regeion, disregard
                    blockStartsF[index] = -1
                else: 
                    if  elem < xAxisMin: 
                        startOffset = xAxisMin - elem
                        blockStartsF[index] = xAxisMin
                        blockSizesF[index] = blockSizesF[index] - startOffset
        else: # Gene extends to the left and right of the region, cut on both ends
            loc = 'both'
            blockStartsF = [int(x) + chromStart for x in blockStarts.rstrip(',').split(',')]
            blockSizesF = [int(x) for x in blockSizes.rstrip(',').split(',')]
            for index, elem in enumerate(blockStartsF):
                blockEnd = elem + blockSizesF[index]
                if  blockEnd < xAxisMin or elem >= xAxisMax: # Block lies left or right of relevant region, disregard
                    blockStartsF[index] = -1
                else:
                    if  elem < xAxisMin: # Block overlaps on the left
                        startOffset = xAxisMin - elem
                        blockStartsF[index] = xAxisMin
                        blockSizesF[index] = blockSizesF[index] - startOffset
                    if blockEnd > xAxisMax: # Block overlaps on the right
                        blockSizesF[index] = blockSizesF[index] - (blockEnd-xAxisMax)
    # Additional checks for bad inputs
    for index, elem in enumerate(blockStartsF):
        if elem > xAxisMax:
            blockStartsF[index] = -1 
        else:
            if elem + blockSizesF[index] > xAxisMax:
                blockSizesF[index] = xAxisMax - elem
    return (blockStartsF, blockSizesF, loc)
                
def calculateBlocks(thickStart, thickEnd, blockStart, blockEnd, blockVals, blockWidths, blockYs, blockHeight, strand):
    """ This function determines the actual shape of each block, based on wether it's
        located in coding or noncoding region, or on the boundary. The function directly appends to the input lists
        
        Positional Arguments:
        thickStart -- The start point of the coding region.
        thickEnd -- Endpoint of the coding region.
        blockStart -- Startin point of the block. A value of -1 means this block is not in the relevant area .
        and will be ignored
        blockEnd -- The end point of the block.
        blockVals -- This lists holds the x-coordinate for each block center.
        blockWidths -- This list holds the width of each block.
        blockYa -- Holds the y value of the block, this determines wether it's a coding or noncoding block.
        blockHeight -- The height value to use for coding blocks. Non coding blocks are half height.
    """
    blockStart = blockStart
    if blockStart != -1 and blockStart != blockEnd:
        blockEnd = blockEnd  # Same as codingRegionEnd
        codingRegionStart = int(thickStart)
        codingRegionEnd = int(thickEnd)
        if (blockStart >= codingRegionStart) & (blockEnd <= codingRegionEnd):
            # Block is inside coding region
            blockVals.append(blockStart + ((blockEnd-1) - blockStart) / 2)
            blockWidths.append(blockEnd - blockStart )
            blockYs.append(blockHeight)
        if (blockStart >= codingRegionStart) & (blockEnd > codingRegionEnd):
            if (blockStart >= codingRegionEnd):
                # Block is right of coding region
                blockVals.append(blockStart + ((blockEnd-1) - blockStart) / 2)
                blockWidths.append(blockEnd - blockStart )
                blockYs.append(blockHeight / 2)
            else:
                # Block overlaps coding region on the left
                blockVals.append(blockStart + ((codingRegionEnd -1)- blockStart) / 2)
                blockWidths.append(codingRegionEnd - blockStart )
                blockYs.append(blockHeight)
                blockVals.append((codingRegionEnd-1)+ (blockEnd - (codingRegionEnd-1)) / 2)
                blockWidths.append(blockEnd - (codingRegionEnd ))
                blockYs.append(blockHeight / 2)
        if (blockStart < codingRegionStart) & (blockEnd <= codingRegionEnd):
            if blockEnd <= codingRegionStart:
                # Block is Left of coding region
                blockVals.append(blockStart + ((blockEnd-1) - blockStart) / 2)
                blockWidths.append(blockEnd - blockStart)
                blockYs.append(blockHeight / 2)
            else:
                # Block overlaps coding region on the right
                blockVals.append(blockStart + ((codingRegionStart-1) - blockStart) / 2)
                blockWidths.append((codingRegionStart-1) - blockStart + 1)
                blockYs.append(blockHeight / 2)
                blockVals.append(codingRegionStart + (blockEnd - (codingRegionStart+1)) / 2)
                blockWidths.append(blockEnd - codingRegionStart)
                blockYs.append(blockHeight)
        if (blockStart < codingRegionStart) & (blockEnd > codingRegionEnd):
            # Block completely contains coding region
            blockVals.append(blockStart + ((codingRegionStart-1) - blockStart) / 2)
            blockWidths.append((codingRegionStart-1) - blockStart + 1)
            blockYs.append(blockHeight / 2)
            blockVals.append(codingRegionStart + (codingRegionEnd - codingRegionStart) / 2)
            blockWidths.append(codingRegionEnd - codingRegionStart + 1)
            blockYs.append(blockHeight)
            blockVals.append(codingRegionEnd + (blockEnd - codingRegionEnd) / 2)
            blockWidths.append(blockEnd - (codingRegionEnd + 1))
            blockYs.append(blockHeight / 2)

def createSequenceTrace(seqDisp, strand, combinedSeq, xAxisMin, xAxisMax):
    """ Function to generate sequence display trace, either heatmap or scatter

    Positional arguments:
    seqDisp -- Determines which trace type is used.
    strand -- If on minus strand invert dna sequence.
    combinedSeq -- Sequence for display.
    xAxisMin -- Startpoint.
    xAxisMax -- Endpoit.
    """
    if seqDisp == 'letterSeq':
        xA = []
        xC = []
        xG = []
        xT = []
        Err = []
        if strand == '+':
            varMap = {'A': xA, 'C': xC, 'G': xG, 'T': xT}
        else:  # Reverse complement
            varMap = {'A': xT, 'C': xG, 'G': xC, 'T': xA}
        for i in range(xAxisMin, xAxisMax):
            try:
                varMap[combinedSeq[i - xAxisMin]].append(i)
            except KeyError:
                Err.append(i)
        aTrace = go.Scatter(
            text=['A'] * len(xA),
            textfont=dict(color=cfg.colorA),
            mode='text',
            name='seqA',
            y=[1] * len(xA),
            x=xA,
            showlegend=False,
            opacity=0.5,
            hoverinfo='x',
            textposition="bottom center"
        )
        cTrace = go.Scatter(
            text=['C'] * len(xC),
            mode='text',
            textfont=dict(color=cfg.colorC),
            y=[1] * len(xC),
            x=xC,
            name='seqC',
            showlegend=False,
            opacity=0.5,
            hoverinfo='x',
            textposition="bottom center"
        )
        gTrace = go.Scatter(
            text=['G'] * len(xG),
            textfont=dict(color=cfg.colorG),
            mode='text',
            y=[1] * len(xG),
            name='seqG',
            x=xG,
            showlegend=False,
            opacity=0.5,
            hoverinfo='x',
            textposition="bottom center"
        )
        tTrace = go.Scatter(
            text=['T'] * len(xT),
            textfont=dict(color=cfg.colorT),
            mode='text',
            y=[1] * len(xT),
            name='seqT',
            x=xT,
            showlegend=False,
            opacity=0.5,
            hoverinfo='x',
            textposition="bottom center"
        )
        errorTrace = go.Scatter(
            text=['N'] * len(Err),
            textfont=dict(color='rgb(0,0,0)'),
            mode='text',
            name='seqN',
            y=[1] * len(Err),
            x=Err,
            showlegend=False,
            opacity=0.5,
            hoverinfo='x',
            textposition="bottom center"
        )
        return [aTrace, cTrace, gTrace, tTrace, errorTrace]
    if seqDisp == 'heatSeq':
        colorE = 'rgb(0, 0, 0)'
        if strand == '+':
            valMap = {'A': 0, 'C': 2, 'G': 3, 'T': 1}
            textMap = {'A': 'A', 'C': 'C', 'G': 'G', 'T': 'T'}
        else:  # Reverse complement
            valMap = {'A': 1, 'C': 3, 'G': 2, 'T': 0}
            textMap = {'A': 'T', 'C': 'G', 'G': 'C', 'T': 'A'}
        zlist = []
        textList = []
        errorsPresent = False
        for i in range(xAxisMin, xAxisMax):
            try:
                zlist.append(valMap[combinedSeq[i - xAxisMin]])
                textList.append(textMap[combinedSeq[i - xAxisMin]])
            except KeyError:
                errorsPresent = True
                zlist.append(4)
                textList.append('N')
        if errorsPresent == True:
            colors = [
                [0, cfg.colorA],
                [0.2, cfg.colorA],
                [0.2, cfg.colorT],
                [0.4, cfg.colorT],
                [0.4, cfg.colorC],
                [0.6, cfg.colorC],
                [0.6, cfg.colorG],
                [0.8, cfg.colorG],
                [0.8, colorE],
                [1.0, colorE]
            ]
        else:
            colors = [
                [0, cfg.colorA],
                [0.25, cfg.colorA],
                [0.25, cfg.colorT],
                [0.5, cfg.colorT],
                [0.5, cfg.colorC],
                [0.75, cfg.colorC],
                [0.75, cfg.colorG],
                [1.0, cfg.colorG]
            ]
        if strand == '-':
            xList = list(range(xAxisMin, xAxisMax-1))
        else:
            xList = list(range(xAxisMin, xAxisMax))
        heatTrace = go.Heatmap(
            z=[zlist],
            x=xList,
            text=[textList],
            colorscale=colors,
            showscale=True,
            name='seq',
            hoverinfo='x+text',
            colorbar={
                'x' : 1.0,
                'y' : 0.0,
                'tickmode' : 'array',
                'tickvals' : [0.4,1.1,1.86,2.6  ],
                'ticktext' : ['A', 'T', 'C', 'G'],
                'yanchor' : 'bottom',
                'len' : 1.0
            }
        )
        return [heatTrace]


def createGeneModelPlot(isoforms, xAxisMin, xAxisMax, blockHeight, strand):
    """Generates gene model based on the given blocks and coding region.

    Positional arguments:
    isoforms -- List of all isoforms overlapping relevant region.
    xAxisMin -- Start of the relevant region.
    xAxisMax -- End of the relevant region.
    blockHeight -- Hight for thick blocks.
    strand -- Strand that the selected gene is on, for coloring.
    """
    traces = []
    # This loop will check and if necessary cut blockstarts and sizes
    # depending on the form of overlap
    for i in isoforms.itertuples():
        # Set color depending on the strand the isoform is located on
        if i.strand != strand: # Isoform is on reverse strand, color grey
            color = 'rgb(128,128,128)'
            sign = -1
        else: # Isoform is on same strand as selected gene, color black
            color = 'rgb(0,0,0)'
            sign = 1
        # Calculate proper blockStarts and blockSizes
        blockConstraints = setUpBlockConstraints(i.chromStart, i.chromEnd, i.blockStarts, i.blockSizes, xAxisMin, xAxisMax)
        blockStarts = blockConstraints[0]
        blockSizes= blockConstraints[1]
        loc = blockConstraints[2] 
        # Lists for the values needed to draw traces
        blockVals = []
        blockWidths = []
        blockYs = []
        name = i.transID
        # Calculate blocks from block start and end positions, as well as thickness
        for j in range(len(blockStarts)):
            blockStart = blockStarts[j]
            calculateBlocks(i.thickStart, i.thickEnd, blockStart, blockStart + blockSizes[j] , blockVals, blockWidths, blockYs, blockHeight, strand)
        # Find first and last block to draw line properly
        f = lambda i: blockVals[i]
        lineCoords = []
        lineName = ''
        lineHover = 'none'
        lineLegend = False
        try: # Draw line from first to last Exon
            amaxBlockVals = max(range(len(blockVals)), key=f)
            aminBlockVals = min(range(len(blockVals)), key=f)
            if loc == 'left':
                lineCoords.append(blockVals[aminBlockVals] - blockWidths[aminBlockVals] / 2)
                if sign > 0 :
                    lineCoords.append(xAxisMin)
                else:
                    lineCoords.append(xAxisMax-1)                      
            elif loc == 'right':
                lineCoords.append(blockVals[amaxBlockVals] + blockWidths[amaxBlockVals] / 2)
                if sign > 0 :
                    lineCoords.append(xAxisMax-1)
                else:
                    lineCoords.append(xAxisMin)   
            elif loc == 'both':
                lineCoords.append(xAxisMax-1)
                lineCoords.append(xAxisMin)  
            else:
                lineCoords.append(blockVals[amaxBlockVals] + blockWidths[amaxBlockVals] / 2)
                lineCoords.append(blockVals[aminBlockVals] - blockWidths[aminBlockVals] / 2)
        except ValueError: # Case that no exons are present and only a line is drawn
            lineCoords.append(xAxisMin)
            lineCoords.append(xAxisMax)
            lineName = name
            lineHover = 'name'
            lineLegend = True
        # The line of the gene model
        line = go.Scatter(
            x = lineCoords,
            y = [0, 0],
            name = lineName,
            hoverinfo = lineHover,
            hoverlabel = {
                'namelength' : -1, },
            mode = 'lines',
            line = dict(
                color = color,
            ),
            showlegend = lineLegend,
            legendgroup = name
        )
        # Trace for the blocks of the gene model
        blocks = go.Bar(
            x = blockVals,
            y = [i * 2 for i in blockYs],
            base = [0 - (i / 2) * 2 for i in blockYs],
            name = name,
            hoverinfo = 'name',
            hoverlabel = {
                'namelength' : -1, },
            width = blockWidths,
            marker = go.bar.Marker(
                color = color
            ),
            showlegend = True,
            legendgroup = name
        )
        traces.append([line, blocks])
    # Return traces for gene model
    return traces
```
