## Additional File 1 for "SEQing: web-based visualization of iCLIP and RNA-seq data in an interactive python framework": rna_tab.py

```
#!/usr/bin/env python3
# -*- coding: utf-8 -*-

import dash
import pandas
from app import app
import cfg
import time 
import pickle
import dash_html_components as html
from plotly import tools
import plotly.graph_objs as go
from iclip_tab import createGeneModelPlot
import plotly.utils as pu

@app.callback(
    dash.dependencies.Output('rnaDescDiv', component_property='children'),
    [dash.dependencies.Input('geneDrop', 'value')],
)
def rnaDesc(name):
    if cfg.descAvail:
        try:
            return [
                html.P(
                    cfg.geneDescriptions.loc[
                        cfg.geneDescriptions['ensembl_gene_id'] == name,
                        ['description']
                    ].iloc[0]
                )
            ]
        except IndexError:
            return ['No description available']
        except KeyError:
            return ['No description available']
    else:
        return ['No description available']

@app.callback(
    dash.dependencies.Output('spliceGraph', 'figure'),
    [dash.dependencies.Input('spliceMem', 'data'),
     dash.dependencies.Input('rnaParamList', 'values'),
     dash.dependencies.Input('rnaRadio', 'value'),
     dash.dependencies.Input('covColorFinal', 'data'),
     dash.dependencies.Input('eventColorFinal', 'data'),
     dash.dependencies.Input('legendSpacingDiv', 'data'),
     dash.dependencies.Input('coverageScale', 'value'),
     dash.dependencies.Input('sequenceRadio', 'value'),
     dash.dependencies.Input('eventScale', 'value'),
     dash.dependencies.Input('bsGraphMem', 'data')]
)
def showRNA(figData, dataSets, displayType, covColor, eventColor, legendSpacing, coverageScale, seqDisp, eventScale, bsMem):
    """Update callback that selects traces to be displayed based on settings.

    Positional arguments:
    figData -- Trace data from the data callback.
    datasets -- List of datasets to display.
    displayType -- Type of splice event display.
    covColor -- Colors for coverage traces.
    eventColorl -- Colots for splice events.
    legendSpacing -- Specifies margin between colorbar and other legend items.
    coverageScale -- Scaling factor for coverage plots.
    eventScale -- Scaling factor for event plots.
    """
    legendColumnSpacing = legendSpacing
    figData = figData
    traces = figData['rnaTraces']
    geneModels = figData['geneModels']
    coverageColors = covColor
    try:
        seqTrace = bsMem[seqDisp]
        if seqDisp == 'heatSeq':
            for i in seqTrace:
                i['showscale'] = False
    except:
        seqTrace = []
    #print(seqTrace)
    eventColors = eventColor
    eventIndices = [] # Save indices of all elements that contain event traces
    rnaDataSets = sorted(list(cfg.coverageData.keys()))
    displayed_rnaDataSet = []
    maxYDict = figData['maxYList']
    axisTitles = []
    yVals = []
    for rm in sorted(dataSets):
        for set in rnaDataSets:
            if rm == set.split('_')[0]:
                displayed_rnaDataSet.append(set)
    finTraces = []
    eventIndices = []
    for index, t in enumerate(traces):
        try:
            if len(t[displayType]) > 1:
                try:
                    if t[displayType][0]['meta'] in displayed_rnaDataSet:
                        if displayType == 'two':
                            for i in t[displayType]:
                                newColor = eventColors[i['name']]
                                i['marker'] = {'color' : newColor}
                        finTraces.append(t[displayType])      
                        eventIndices.append(index//2)
                        axisTitles.append('')
                except KeyError:
                    if t[displayType]['meta'] in displayed_rnaDataSet:
                        if displayType == 'two':
                            newColor = eventColors[t['name']]
                            t['marker'] = {'color' : newColor}
                        finTraces.append(t[displayType])      
                        eventIndices.append(index//2)
                        axisTitles.append('')
            else:
                if t[displayType][0] in displayed_rnaDataSet:
                    finTraces.append([])
                    axisTitles.append('')
                    eventIndices.append(index//2)

        except KeyError:
            if t['meta'] in displayed_rnaDataSet:
                newColor = coverageColors[t['meta'].split('_')[0]]
                yVals.append(maxYDict[t['meta']])
                axisTitles.append('')
                t['fillcolor'] = newColor
                finTraces.append(t)              
    numIsoforms = len(geneModels) # Number of isoforms in the gene model
    numRows = len(finTraces)+numIsoforms+1#+1 for sequence trace
    
    # Setup row heights based on available data
    
    plotSpace = 0.9  # Space taken up by data tracks
    spacingSpace = 1.0 - plotSpace  # Space left for spacer tracks
    rowHeight = plotSpace / numRows
    if numRows > 1:
        vSpace = spacingSpace / (numRows - 1)
    else:
        vSpace = spacingSpace
    rowHeights = []
    rowHeights.append(rowHeight/2)
    eventHeights = []
    eventMaxHeights = figData['maxHeights']
    for index, i in enumerate(eventMaxHeights):
        if index in eventIndices:
            if i == 0:
                eventHeights.append(0)           
            if i > 0 and i <= 5:
                eventHeights.append(1)
            if i >= 6 and i < 10:
                eventHeights.append(2)
            if i >= 10:
                eventHeights.append(i % 5 +1)
    if cfg.spliceEventAvail:
        for i in range(1,numRows):
            if i > len(finTraces): 
                rowHeights.append(0.5 * rowHeight) # Gene model row
            elif (i % 2 == 0):
                try:
                    rowHeights.append(eventHeights[(i//2)-1] * rowHeight * eventScale) # Splice event row
                except IndexError:
                    rowHeights.append(0)
            else:
                rowHeights.append(3 * rowHeight * coverageScale) # Coverage row
    else:
        for i in range(1,numRows):
            if i > len(finTraces): rowHeights.append(0.5 * rowHeight) # Gene model row
            else:
                rowHeights.append(3 * rowHeight * coverageScale) # Coverage row
    fig = tools.make_subplots(print_grid=False, rows=numRows, cols=1,
                              shared_xaxes=True, row_width=rowHeights[::-1], vertical_spacing = vSpace)
        # Layouting of the figure
    eventIndicesDraw = [] # Save indices of all elements that contain event traces
    for i in seqTrace:
        fig.append_trace(i, 1, 1)
    for index, t in enumerate(finTraces):
        try:
            fig.append_trace(t, index + 2, 1)
        except ValueError:
            eventIndicesDraw.append(index)
    for i in eventIndicesDraw: # Add event traces after all coverage traces have been added for legend item positioning
        for x in finTraces[i]:
            fig.append_trace(x, i + 2, 1)    
    counter = len(finTraces)+1
    for model in geneModels:
        for part in model:
            fig.append_trace(part, counter+1, 1)
        counter += 1
    fig['layout']['xaxis'].update(nticks=6)
    fig['layout']['xaxis'].update(tickmode='array')
    fig['layout']['xaxis'].update(showgrid=True)
    fig['layout']['xaxis'].update(ticks='outside')
    fig['layout']['xaxis'].update(ticksuffix='b')
    fig['layout']['xaxis'].update(ticksuffix='b')
    fig['layout'].update(hovermode='closest')
    fig['layout']['yaxis'].update(fixedrange=True)
    fig['layout'].update(barmode='relative')

    # Reverse x-axis if gene is on - strand to always show models in 3'->5'
    if figData['strand'] == '-':
        fig['layout']['xaxis'].update(autorange='reversed')
    for i in range(1, numRows+1):  # prevent zoom on y axis
        fig['layout']['yaxis' + str(i)].update(fixedrange=True)
    try:
        maxYVal = max(yVals)
    except ValueError:
        maxYVal = 0
    blockHeight = 0.4
    for i in range(1, numRows):   
            if cfg.spliceEventAvail:
                if i % 2 != 0 and i <= len(finTraces): # Coverage row
                    fig['layout']['yaxis' + str(i+1)].update(range=[0, maxYVal],title={'text': axisTitles[i-1]})
                    fig['layout']['yaxis' + str(i+1)].update(showticklabels=True, showgrid=True, zeroline=True)
                else: # Event or gene model row
                    if i > len(finTraces): # Gene model row
                        fig['layout']['yaxis' + str(i+1)].update(showticklabels=False, showgrid=False, zeroline=False)
                        fig['layout']['yaxis' + str(i+1)].update(range=[-blockHeight, blockHeight])
                    else: # Event row
                        fig['layout']['yaxis' + str(i+1)].update(showticklabels=False, showgrid=False, zeroline=False, title={'text': axisTitles[i-1]})
            else:
                if i <= len(finTraces): # Coverage row
                    fig['layout']['yaxis' + str(i+1)].update(range=[0, maxYVal], title={'text': axisTitles[i-1]})
                    fig['layout']['yaxis' + str(i+1)].update(showticklabels=True, showgrid=True, zeroline=True)
                else: # Gene model row
                    fig['layout']['yaxis' + str(i+1)].update(showticklabels=False, showgrid=False, zeroline=False)
                    fig['layout']['yaxis' + str(i+1)].update(range=[-blockHeight, blockHeight], )
    # Setup plot height, add 85 to account for margins
    fig['layout'].update(margin=go.layout.Margin(l=60, r=40, t=25, b=60),)
    fig['layout']['yaxis'].update(visible = False, showticklabels=False, showgrid=False, zeroline=False)
    rowScales = [x/rowHeight for x in rowHeights]
    size = 0 
    for i in rowScales:
        size += 50*i
    fig['layout']['height'] = (size + 85)
    # set spacing for the second legend column
    fig['layout']['legend'].update(x = legendColumnSpacing)
    #print('Showcallback: ' + str(end-start))
    return fig

@app.callback(
    dash.dependencies.Output('spliceMem', 'data'),
    [dash.dependencies.Input('geneDrop', 'value')],
    [dash.dependencies.State('rnaRadio', 'value'),
     dash.dependencies.State('rnaParamList', 'values'),
     dash.dependencies.State('covColorFinal', 'data'),
     dash.dependencies.State('eventColorFinal', 'data'),
     dash.dependencies.State('legendSpacingDiv', 'data'),
     dash.dependencies.State('coverageScale', 'value'),
     dash.dependencies.State('eventScale', 'value')]
)
def rnaCallback(geneName, displayMode,rnaParamList, colorsFinal, eventColorsFinal, legendSpacing,
                coverageScale, eventScale):
    """Data callback that selects relevant data and creates all possible traces.

        Positional arguments:
        geneName -- Name of the selected gene in order to filter the data.
        displaymode --determines how splice events will be visualized.
        rnaParamList -- Selected RNA data sets to plot.
        colorsFinal -- Last confirmed color.
        eventColorsFinal -- Last confirmed colors for splice events.
        legendSpacing -- Specifies margin between colorbar and other legend items.
        coverageScale -- Scaling factor for coverage plots.
        eventScale -- Scaling factor for event plots.
        """
    colors = colorsFinal
    figData = {}
    
    # Select appropriate data from gene annotations
    currentGene = pandas.DataFrame()
    for index, elem in enumerate(cfg.geneAnnotations):
        currentGene = elem[elem['geneID'].str.contains(geneName)]
        if not currentGene.empty:
            break

    # Get axis minimum and maximum over all isoforms. Also get current chromosome
    xAxisMax = currentGene['chromEnd'].max()
    xAxisMin = currentGene['chromStart'].min()
    chrom = currentGene['chrom'].iloc[0]
    strand = currentGene['strand'].iloc[0]
    figData.update({'strand': strand})
    color_dict = colors  # Color per mutant
    figData.update({'covColors' : color_dict})
    # Filter out needed datasets
    rnaDataSets = sorted(list(cfg.coverageData.keys()))
    displayed_rnaDataSet = rnaDataSets
   # for rm in sorted(rnaParamList):
    #    for set in rnaDataSets:
     #       if rm == set.split('_')[0]:
      #          displayed_rnaDataSet.append(set)

    # Dicts for lists of axis values
    xVals = {}
    yVals = {}
    maxYVal = 0 # Used to scale y-axes later
    maxYVals = {}
    eventDict = {} # stores dataframes with relevant splice event data
    iterTime = 0
    evSel = 0
    covSel = 0
    for ds in sorted(displayed_rnaDataSet): # Select relevant coverage files from Index
        covStart = time.time()
        spliceSlice = coverageDataSelection(ds, xAxisMin, xAxisMax, chrom)
        covEnd = time.time()
        covSel += covEnd-covStart
        # Pre-init y-value list
        yVal = [0] * (len(range(xAxisMin, xAxisMax)))
        organism = ds.split("_")[0] # Prefix of the curret data frame, first filter
        spliceEvents = pandas.DataFrame() # will hold splice event data for the current data set
        evStart = time.time()
        if any(organism in s for s in cfg.spliceEventNames[1]): # Check if there are splice events for the current prefix
            for d in sorted(cfg.spliceEventDFs.keys()):
                if ds in d: # Check for remaining filename, to match the correct files
                    # Criteria to filter relevant lines from current dataframe
                    bcrit11 = cfg.spliceEventDFs[d]['chrom'] == chrom
                    bcrit21 = cfg.spliceEventDFs[d]['chromStart'] >= xAxisMin
                    bcrit22 = cfg.spliceEventDFs[d]['chromStart'] <= xAxisMax
                    bcrit31 = cfg.spliceEventDFs[d]['chromEnd'] >= xAxisMin
                    bcrit32 = cfg.spliceEventDFs[d]['chromEnd'] <= xAxisMax
                    spliceEvents = cfg.spliceEventDFs[d].loc[bcrit11 & ((bcrit21 & bcrit22) | (bcrit31 & bcrit32))]
        # Use itertuples to iterate over rows, since itertuples is supposed to be faster
        evEnd = time.time()
        evSel += evEnd-evStart
        iterStart = time.time()
        for row in spliceSlice.itertuples():
            # Increment all values covered by the current row, will overshoot when row crosses border of gene, thus try except
            for j in range(row.chromStart, row.chromEnd):
                if (j - xAxisMin) >=0: # Ensure we don't fall into negative list indices
                    try:
                        yVal[j - xAxisMin] += row.count
                    except IndexError:
                            pass
        iterEnd = time.time()
        iterTime += iterEnd-iterStart
         # Store reference to value list in dict
        yVals[ds] = yVal
        # Safe event dataframe to be used in the next function
        eventDict[ds] = spliceEvents
        # Create x-axis values
        xVal = list(range(xAxisMin, xAxisMax))
        xVals[ds] = xVal
        # Find maximum y-axis value for axis scaling
        maxYVals.update({ds: max(yVal)})
        if max(yVal) > maxYVal: maxYVal = max(yVal)
    figData.update({'maxY' : maxYVal})
    figData.update({'maxYList' : maxYVals})
    # Create RNA-seq traces from data
    rnaSeqPlotData = createRNAPlots(xVals, yVals, eventDict, displayed_rnaDataSet, 
                          color_dict, displayMode, eventColorsFinal)
    traces = rnaSeqPlotData[0]
    eventMaxHeights = rnaSeqPlotData[1]
    axisTitles = rnaSeqPlotData[2]
    figData.update({'rnaTraces' : traces})
    figData.update({'maxHeights' : eventMaxHeights})
    figData.update({'axisTitles' : axisTitles})
    overlappingGenes = []
    for i in cfg.geneAnnotations: # Select data for gene models from all annotation files
        bcrit11 = i['chrom'] == chrom
        bcrit21 = i['chromStart'] >= xAxisMin
        bcrit22 = i['chromStart'] <= xAxisMax
        bcrit31 = i['chromEnd'] >= xAxisMin
        bcrit32 = i['chromEnd'] <= xAxisMax
        bcrit41 = i['chromStart'] <= xAxisMin
        bcrit42 = i['chromEnd'] >= xAxisMax
        preDF = i.loc[bcrit11 & ((bcrit21 & bcrit22) | (bcrit31 & bcrit32) | (bcrit41 & bcrit42))]
        result = preDF[~preDF['geneID'].str.contains(geneName)]
        overlappingGenes.append(result)
        
    overlaps = pandas.concat(overlappingGenes)
    isoformList = pandas.concat([currentGene, overlaps]) 
    blockHeight = 0.4

    # Calculate gene models. We have to distinguish between coding region and non-coding region
    geneModels = createGeneModelPlot(isoformList, xAxisMin, xAxisMax, blockHeight, strand)
    figData.update({'geneModels' : geneModels})
    return figData

def coverageDataSelection(ds, xAxisMin, xAxisMax, chrom):
    """ This function performs selection and loading of relevant coverage data. 
        Due to size, coverage data is indexed, and only needed files will be laoded
        on demand.
        
        Positional arguments:
        ds -- Name of the dataset .
        xAxisMin -- Left border of relevant area.
        xAxisMax -- Right border of relevant area.
        chrom -- Chromosome to search on.
    """
    fileIndex = cfg.coverageData[ds]
    dfBcrit11 = fileIndex['start'] <= xAxisMin
    dfBcrit12 = fileIndex['end'] >= xAxisMin
    dfBcrit21 = fileIndex['start'] <= xAxisMax
    dfBcrit22 = fileIndex['end'] >= xAxisMax
    dfBcrit31 = fileIndex['start'] >= xAxisMin
    dfBcrit32 = fileIndex['end'] <= xAxisMax
    relevantFiles = fileIndex.loc[(dfBcrit11 & dfBcrit12) | (dfBcrit21 & dfBcrit22) | (dfBcrit31 & dfBcrit32)]
    finalDF = []
    for i in relevantFiles.itertuples(): # Build dataframe from parts
        try:
            df = pickle.load(open(i.fileName, 'rb'))
            finalDF.append(df)
        except FileNotFoundError:
            print(i.fileName + ' was not found')
    try: # Final selection of relevant values from the combined df
        finalDF = pandas.concat(finalDF)
        bcrit11 = finalDF['chrom'] == chrom
        bcrit21 = finalDF['chromStart'] >= xAxisMin
        bcrit22 = finalDF['chromStart'] <= xAxisMax
        bcrit31 = finalDF['chromEnd'] >= xAxisMin
        bcrit32 = finalDF['chromEnd'] <= xAxisMax
        return finalDF.loc[bcrit11 & ((bcrit21 & bcrit22) | (bcrit31 & bcrit32))]
    except ValueError:
        return pandas.DataFrame()    

def overlap(a, b):
    """check if two intervals overlap.

    Positional arguments:
    a -- First interval.
    b -- Second interval.
    """
    return a[1] > b[0] and a[0] < b[1]


def createRNAPlots(xVals, yVals, eventData, displayed, colorDict,
                    displayMode, 
                    eventColorsFinal):
    """Create the plots for both coverage and splice events.

    Positional arguments:
    xVals -- x-axis values for coverage plot.
    yVals -- y-axis values for coverage plot.
    eventData -- Dict containing the dataframes with relevant splice events.
    displayed -- Displayed datasets.
    colorDict -- Colors for the coverage plots.
    displayMode -- Determines how splice events are visualized.
    eventColorsFinal -- Last confirmed colors for splice events.
    """
    # Select correct colorset depending on whih button was pressed last
    evColors = eventColorsFinal
    data = [] # Will hold all generated traces
    axisTitles = [] # Will hold axis titles to be added in the main callback
    eventMaxHeights = [] # Will hold number of stacked event rows for layouting
    legendSet = {} # Keeps track of legend items to avoid duplicates for event types
    for val in cfg.eventTypes:
        legendSet[val] = False
    legendSet['one'] = True
    covTime = 0
    for ds in sorted(displayed):
        if cfg.spliceAvail:
            covStart = time.time()
            data.append(createAreaChart(xVals, yVals, ds, colorDict, axisTitles))
            covEnd = time.time()
            covTime += covEnd-covStart
        if cfg.spliceEventAvail:
            data.append(createEventPlots(eventData, ds, axisTitles, eventMaxHeights, evColors, legendSet))
    return (data, eventMaxHeights, axisTitles)

def createAreaChart(xVals, yVals, ds, colorDict, axisTitles):
    """ Creates an area chart from provided values. axisTitles is modified by this function.
    
        Positional arguments:
        xVals -- X-axis values for the area chart.
        yVals -- Y-axis values for the area chart.
        ds -- Dataset this chart is for, used for color selection and naming.
        colorDict -- Dict holding colors by datasets.
        axisTitles -- List of y-axis titles.
    """
    xAxis = xVals[ds]
    yAxis = yVals[ds]
    organism = ds.split('_')[0]
    orgColor = colorDict[organism]
    trace = go.Scatter(
        x=xAxis,
        y=yAxis,
        name=ds,
        meta = ds,
        fill='tozeroy',
        fillcolor=orgColor,
        hoveron='points+fills',
        line=dict(color='black'),
        text=ds,
        hoverinfo='y',
        cliponaxis=True
    )
    axisTitles.append('')
    return trace

def createEventPlots(eventData, ds, axisTitles, eventMaxHeights, evColors, legendSet):
    """ This function is a wrapper for splice event plot creation. It calls the correct 
        subfunction depending on dosplayMode. The function modifies axisTitles,eventMaxHeights
        and legendSet
    
        Positional arguments:
        eventData -- The dataframe containing the splice event data.
        ds -- Name of the dataset plots should be created for.
        axistitles -- List that holds axistitles.
        eventMaxHeights -- List that contains the number of stacked event rows for each event trace.
        evColors -- Colors for the different splice event types.
        legendSet -- Keeps track of which legend items to show to avoid duplicates.
    """
    maxStack = 0 # keeps track of the maximum number of stacked bars, to avoid empty rows
    traceDict = {}
    for i in ['one', 'two']:
        intervals = [] # Used to calculate overlaps, stores used intervals as well as row that interval was put on
        eventXValues = {} # Stores x-axis values per event type
        eventWidths = {} # Stores widths per event type
        eventBases = {} # Stores y offset per event type
        eventScores = {}
        # Iterate through dataframe rows and calculate stacking aswell as bar parameters

        for row in eventData[ds].itertuples():
            maxStack = calculateEvents(row.type, row.chromStart, row.chromEnd, row.score, eventXValues, eventWidths, eventBases, eventScores, intervals, maxStack)
        # eventMaxHeights will be used to scale the size of event traces based on the number
        # of stacked event rows  
        traces = []
        for k in sorted(eventXValues.keys()):
            legend = False
            if i == 'one' and legendSet['one'] == True:
                legend = True
                legendSet['one'] = False
            if i == 'two':
                if legendSet[k] == False: # Legend item for this event type is not displayed, display it
                    legendSet[k] = True
                    legend = True
                else:
                    legend = False
            traceColor = 'darkblue'
            #legend = True
            if i == 'two':
                traceColor = evColors[k]
                legendGroup = k
                traceName = k
            else:
                legendGroup = 'eventRegions'
                traceName = 'event regions'
            trace = go.Bar(
               # text = eventScores,
                #hoverinfo = 'x+text',
                x=eventXValues[k],
                y=[1]*len(eventXValues[k]),
                width = eventWidths[k],
                base = eventBases[k],
                name = traceName,
                meta = ds,
                showlegend = legend,
                legendgroup = legendGroup, # Group traces from different datasets so they all repsond to the one legend item
                insidetextfont=dict(
                    family="Arial",
                    color="black"
                ),
                text = eventScores[k],
                hoverinfo = 'x+text',
                marker=dict(
                    color= traceColor,
                )
            )
            traces.append(trace)
        if len(traces) > 0:
            traceDict.update({i : traces})
        else:
            traceDict.update({i : [ds]})
    else: # Displaymode: Score heatmap
        colorScale = (-1.0,1.0) # Scores range from -1 to 1, setup color scale for consistent coloring
        intervals = [] # Used to calculate overlaps, stores used intervals as well as row that interval was put on
        eventXValues = [] # Stores x-axis values per event type
        eventWidths = [] # Stores widths per event type
        eventBases = [] # Stores y offset per event type
        eventScores = [] # score for each event
        # Iterate through dataframe rows and calculate stacking aswell as bar parameters
       # maxStack = 0 # keeps track of the maximum number of stacked bars, to avoid empty rows
        for row in eventData[ds].itertuples():
            maxStack = calculateEventsScoreColored(row.chromStart, row.chromEnd, row.score, eventXValues, eventWidths, eventBases, eventScores, intervals, maxStack)
        # eventMaxHeights will be used to scale the size of event traces based on the number
        # of stacked event rows
        #eventMaxHeights.append(maxStack)    
        trace = go.Bar(
            x=eventXValues,
            y=[1]*len(eventXValues),
            width = eventWidths,
            base = eventBases,
            meta = ds,
            showlegend = False,
            insidetextfont=dict(
                family="Arial",
                color="black"
            ),
            text = eventScores,
            hoverinfo = 'x+text',
            marker=dict(
                color= eventScores,
                colorscale=[[0.25, "rgb(165,0,38)"],
                            [0.5, "rgb(0,0,0)"],
                            [0.75, "rgb(49,54,149)"]],
                showscale = True,
                cmin = colorScale[0],
                cmax = colorScale[1],
                colorbar =dict(
                    len = 1.0,
                    y = 0.0,
                    x = 1.0,
                    yanchor = 'bottom',
                )
            )
        )
        if len(eventXValues) > 0:
            traceDict.update({'three' : trace})
        else:
            traceDict.update({'three' : [ds]})
    axisTitles.append('')
    eventMaxHeights.append(maxStack)
    return traceDict

def calculateEvents(key, chromStart, chromEnd, score, eventXValues, eventWidths, eventBases, eventScores, intervals, maxStack, oneKey=False):
    """ This function calculates a bar and its vertical position for displayMode one and two. 
        It directly modifies eventXValues, eventWidths, eventBases, eventScores and intervals 
        and returns a new value for maxstack. 
        
        Positional arguments:
        key -- The type of splice event this event belongs to.
        chromstart -- Start of the event.
        chromEnd -- End of the event.
        score -- Score of the event.
        eventXValues -- Holds x-axis center coordinates of all events that have been processed already.
        eventWidths -- Holds widths of all processed events.
        eventBases -- Hold the vertical offset of all processed events.
        eventScores -- Holds the scores of all processed events.
        intervals -- Stores events as tuple containing start and end positions as well as vertical offset.
        maxStack -- Keeps track of the currently highest event row for layouting purposes.
    """
    # Avoid problems with events that don't follow the convention of chromStart < chromEnd
    maxVal = max(chromStart, chromEnd)
    minVal = min(chromStart, chromEnd)
    if len(intervals) == 0: # Row is the first row, no comparisons
        try: # If list already exist append
            eventXValues[key].append(minVal + ((maxVal-1) - minVal) / 2)
            eventWidths[key].append(maxVal - minVal)
            eventBases[key].append(0)
            eventScores[key].append(score)
            intervals.append(((minVal, maxVal),0))
        except KeyError: # Else create corresponding lists in dictionary
            eventXValues[key] = [minVal + ((maxVal-1) - minVal) / 2]
            eventWidths[key] = [maxVal - minVal]
            eventBases[key] = [0]
            eventScores[key] = [score]
            intervals.append(((minVal, maxVal),0))
        maxStack = 1
    else: # Row is not the first row, check through already processed intervals to calculate offset
        numOverlaps = 0
        heights = [] # Store all rows on which overlaps occur
        for i in intervals:
            if overlap(i[0], (minVal, maxVal)) == True:
                heights.append(i[1])
                numOverlaps += 1
        if len(heights) > 0:
            slot = 0 # First free row the new bar can be placed on
            for value in range(0, max(heights)+2): # Find first open slot
                if value not in heights:
                    slot = value
                    break
            numOverlaps = slot # Set numOverlaps accordingly
        try: # Try to append values to the list
            eventXValues[key].append(minVal + ((maxVal-1) - minVal) / 2)
            eventWidths[key].append(maxVal - minVal)
            if numOverlaps >= maxStack: # Check if a new row needs to be created to place this event
                if numOverlaps > maxStack + 1:
                    maxStack += 1
                    numOverlaps = maxStack
                else:
                    maxStack = numOverlaps + 1
            eventBases[key].append(numOverlaps + 0.5*numOverlaps)
            eventScores[key].append(score)
            intervals.append(((minVal, maxVal),numOverlaps))
        except KeyError: # Create the list corresponding to key
            eventXValues[key]  = [minVal + ((maxVal-1) - minVal) / 2]
            eventWidths[key] = [maxVal - minVal]
            if numOverlaps >= maxStack: # Check if a new row needs to be created to place this event
                if numOverlaps > maxStack + 1:
                    maxStack += 1
                    numOverlaps = maxStack
                else:
                    maxStack = numOverlaps + 1
            eventBases[key] = [numOverlaps + 0.5*numOverlaps]
            eventScores[key] = [score]
            intervals.append(((minVal, maxVal), numOverlaps))
    return maxStack

def calculateEventsScoreColored(chromStart, chromEnd, score, eventXValues, eventWidths, eventBases, eventScores, intervals, maxStack):
    """ This function calculates a bar and its vertical position for displayMode three. 
        It directly modifies eventXValues, eventWidths, eventBases, eventScores and intervals 
        and returns a new value for maxstack. 
    
        Positional arguments:
        chromstart -- Start of the event.
        chromEnd -- End of the event.
        score -- Score of the event.
        eventXValues -- Holds x-axis center coordinates of all events that have been processed already.
        eventWidths -- Holds widths of all processed events.
        eventBases -- Hold the vertical offset of all processed events.
        eventScores -- Holds the scores of all processed events.
        intervals -- Stores events as tuple containing start and end positions as well as vertical offset.
        maxStack -- Keeps track of the currently highest event row for layouting purposes.
    """
    maxVal = max(chromStart, chromEnd) 
    minVal = min(chromStart, chromEnd)
    if len(intervals) == 0: # Row is the first row, no comparisons
        eventXValues.append(minVal + ((maxVal-1) - minVal) / 2)
        eventWidths.append(maxVal - minVal)
        eventBases.append(0)
        eventScores.append(score)
        intervals.append(((minVal, maxVal),0))
        maxStack = 1
    else: # Row is not the first row, check through already processed intervals to calculate offset
        numOverlaps = 0
        heights = [] # Store all rows on which overlaps occur
        for i in intervals:
            if overlap(i[0], (minVal, maxVal)) == True:
                heights.append(i[1])
                numOverlaps += 1
        if len(heights) > 0:
            slot = 0 # First free row the new bar can be placed on
            for value in range(0, max(heights)+2): # Find first open slot
                if value not in heights:
                    slot = value
                    break
            numOverlaps = slot # Set numOverlaps accordingly
        eventXValues.append(minVal + ((maxVal-1) - minVal) / 2)
        eventWidths.append(maxVal - minVal)
        if numOverlaps >= maxStack:
            if numOverlaps > maxStack + 1: # Check if a new row needs to be created to place this event
                maxStack += 1
                numOverlaps = maxStack
            else:
                maxStack = numOverlaps + 1
        eventBases.append(numOverlaps + 0.5*numOverlaps)
        eventScores.append(score)
        intervals.append(((minVal, maxVal),numOverlaps))
    return maxStack
```
