## Additional File 1 for "SEQing: web-based visualization of iCLIP and RNA-seq data in an interactive python framework": settings_tab.py

```
#!/usr/bin/env python3
# -*- coding: utf-8 -*-

from app import app
import dash
import dash_html_components as html
import cfg

@app.callback(
    dash.dependencies.Output('bsGraph', 'config'),
    [dash.dependencies.Input('imgFormatDrop', 'value')]
)
def changeFormatiCLIP(imgFormat):
    """ Changes the image format for the iCLIP plot.
    
    Positional arguments:
    imgFormat -- Image format to use.
    """
    return {'toImageButtonOptions' : {'filename' : 'iCLIP', 'width' : None,
                'scale' : 1.0, 'height' : None, 'format' : imgFormat} }
          
@app.callback(
    dash.dependencies.Output('spliceGraph', 'config'),
    [dash.dependencies.Input('imgFormatDrop', 'value')]
)
def changeFormatRNA(imgFormat):
    """ Changes the image format for the iCLIP plot
    
    Positional arguments:
    imgFormat -- Image format to use.
    """
    return {'toImageButtonOptions' : {'filename' : 'RNA', 'width' : None,
                'scale' : 1.0, 'height' : None, 'format' : imgFormat} }
                                     
@app.callback(
    dash.dependencies.Output('legendSpacingDiv', 'data'),
    [dash.dependencies.Input('legendSpacingSlider', 'value')]
)
def changeLegendSpacing(value):
    return value


@app.callback(
    dash.dependencies.Output('rDisp', component_property='children'),
    [dash.dependencies.Input('rInput', component_property='value')]
)
def showR(r):
    """Callback to display current value for red.

    Positional arguments:
    r -- Value for red.
    """
    return html.P(html.B('R: ' + str(r)))


@app.callback(
    dash.dependencies.Output('gDisp', component_property='children'),
    [dash.dependencies.Input('gInput', component_property='value')]
)
def showG(g):
    """Callback to display current value for green

    Positional arguments:
    g -- Value for green.
    """

    return html.P(html.B('G: ' + str(g)))


@app.callback(
    dash.dependencies.Output('bDisp', component_property='children'),
    [dash.dependencies.Input('bInput', component_property='value')]
)
def showB(b):
    """Callback to display current value for blue.

    Positional arguments:
    b -- Value for blue.
    """

    return html.P(html.B('B: ' + str(b)))


@app.callback(
    dash.dependencies.Output('preview', component_property='style'),
    [dash.dependencies.Input('rInput', 'value'),
     dash.dependencies.Input('gInput', 'value'),
     dash.dependencies.Input('bInput', 'value')]
)
def previewColor(r, g, b):
    """Callback for rgb color preview.
    Positional arguments:
    r -- Value for red.
    g -- Value for green.
    b -- Value for blue.
    """

    if r == None or b == None or g == None:
        return {'backgroundColor': 'rgb(255, 255, 255)', 'color': 'rgb(255, 255, 255)'}
    else:
        return {'backgroundColor': 'rgb(' + str(r) + ',' + str(g) + ','
                                   + str(b) + ')', 'color': 'rgb(' + str(r)
                                                            + ',' + str(g) + ',' + str(b) + ')'}


@app.callback(
    dash.dependencies.Output('rInput', component_property='value'),
    [dash.dependencies.Input('colorDrop', 'value'),
     dash.dependencies.Input('geneDrop', 'value')],
    [dash.dependencies.State('colorFinal', 'data')]
)
def rCallback(dataset, geneid, colors):
    """Callback to set initial value of red slider from dict.

    Positional arguments:.
    dataset -- Currently selected dataset.
    geneid -- not needed, only to register input.
    colors -- Dictionary containing the color values.
    """

    colorsDict = colors
    try:
        colorVal = colorsDict[dataset][4:-1].split(',')[0]
        return int(colorVal)
    except KeyError:
        return 0


@app.callback(
    dash.dependencies.Output('gInput', component_property='value'),
    [dash.dependencies.Input('colorDrop', 'value'),
     dash.dependencies.Input('geneDrop', 'value')],
    [dash.dependencies.State('colorFinal', 'data')]
)
def gCallback(dataset, geneid, colors):
    """Callback to set initial value of green slider from dict.

    Positional arguments:
    dataset -- Currently selected dataset.
    geneid -- Not needed, only to register input.
    colors -- Dictionary containing the color values.
    """
    colorsDict = colors
    try:
        colorVal = colorsDict[dataset][4:-1].split(',')[1]
        return int(colorVal)
    except KeyError:
        return 0


@app.callback(
    dash.dependencies.Output('bInput', component_property='value'),
    [dash.dependencies.Input('colorDrop', 'value'),
     dash.dependencies.Input('geneDrop', 'value')],
    [dash.dependencies.State('colorFinal', 'data')]
)
def bCallback(dataset, geneid, colors):
    """Callback to set initial value of blue slider from dict.

    Positional arguments:
    dataset -- Currently selected dataset.
    geneid -- Not needed, only to register input
    colors -- Dictionary containing the color values.
    """
    colorsDict = colors
    try:
        colorVal = colorsDict[dataset][4:-1].split(',')[2]
        return int(colorVal)
    except KeyError:
        return 0


@app.callback(
    dash.dependencies.Output('colorFinal', component_property='data'),
    [dash.dependencies.Input('colorConfirm', 'n_clicks')],
    [dash.dependencies.State('rInput', 'value'),
     dash.dependencies.State('gInput', 'value'),
     dash.dependencies.State('bInput', 'value'),
     dash.dependencies.State('colorDrop', 'value'),
     dash.dependencies.State('colorFinal', 'data')]
)
def confirmColor(nclicks, r, g, b, dataset, backup):
    """ Callback to confirm a color. This will overwrite the previous one.

    Positional arguments:
    nclicks -- Button value.
    r -- Red value.
    g -- Green value.
    b -- Blue value.
    dataset -- Dataset to overwrite color of.
    backup -- Previous value in case of error.
    """
    if r == None or b == None or g == None:
        return backup
    else:
        colorDict = backup
        colorString = 'rgb(' + str(r) + ', ' + str(g) + ', ' + str(b) + ')'
        colorDict.update({dataset: colorString})
        return colorDict


@app.callback(
    dash.dependencies.Output('colorDiv', component_property='data'),
    [dash.dependencies.Input('rInput', 'value'),
     dash.dependencies.Input('gInput', 'value'),
     dash.dependencies.Input('bInput', 'value')],
    [dash.dependencies.State('colorDrop', 'value'),
     dash.dependencies.State('colorDiv', 'data')]
)
def changeColor(r, g, b, dataset, oldColors):
    """Callback to set new color values.

    Positional arguments:
    r -- Red value.
    g -- Green value.
    b -- Blue value.
    dataset -- Currently selected dataset.
    oldColors -- Previous colors in case none values are provided for r/g/b.
    """
    if r == None or b == None or g == None:
        return oldColors
    else:
        colorDict = oldColors
        colorString = 'rgb(' + str(r) + ', ' + str(g) + ', ' + str(b) + ')'
        colorDict.update({dataset: colorString})
        return colorDict

    
@app.callback(
    dash.dependencies.Output('covRDisp', component_property='children'),
    [dash.dependencies.Input('covRInput', component_property='value')]
)
def showRCov(r):
    """Callback to display current value for red.

    Positional arguments:
    r -- Value for red.
    """

    return html.P(html.B('R: ' + str(r)))


@app.callback(
    dash.dependencies.Output('covGDisp', component_property='children'),
    [dash.dependencies.Input('covGInput', component_property='value')]
)
def showGCov(g):
    """Callback to display current value for green.

    Positional arguments:
    g -- Value for green.
    """

    return html.P(html.B('G: ' + str(g)))


@app.callback(
    dash.dependencies.Output('covBDisp', component_property='children'),
    [dash.dependencies.Input('covBInput', component_property='value')]
)
def showBCov(b):
    """Callback to display current value for blue.

    Positional arguments:
    b -- Value for blue.
    """

    return html.P(html.B('B: ' + str(b)))


@app.callback(
    dash.dependencies.Output('covPreview', component_property='style'),
    [dash.dependencies.Input('covRInput', 'value'),
     dash.dependencies.Input('covGInput', 'value'),
     dash.dependencies.Input('covBInput', 'value')]
)
def previewColorCov(r, g, b):
    """Callback for rgb color preview.

    Positional arguments:
    r -- Value for red.
    g -- Value for green.
    b -- Value for blue.
    """

    if r == None or b == None or g == None:
        return {'backgroundColor': 'rgb(255, 255, 255)', 'color': 'rgb(255, 255, 255)'}
    else:
        return {'backgroundColor': 'rgb(' + str(r) + ',' + str(g) + ','
                                   + str(b) + ')', 'color': 'rgb(' + str(r)
                                                            + ',' + str(g) + ',' + str(b) + ')'}


@app.callback(
    dash.dependencies.Output('covRInput', component_property='value'),
    [dash.dependencies.Input('covColorDrop', 'value')],
    [dash.dependencies.State('covColorFinal', 'data')]
)
def rCallbackCov(dataset, colors):
    """Callback to set initial value of red slider from dict.

    Positional arguments:
    dataset -- Currently selected dataset.
    colors -- Dictionary containing the color values.
    """

    colorsDict = colors
    try:
        colorVal = colorsDict[dataset][4:-1].split(',')[0]
        return int(colorVal)
    except KeyError:
        return 0


@app.callback(
    dash.dependencies.Output('covGInput', component_property='value'),
    [dash.dependencies.Input('covColorDrop', 'value')],
    [dash.dependencies.State('covColorFinal', 'data')]
)
def gCallbackCov(dataset, colors):
    """Callback to set initial value of green slider from dict.

    Positional arguments:
    dataset -- Currently selected dataset.
    colors -- Dictionary containing the color values.
    """
    colorsDict = colors
    try:
        colorVal = colorsDict[dataset][4:-1].split(',')[1]
        return int(colorVal)
    except KeyError:
        return 0


@app.callback(
    dash.dependencies.Output('covBInput', component_property='value'),
    [dash.dependencies.Input('covColorDrop', 'value')],
    [dash.dependencies.State('covColorFinal', 'data')]
)
def bCallbackCov(dataset, colors):
    """Callback to set initial value of blue slider from dict.

    Positional arguments:
    dataset -- Currently selected dataset.
    colors -- Dictionary containing the color values.
    """
    colorsDict = colors
    try:
        colorVal = colorsDict[dataset][4:-1].split(',')[2]
        return int(colorVal)
    except KeyError:
        return 0


@app.callback(
    dash.dependencies.Output('covColorFinal', component_property='data'),
    [dash.dependencies.Input('covColorConfirm', 'n_clicks')],
    [dash.dependencies.State('covRInput', 'value'),
     dash.dependencies.State('covGInput', 'value'),
     dash.dependencies.State('covBInput', 'value'),
     dash.dependencies.State('covColorDrop', 'value'),
     dash.dependencies.State('covColorFinal', 'data')]
)
def conFirmColorCov(nclicks, r, g, b, dataset, backup):
    """ Callback to confirm a color. This will overwrite the previous one.

    Positional arguments:
    nclicks -- Button value.
    r -- Red value.
    g -- Green value.
    b -- Blue value.
    dataset -- Dataset to overwrite color of.
    backup -- Previous value in case of error.
    """
    if r == None or b == None or g == None:
        return backup
    else:
        colorDict = backup
        colorString = 'rgb(' + str(r) + ', ' + str(g) + ', ' + str(b) + ')'
        colorDict.update({dataset: colorString})
        return colorDict


@app.callback(
    dash.dependencies.Output('covColorDiv', component_property='data'),
    [dash.dependencies.Input('covRInput', 'value'),
     dash.dependencies.Input('covGInput', 'value'),
     dash.dependencies.Input('covBInput', 'value')],
    [dash.dependencies.State('covColorDrop', 'value'),
     dash.dependencies.State('covColorDiv', 'data')]
)
def changeColorCov(r, g, b, dataset, oldColors):
    """Callback to set new color values.

    Positional arguments:
    r -- Red value.
    g -- Green value.
    b -- Blue value.
    dataset -- Currently selected dataset.
    oldColors -- Previous colors in case none values are provided for r/g/b.
    """
    if r == None or b == None or g == None:
        return oldColors
    else:
        colorDict = oldColors
        colorString = 'rgb(' + str(r) + ', ' + str(g) + ', ' + str(b) + ')'
        colorDict.update({dataset: colorString})
        return colorDict

@app.callback(
    dash.dependencies.Output('eventRDisp', component_property='children'),
    [dash.dependencies.Input('eventRInput', component_property='value')]
)
def showREvent(r):
    """Callback to display current value for red.

    Positional arguments:
    r -- Value for red.
    """

    return html.P(html.B('R: ' + str(r)))


@app.callback(
    dash.dependencies.Output('eventGDisp', component_property='children'),
    [dash.dependencies.Input('eventGInput', component_property='value')]
)
def showEvent(g):
    """Callback to display current value for green.

    Positional arguments:
    g -- Value for green.
    """

    return html.P(html.B('G: ' + str(g)))


@app.callback(
    dash.dependencies.Output('eventBDisp', component_property='children'),
    [dash.dependencies.Input('eventBInput', component_property='value')]
)
def showBevent(b):
    """Callback to display current value for blue.

    Positional arguments:
    b -- Value for blue.
    """

    return html.P(html.B('B: ' + str(b)))


@app.callback(
    dash.dependencies.Output('eventPreview', component_property='style'),
    [dash.dependencies.Input('eventRInput', 'value'),
     dash.dependencies.Input('eventGInput', 'value'),
     dash.dependencies.Input('eventBInput', 'value')]
)
def previewColorEvent(r, g, b):
    """Callback for rgb color preview.

    Positional arguments:
    r -- Value for red.
    g -- Value for green.
    b -- Value for blue.
    """

    if r == None or b == None or g == None:
        return {'backgroundColor': 'rgb(255, 255, 255)', 'color': 'rgb(255, 255, 255)'}
    else:
        return {'backgroundColor': 'rgb(' + str(r) + ',' + str(g) + ','
                                   + str(b) + ')', 'color': 'rgb(' + str(r)
                                                            + ',' + str(g) + ',' + str(b) + ')'}


@app.callback(
    dash.dependencies.Output('eventRInput', component_property='value'),
    [dash.dependencies.Input('eventColorDrop', 'value')],
    [dash.dependencies.State('eventColorFinal', 'data')]
)
def rCallbackEvent(dataset, colors):
    """Callback to set initial value of red slider from dict.

    Positional arguments:
    dataset -- Currently selected dataset.
    colors -- Dictionary containing the color values.
    """

    colorsDict = colors
    try:
        colorVal = colorsDict[dataset][4:-1].split(',')[0]
        return int(colorVal)
    except KeyError:
        return 0


@app.callback(
    dash.dependencies.Output('eventGInput', component_property='value'),
    [dash.dependencies.Input('eventColorDrop', 'value')],
    [dash.dependencies.State('eventColorFinal', 'data')]
)
def gCallbackEvent(dataset, colors):
    """Callback to set initial value of green slider from dict.

    Positional arguments:
    dataset -- Currently selected dataset.
    colors -- Dictionary containing the color values.
    """
    colorsDict = colors
    try:
        colorVal = colorsDict[dataset][4:-1].split(',')[1]
        return int(colorVal)
    except KeyError:
        return 0


@app.callback(
    dash.dependencies.Output('eventBInput', component_property='value'),
    [dash.dependencies.Input('eventColorDrop', 'value')],
    [dash.dependencies.State('eventColorFinal', 'data')]
)
def bCallbackEvent(dataset, colors):
    """Callback to set initial value of blue slider from dict.

    Positional arguments:
    dataset -- Currently selected dataset.
    colors -- Dictionary containing the color values.
    """
    colorsDict = colors
    try:
        colorVal = colorsDict[dataset][4:-1].split(',')[2]
        return int(colorVal)
    except KeyError:
        return 0


@app.callback(
    dash.dependencies.Output('eventColorFinal', component_property='data'),
    [dash.dependencies.Input('eventColorConfirm', 'n_clicks')],
    [dash.dependencies.State('eventRInput', 'value'),
     dash.dependencies.State('eventGInput', 'value'),
     dash.dependencies.State('eventBInput', 'value'),
     dash.dependencies.State('eventColorDrop', 'value'),
     dash.dependencies.State('eventColorFinal', 'data')]
)
def conFirmColorEvent(nclicks, r, g, b, dataset, backup):
    """ Callback to confirm a color. This will overwrite the previous one.

    Positional arguments:
    nclicks -- Button value.
    r -- Red value.
    g -- Green value.
    b -- Blue value.
    dataset -- Dataset to overwrite color of.
    backup -- Previous value in case of error.
    """
    if r == None or b == None or g == None:
        return backup
    else:
        colorDict = backup
        colorString = 'rgb(' + str(r) + ', ' + str(g) + ', ' + str(b) + ')'
        colorDict.update({dataset: colorString})
        return colorDict


@app.callback(
    dash.dependencies.Output('eventColorDiv', component_property='data'),
    [dash.dependencies.Input('eventRInput', 'value'),
     dash.dependencies.Input('eventGInput', 'value'),
     dash.dependencies.Input('eventBInput', 'value')],
    [dash.dependencies.State('eventColorDrop', 'value'),
     dash.dependencies.State('eventColorDiv', 'data')]
)
def changeColorEvent(r, g, b, dataset, oldColors):
    """Callback to set new color values.

    Positional arguments:
    r -- Red value.
    g -- Green value.
    b -- Blue value.
    dataset -- Currently selected dataset.
    oldColors -- Previous colors in case none values are provided for r/g/b.
    """
    if r == None or b == None or g == None:
        return oldColors
    else:
        colorDict = oldColors
        colorString = 'rgb(' + str(r) + ', ' + str(g) + ', ' + str(b) + ')'
        colorDict.update({dataset: colorString})
        return colorDict


@app.callback(
    dash.dependencies.Output('headline', component_property='children'),
    [dash.dependencies.Input('geneDrop', 'value')]
)
def setHeadline(name):
    """Callback to set the headline.

    Positional arguments:
    clicks -- Related to button, not needed otherwise.
    name -- Name of the currently selected gene.
    """
    for i in cfg.geneAnnotations:
        currentGene = i[i['geneID'].str.contains(name)]
        if not currentGene.empty:
            break
    strand = currentGene['strand'].iloc[0]
    title = name + ' (' + strand + ')'
    return title
```
