## Additional File 1 for "SEQing: web-based visualization of iCLIP and RNA-seq data in an interactive python framework": unitTesting.py

```
#!/usr/bin/env python
# -*- coding: utf-8 -*-

"""Unit tests for SEQing"""
import unittest
import rna_tab as rna
import iclip_tab as iclip
import settings_tab as settings
import description_tab as details
import validator as val
import converter as conv
import json
import dash_html_components as html
from Bio.Alphabet import generic_dna
from Bio.Seq import Seq
from Bio.SeqRecord import SeqRecord
import collections
import pandas
import numpy as np

class TestConverter(unittest.TestCase):
    def testConvertGTFToBed(self):
        gtfHeader = ['seqname', 'source', 'feature', 'start', 'end', 'score',
               'strand', 'frame', 'attribute']
        bedHeader = ['chrom','chromStart','chromEnd','name','score','strand','thickStart',
             'thickEnd','itemRGB','blockCount','blockSizes','blockStarts']
        dtypes = {'seqname' : 'object', 'source' : 'object', 'feature' : 'object', 'start' : 'uint32', 'end': 'uint32', 'score' : 'object',
                              'strand' : 'category', 'frame' : 'object', 'attribute' : 'object'}
        dtypesBed12 = {'chrom' : 'category', 'chromStart' : 'uint32','chromEnd': 'uint32', 'name' : 'object', 'score' : 'int16', 'strand' : 'category','thickStart' : 'uint64',
                 'thickEnd' : 'uint64', 'blockCount' : 'uint32','itemRGB' : 'int16','blockSizes' : 'object','blockStarts' : 'object'}
        testCases = []
        # Case that should work
        bedFile = []
        bedFile.append(['Chr1', 'Araport11', '5UTR', 3631, 3759, '.', '+', '.', 'transcript_id "AT1G01010.1"; gene_id "AT1G01010";'])
        bedFile.append(['Chr1', 'Araport11', 'exon', 3631, 3913, '.', '+', '.', 'transcript_id "AT1G01010.1"; gene_id "AT1G01010";'])
        bedFile.append(['Chr1', 'Araport11', 'CDS', 3760, 3913, '.', '+', '.', 'transcript_id "AT1G01010.1"; gene_id "AT1G01010";'])
        bedFile.append(['Chr1', 'Araport11', '5UTR', 3631, 3759, '.', '+', '.', 'transcript_id "AT1G01010.1"; gene_id "AT1G01010";'])    
        df = pandas.DataFrame(data = bedFile, columns = gtfHeader)
        for key, dtype in dtypes.items():
            df[key] = df[key].astype(dtype)
        outFile = []
        outFile.append(['Chr1',3630,3913,'AT1G01010.1',0,'+',3759,3913,0,1,'283','0'])
        outDF = pandas.DataFrame(data = outFile, columns = bedHeader)
        for key, dtype in dtypesBed12.items():
            outDF[key] = outDF[key].astype(dtype)        
        testCases.append((df,outDF))
        for i in testCases:
            self.assertTrue(conv.convertGTFToBed(i[0]).equals(i[1]))
            
class TestValidator(unittest.TestCase):
    def testISRGB(self):
        testCases = []
        testCases.append((None, False))
        testCases.append((22, False))
        testCases.append(([2,3,4,5], False))
        testCases.append(('', False))
        testCases.append((' ', False))
        testCases.append(('rgb(22,22,22)', True))
        testCases.append(('rgb(-22,22,22)', False))
        testCases.append(('rgb(22,-22,22)', False))
        testCases.append(('rgb(22,22,-22)', False))
        testCases.append(('rgb(2222,22,22)', False))
        testCases.append(('rgb(22,2222,22)', False))
        testCases.append(('rgb(22,22,2222)', False))
        for i in testCases:
            if i[1] == False:
                self.assertFalse(val.isRGB(i[0]))
            else:
                self.assertTrue(val.isRGB(i[0]))              

    def testValidateGTF(self):
        gtfHeader = ['seqname', 'source', 'feature', 'start', 'end', 'score',
               'strand', 'frame', 'attribute']
        dtypes = {'seqname' : 'object', 'source' : 'object', 'feature' : 'object', 'start' : 'uint32', 'end': 'uint32', 'score' : 'object',
                              'strand' : 'category', 'frame' : 'object', 'attribute' : 'object'}
        testCases = []
        # Case that should work
        bedFile = []
        bedFile.append(['test.1', 'source', 'gene' , 300, 500, '.', '-', '.', '.'])
        bedFile.append(['test.2', 'source', 'gene' , 600, 700, '.', '+', '.', '.'])
        df = pandas.DataFrame(data = bedFile, columns = gtfHeader)
        for key, dtype in dtypes.items():
            df[key] = df[key].astype(dtype)
        testCases.append((df,(True,'')))
        # Case with only a single strand symbol present
        bedFile = []
        bedFile.append(['test.1', 'source', 'gene' , 300, 500, '.', '-', '.', '.'])
        bedFile.append(['test.2', 'source', 'gene' , 600, 700, '.', '-', '.', '.'])
        df = pandas.DataFrame(data = bedFile, columns = gtfHeader)
        for key, dtype in dtypes.items():
            df[key] = df[key].astype(dtype)
        testCases.append((df,(True,'')))
        # Case that should fail on missing values
        bedFile = []
        bedFile.append(['test.1', 'source', 'gene' , 300, 500, np.NaN, '-', '.', '.'])
        bedFile.append([np.NaN, 'source', 'gene' , 600, 799, '.', '-', '.', '.'])
        df = pandas.DataFrame(data = bedFile, columns = gtfHeader)
        for key, dtype in dtypes.items():
            df[key] = df[key].astype(dtype)
        testCases.append((df,(False,'Missing values')))
        # Case that should fail on bad strand symbol
        bedFile = []
        bedFile.append(['test.1', 'source', 'gene' , 300, 500, '.', '-', '.', '.'])
        bedFile.append(['test.2', 'source', 'gene' , 600, 799, '.', ',', '.', '.'])
        df = pandas.DataFrame(data = bedFile, columns = gtfHeader)
        for key, dtype in dtypes.items():
            df[key] = df[key].astype(dtype)
        testCases.append((df,(False,'Bad strand symbol')))
        for i in testCases:
            result = val.validateGTF(i[0])
            self.assertEqual(result[0], i[1][0])
            self.assertEqual(result[1][0:13], i[1][1][0:13])
        self.assertFalse(val.validateGTF(None)[0])
        self.assertFalse(val.validateGTF(1)[0])
        self.assertFalse(val.validateGTF("2222")[0])
        self.assertFalse(val.validateGTF(pandas.DataFrame())[0])

    def testValidateBed12(self):
        bedHeader = ['chrom','chromStart','chromEnd','name','score','strand','thickStart',
             'thickEnd','itemRGB','blockCount','blockSizes','blockStarts']
        dtypes = {'chrom' : 'category', 'chromStart' : 'uint32','chromEnd': 'uint32', 'name' : 'object', 'score' : 'int16', 'strand' : 'category','thickStart' : 'uint64',
                 'thickEnd' : 'uint64', 'blockCount' : 'uint32','blockSizes' : 'object','blockStarts' : 'object'}
        testCases = []
        # Case that should work
        bedFile = []
        bedFile.append(['chrom1', 0, 10 , 'test.1', 0, '-', 0, 10, 0, 2, '2,4','0,3'])
        bedFile.append(['chrom1', 20, 40 , 'test.1', 0, '+', 1, 10, 0, 2,'2,3','20,23'])
        df = pandas.DataFrame(data = bedFile, columns = bedHeader)
        for key, dtype in dtypes.items():
            df[key] = df[key].astype(dtype)
        testCases.append((df,(True,'')))
        # Case with only a single strand symbol present
        bedFile = []
        bedFile.append(['chrom1', 0, 10 , 'test.1', 0, '-', 0, 10, 0, 2, '2,4','0,3'])
        bedFile.append(['chrom1', 20, 40 , 'test.1', 0, '-', 1, 10, 0, 2,'2,3','20,23'])
        df = pandas.DataFrame(data = bedFile, columns = bedHeader)
        for key, dtype in dtypes.items():
            df[key] = df[key].astype(dtype)
        testCases.append((df,(True,'')))
        # Case that should fail on missing values
        bedFile = []
        bedFile.append(['chrom1', 0, 10 , np.NaN, 0, '-', 0, 10, 0, 2, '2,4','0,3'])
        bedFile.append(['chrom1', 20, 40 , 'test.1', 0, '-', 1, 10, 0, 2,'2,3','20,23'])
        df = pandas.DataFrame(data = bedFile, columns = bedHeader)
        for key, dtype in dtypes.items():
            df[key] = df[key].astype(dtype)
        testCases.append((df,(False,'Missing values')))
        # Case that should fail on bad strand symbol
        bedFile = []
        bedFile.append(['chrom1', 0, 10 , 'test.1', 0, 'g', 0, 10, 0, 2, '2,4','0,3'])
        bedFile.append(['chrom1', 20, 40 , 'test.1', 0, '-', 1, 10, 0, 2,'2,3','20,23'])
        df = pandas.DataFrame(data = bedFile, columns = bedHeader)
        for key, dtype in dtypes.items():
            df[key] = df[key].astype(dtype)
        testCases.append((df,(False,'Bad strand symbol')))
        for i in testCases:
            result = val.validateBed12(i[0])
            self.assertEqual(result[0], i[1][0])
            self.assertEqual(result[1][0:13], i[1][1][0:13])
        self.assertFalse(val.validateGTF(None)[0])
        self.assertFalse(val.validateBed12(1)[0])
        self.assertFalse(val.validateBed12("2222")[0])
        self.assertFalse(val.validateBed12(pandas.DataFrame())[0])

    def testValidateBedGraph(self):
        rawHeader = ['chrom','chromStart','chromEnd','count']
        dtypes = {'chrom' : 'category' ,'chromStart' : 'uint64','chromEnd' : 'uint64', 'count' : 'uint32'}
        testCases = []
        # Case that should work
        bedFile = []
        bedFile.append(['chrom1', 0, 10 , 0])
        bedFile.append(['chrom1', 20, 40 , 2])
        df = pandas.DataFrame(data = bedFile, columns = rawHeader)
        for key, dtype in dtypes.items():
            df[key] = df[key].astype(dtype)
        testCases.append((df,(True,'')))
        # Case that should fail on missing values
        bedFile = []
        bedFile.append([np.NaN, 0, 10 , 0])
        bedFile.append(['chrom1', 20, 40 , 2])
        df = pandas.DataFrame(data = bedFile, columns = rawHeader)
        for key, dtype in dtypes.items():
            df[key] = df[key].astype(dtype)
        testCases.append((df,(False,'Missing values')))
        for i in testCases:
            result = val.validateBedGraph(i[0])
            self.assertEqual(result[0], i[1][0])
            self.assertEqual(result[1][0:13], i[1][1][0:13])
        self.assertFalse(val.validateBedGraph(None)[0])
        self.assertFalse(val.validateBedGraph(1)[0])
        self.assertFalse(val.validateBedGraph("2222")[0])
        self.assertFalse(val.validateBedGraph(pandas.DataFrame())[0])

    def testValidateBed(self):
        bsHeader = ['chrom', 'chromStart','chromEnd','type', 'score', 'strand']
        dtypes = {'chrom' : 'category', 'chromStart' : 'uint64','chromEnd' : 'uint64','type' : 'category', 'score' : 'float32', 'strand' : 'category'}
        testCases = []
        # Case that should work
        bedFile = []
        bedFile.append(['chrom1', 0, 10 , '.', 0 , '-'])
        bedFile.append(['chrom1', 20, 40 , '.', 0, '+'])
        df = pandas.DataFrame(data = bedFile, columns = bsHeader)
        for key, dtype in dtypes.items():
            df[key] = df[key].astype(dtype)
        testCases.append((df,(True,'')))
        # Case with only a single strand symbol present
        bedFile = []
        bedFile.append(['chrom1', 0, 10 , '.', 0 , '-'])
        bedFile.append(['chrom1', 20, 40 , '.', 0, '-'])
        df = pandas.DataFrame(data = bedFile, columns = bsHeader)
        for key, dtype in dtypes.items():
            df[key] = df[key].astype(dtype)
        testCases.append((df,(True,'')))
        # Case that should fail on missing values
        bedFile = []
        bedFile.append([np.NaN, 0, 10 , '.', 0 , '-'])
        bedFile.append(['chrom1', 20, 40 , '.', 0, '-'])
        df = pandas.DataFrame(data = bedFile, columns = bsHeader)
        for key, dtype in dtypes.items():
            df[key] = df[key].astype(dtype)
        testCases.append((df,(False,'Missing values')))
        # Case that should fail on bad strand symbol
        bedFile = []
        bedFile.append(['chrom1', 0, 10 , '.', 0 , '-'])
        bedFile.append(['chrom1', 20, 40 , '.', 0, 'df'])
        df = pandas.DataFrame(data = bedFile, columns = bsHeader)
        for key, dtype in dtypes.items():
            df[key] = df[key].astype(dtype)
        testCases.append((df,(False,'Bad strand symbol')))
        for i in testCases:
            result = val.validateBed(i[0])
            self.assertEqual(result[0], i[1][0])
            self.assertEqual(result[1][0:13], i[1][1][0:13])
        self.assertFalse(val.validateBed(None)[0])
        self.assertFalse(val.validateBed(1)[0])
        self.assertFalse(val.validateBed("2222")[0])
        self.assertFalse(val.validateBed(pandas.DataFrame())[0])        
        
class TestDashboard(unittest.TestCase):
    def testFormatChangeiCLIP(self):
        self.maxDiff = None
        null = None
        for test in ['svg', 'png']:
            self.assertEqual(eval(settings.changeFormatiCLIP(test)), 
                {'response': {'props': {'config':{'toImageButtonOptions' : {'filename' : 'iCLIP', 'width' : None,
                'scale' : 1.0, 'height' : None, 'format' : test} }}}})
  
    def testFormatChangeRNA(self):
        self.maxDiff = None
        null = None
        for test in ['svg', 'png']:
            self.assertEqual(eval(settings.changeFormatRNA(test)), 
                {'response': {'props': {'config':{'toImageButtonOptions' : {'filename' : 'RNA', 'width' : None,
                'scale' : 1.0, 'height' : None, 'format' : test} }}}})
    
    def testChangeLegendSpacing(self):
        self.maxDiff = None
        for test in [0, 0.1, 0.2, 0.3]:
            self. assertEqual(eval(settings.changeLegendSpacing(test)),  {'response': {'props': {'children':str(test)}}} )
          
    def testConirmColor(self):
        testCases = []
        testCases.append(('test', json.dumps({'test' :'rgb(0,0,0)'}), {'test' : 'rgb(23, 23, 23)'}, (23,23,23)))
        testCases.append(('test', json.dumps({'test' :'rgb(0,0,0)'}), {'test' :'rgb(0,0,0)'} , (None,23,23)))
        testCases.append(('test', json.dumps({'test' :'rgb(0,0,0)'}), {'test' :'rgb(0,0,0)'}, (23,None,23)))
        testCases.append(('test', json.dumps({'test' :'rgb(0,0,0)'}), {'test' :'rgb(0,0,0)'}, (23,23,None)))
        testCases.append(('set2', json.dumps({'set2' :'rgb(0,0,0)'}), {'set2' : 'rgb(244, 23, 23)'}, (244,23,23)))
        for i in testCases:
            self.assertEqual(eval(settings.confirmColor(0, i[3][0], i[3][1], i[3][2], i[0], i[1])),{'response': {'props': {'children' : json.dumps(i[2])}}} ) 
    
    def testChangeColor(self):
        testCases = []
        testCases.append(('test',(22,34,134), json.dumps({'test' :'rgb(0,0,0)'}), {'test' :'rgb(22, 34, 134)'}))
        testCases.append(('test',(22,None,134), json.dumps({'test' :'rgb(0,0,0)'}), {'test' :'rgb(0,0,0)'}))
        testCases.append(('test',(None,34,134), json.dumps({'test' :'rgb(0,0,0)'}), {'test' :'rgb(0,0,0)'}))
        testCases.append(('test',(22,34,None), json.dumps({'test' :'rgb(0,0,0)'}), {'test' :'rgb(0,0,0)'}))
        for i in testCases:
            self.assertEqual(eval(settings.changeColor(i[1][0], i[1][1], i[1][2], i[0], i[2])),{'response': {'props': {'children' : json.dumps(i[3])}}})
    
    def testPreviewColorEvent(self):
        self.maxDiff = None
        self.assertEqual(eval(settings.previewColorEvent(23, None, 66)),
            {'response': {'props': {'style': {'backgroundColor': 'rgb(255, 255, 255)', 'color': 'rgb(255, 255, 255)'}}}})
        self.assertEqual(eval(settings.previewColorEvent(23, 48, 66)), 
            {'response': {'props': {'style': {'backgroundColor': 'rgb(23,48,66)', 'color': 'rgb(23,48,66)'}}}})
        
    def testOverlap(self):
        self.assertTrue(rna.overlap((1,4),(2,7)))
        self.assertTrue(rna.overlap((1,4),(1,4)))
        self.assertFalse(rna.overlap((1,4),(1,1)))
        self.assertFalse(rna.overlap((1,4),(5,7)))
        self.assertFalse(rna.overlap((1,4),(4,7)))
    
    def testGenerateMasterSequence(self):
        testCases = []
        records = collections.OrderedDict()
        records['test.1'] = SeqRecord(Seq('ATTTA', generic_dna), id ='test.1::Chr1:1-6', name = 'test.1::Chr1:1-6',
                           description = 'test.1::Chr1:1-6')
        records['test.2'] = SeqRecord(Seq('GCGCGC', generic_dna), id ='test.2::Chr1:6-12', name = 'test.2::Chr1:6-12',
                           description = 'test.2::Chr1:6-12'
                           )
        isoforms = [(1, 6, 'test.1'),(6, 12, 'test.2')]
        caseInput = ([records], isoforms, 12)
        caseOutput = 'ATTTAGCGCGC'
        testCases.append((caseInput, caseOutput))
        
        records = collections.OrderedDict()
        records['test.1'] = SeqRecord(Seq('AT', generic_dna), id ='test.1::Chr1:1-6', name = 'test.1::Chr1:1-6',
                           description = 'test.1::Chr1:1-6')
        records['test.2'] = SeqRecord(Seq('AAAAGCGGGG', generic_dna), id ='test.2::Chr1:6-12', name = 'test.2::Chr1:6-12',
                           description = 'test.2::Chr1:6-12'
                           )
        isoforms = [(1, 6, 'test.1'),(2, 12, 'test.2')]
        caseInput = ([records], isoforms, 12)
        caseOutput = ' AAAAGCGGGG'
        testCases.append((caseInput, caseOutput))
        # test more than 2 sequences
        records = collections.OrderedDict()
        records['test.1'] = SeqRecord(Seq('ATTTA', generic_dna), id ='test.1::Chr1:1-6', name = 'test.1::Chr1:1-6',
                           description = 'test.1::Chr1:1-6')
        records['test.2'] = SeqRecord(Seq('GCGCGC', generic_dna), id ='test.2::Chr1:6-12', name = 'test.2::Chr1:6-12',
                           description = 'test.2::Chr1:6-12'
                           )
        records['test.3'] = SeqRecord(Seq('TACTAC', generic_dna), id ='test.3::Chr1:12-18', name = 'test.2::Chr1:12-18',
                           description = 'test.2::Chr1:12-18'
                           )
        isoforms = [(1, 6, 'test.1'),(6, 11, 'test.2'),(12, 18, 'test.3')]
        caseInput = ([records], isoforms, 18)
        caseOutput = 'ATTTA      TACTAC'
        testCases.append((caseInput, caseOutput))
        # test simple gap insertion
        records = collections.OrderedDict()
        records['test.1'] = SeqRecord(Seq('ATTTA', generic_dna), id ='test.1::Chr1:1-6', name = 'test.1::Chr1:1-6',
                           description = 'test.1::Chr1:1-6')
        records['test.2'] = SeqRecord(Seq('GCGCGC', generic_dna), id ='test.2::Chr1:7-12', name = 'test.2::Chr1:7-12',
                           description = 'test.2::Chr1:7-12'
                           )
        isoforms = [(1, 6, 'test.1'),(7, 13, 'test.2')]
        caseInput = ([records], isoforms, 13)
        caseOutput = 'ATTTA GCGCGC'
        testCases.append((caseInput, caseOutput))
        # test double gaps
        records = collections.OrderedDict()
        records['test.1'] = SeqRecord(Seq('ATTTA', generic_dna), id ='test.1::Chr1:1-6', name = 'test.1::Chr1:1-6',
                           description = 'test.1::Chr1:1-6')
        records['test.2'] = SeqRecord(Seq('GCGCGC', generic_dna), id ='test.2::Chr1:6-12', name = 'test.2::Chr1:6-12',
                           description = 'test.2::Chr1:6-12'
                           )
        records['test.3'] = SeqRecord(Seq('TACTAC', generic_dna), id ='test.3::Chr1:12-18', name = 'test.2::Chr1:12-18',
                           description = 'test.2::Chr1:12-18'
                           )
        isoforms = [(1, 6, 'test.1'),(7, 13, 'test.2'),(14, 20, 'test.3')]
        caseInput = ([records], isoforms, 20)
        caseOutput = 'ATTTA GCGCGC TACTAC'
        testCases.append((caseInput, caseOutput))
        # Test with bad isoform name
        records = collections.OrderedDict()
        records['test.1'] = SeqRecord(Seq('ATTTA', generic_dna), id ='test.1::Chr1:1-6', name = 'test.1::Chr1:1-6',
                           description = 'test.1::Chr1:1-6')
        records['test.2'] = SeqRecord(Seq('GCGCGC', generic_dna), id ='test.2::Chr1:6-12', name = 'test.2::Chr1:6-12',
                           description = 'test.2::Chr1:6-12'
                           )
        isoforms = [(1, 6, 'test.1'),(6, 12, 'test.4')]
        caseInput = ([records], isoforms, 12)
        caseOutput = 'ATTTA      '
        testCases.append((caseInput, caseOutput))
        # Test with faulty name in between two working seqs
        records = collections.OrderedDict()
        records['test.1'] = SeqRecord(Seq('ATTTA', generic_dna), id ='test.1::Chr1:1-6', name = 'test.1::Chr1:1-6',
                           description = 'test.1::Chr1:1-6')
        records['test.2'] = SeqRecord(Seq('GCGCGC', generic_dna), id ='test.2::Chr1:6-12', name = 'test.2::Chr1:6-12',
                           description = 'test.2::Chr1:6-12'
                           )
        records['test.3'] = SeqRecord(Seq('TACTAC', generic_dna), id ='test.2::Chr1:6-12', name = 'test.2::Chr1:6-12',
                           description = 'test.2::Chr1:6-12'
                           )
        isoforms = [(1, 6, 'test.1'),(6, 12, 'test.4'),(12, 18, 'test.3')]
        caseInput = ([records], isoforms, 18)
        caseOutput = 'ATTTA      TACTAC'
        testCases.append((caseInput, caseOutput))
        # Test with working seq overlapping faulty one
        records = collections.OrderedDict()
        records['test.1'] = SeqRecord(Seq('ATTTA', generic_dna), id ='test.1::Chr1:1-6', name = 'test.1::Chr1:1-6',
                           description = 'test.1::Chr1:1-6')
        records['test.2'] = SeqRecord(Seq('GCGCGC', generic_dna), id ='test.2::Chr1:6-12', name = 'test.2::Chr1:6-12',
                           description = 'test.2::Chr1:6-12'
                           )
        records['test.3'] = SeqRecord(Seq('TACTAC', generic_dna), id ='test.2::Chr1:6-12', name = 'test.2::Chr1:6-12',
                           description = 'test.2::Chr1:6-12'
                           )
        isoforms = [(1, 6, 'test.1'),(6, 12, 'test.4'),(10, 16, 'test.3')]
        caseInput = ([records], isoforms, 16)
        caseOutput = 'ATTTA    TACTAC'
        testCases.append((caseInput, caseOutput))
        # Test with sequence too short
        records = collections.OrderedDict()
        records['test.1'] = SeqRecord(Seq('AT', generic_dna), id ='test.1::Chr1:1-6', name = 'test.1::Chr1:1-6',
                           description = 'test.1::Chr1:1-6')
        records['test.2'] = SeqRecord(Seq('GCGCGC', generic_dna), id ='test.2::Chr1:6-12', name = 'test.2::Chr1:6-12',
                           description = 'test.2::Chr1:6-12'
                           )
        isoforms = [(1, 6, 'test.1'),(6, 12, 'test.2')]
        caseInput = ([records], isoforms, 12)
        caseOutput = '     GCGCGC'
        testCases.append((caseInput, caseOutput))
        # Overlap with errornous sequence
        records = collections.OrderedDict()
        records['test.1'] = SeqRecord(Seq('ATTTA', generic_dna), id ='test.1::Chr1:1-6', name = 'test.1::Chr1:1-6',
                           description = 'test.1::Chr1:1-6')
        records['test.2'] = SeqRecord(Seq('GC', generic_dna), id ='test.2::Chr1:6-12', name = 'test.2::Chr1:6-12',
                           description = 'test.2::Chr1:6-12'
                           )
        isoforms = [(1, 6, 'test.1'),(6, 12, 'test.2')]
        caseInput = ([records], isoforms, 12)
        caseOutput = 'ATTTA      '
        testCases.append((caseInput, caseOutput))
        # Standalone with errornous Sequence
        records = collections.OrderedDict()
        records['test.1'] = SeqRecord(Seq('ATTTA', generic_dna), id ='test.1::Chr1:1-6', name = 'test.1::Chr1:1-6',
                           description = 'test.1::Chr1:1-6')
        records['test.2'] = SeqRecord(Seq('GC', generic_dna), id ='test.2::Chr1:6-12', name = 'test.2::Chr1:6-12',
                           description = 'test.2::Chr1:6-12'
                           )
        isoforms = [(1, 6, 'test.1'),(8, 14, 'test.2')]
        caseInput = ([records], isoforms, 14)
        caseOutput = 'ATTTA        '
        testCases.append((caseInput, caseOutput))
        
        records = collections.OrderedDict()
        records['test.1'] = SeqRecord(Seq('ATTTA', generic_dna), id ='test.1::Chr1:1-6', name = 'test.1::Chr1:1-6',
                           description = 'test.1::Chr1:1-6')
        records['test.2'] = SeqRecord(Seq('GCGCGC', generic_dna), id ='test.2::Chr1:6-12', name = 'test.2::Chr1:6-12',
                           description = 'test.2::Chr1:6-12'
                           )
        records['test.3'] = SeqRecord(Seq('TACTAC', generic_dna), id ='test.2::Chr1:6-12', name = 'test.2::Chr1:6-12',
                           description = 'test.2::Chr1:6-12'
                           )
        caseInput = ([records], [], 16)
        caseOutput = ''
        testCases.append((caseInput, caseOutput))
        # None type for sequence
        caseInput = (None, isoforms, 16)
        caseOutput = ''
        testCases.append((caseInput, caseOutput))
        # Negative XAxisMax
        records = collections.OrderedDict()
        records['test.1'] = SeqRecord(Seq('ATTTA', generic_dna), id ='test.1::Chr1:1-6', name = 'test.1::Chr1:1-6',
                           description = 'test.1::Chr1:1-6')
        records['test.2'] = SeqRecord(Seq('GCGCGC', generic_dna), id ='test.2::Chr1:6-12', name = 'test.2::Chr1:6-12',
                           description = 'test.2::Chr1:6-12'
                           )
        isoforms = [(1, 6, 'test.1'),(6, 12, 'test.2')]
        caseInput = ([records], isoforms, -12)
        caseOutput = ''
        testCases.append((caseInput, caseOutput))
        for i in testCases:
            self.assertEqual(iclip.generateMasterSequence(i[0][0], i[0][1], 0, i[0][2]), i[1])
            
    def testCreateDetailRow(self):
        testCases = []
        tableColors = ['rgb(0,0,0)', 'rgb(1,1,1)']
        testCases.append(('attribute', 'data', 1,  html.Tr(children=[html.Td(html.B('data'.replace('_', ' ').title())),
                                     html.Td(html.Table(children=
                                                        html.Tr(html.Td('attribute'.strip()))))],
                           style={'background-color': 'rgb(1,1,1)'}), None))
        testCases.append(('1;2;3;4;5;6', 'data', 2,  
                          html.Tr(children=[html.Td(html.B('data'.replace('_', ' ').title())),
                                    html.Td(html.Details(title = str(6) + ' values', 
                                        children = [
                                            html.Summary(str(6) + ' values'), 
                                            html.Table(children = 
                                                [
                                                    html.Tr(html.Td('1'.strip())),
                                                    html.Tr(html.Td('2'.strip())),
                                                    html.Tr(html.Td('3'.strip())),
                                                    html.Tr(html.Td('4'.strip())),
                                                    html.Tr(html.Td('5'.strip())),
                                                    html.Tr(html.Td('6'.strip()))
                                                ]
                                            )
                                        ]
                                    ))], style={'background-color': 'rgb(0,0,0)'}), 
                            None))
        testCases.append(('1;2;3;4;?5;6', 'data', 2,  
                          html.Tr(children=[html.Td(html.B('data'.replace('_', ' ').title())),
                                    html.Td(html.Details(title = str(6) + ' values', 
                                        children = [
                                            html.Summary(str(6) + ' values'), 
                                            html.Table(children = 
                                                [
                                                    html.Tr(html.Td('1'.strip())),
                                                    html.Tr(html.Td('2'.strip())),
                                                    html.Tr(html.Td('3'.strip())),
                                                    html.Tr(html.Td('4'.strip())),
                                                    html.Tr(html.Td(html.A('?5'[1:], href = '?5'[1:].strip(), target = '_blank'))),
                                                    html.Tr(html.Td('6'.strip()))
                                                ]
                                            )
                                        ]
                                    ))], style={'background-color': 'rgb(0,0,0)'}), 
                            None))
        falseCases = []
        falseCases.append(('1;2;3;4;?5;', 'data', 2,  
                          html.Tr(children=[html.Td(html.B('data'.replace('_', ' ').title())),
                                    html.Td(html.Details(title = str(6) + ' values', 
                                        children = [
                                            html.Summary(str(6) + ' values'), 
                                            html.Table(children = 
                                                [
                                                    html.Tr(html.Td('1'.strip())),
                                                    html.Tr(html.Td('2'.strip())),
                                                    html.Tr(html.Td('3'.strip())),
                                                    html.Tr(html.Td('4'.strip())),
                                                    html.Tr(html.Td(html.A('?5'[1:], href = '?5'[1:].strip(), target = '_blank'))),
                                                    html.Tr(html.Td('6'.strip()))
                                                ]
                                            )
                                        ]
                                    ))], style={'background-color': 'rgb(0,0,0)'}), 
                            None))                          
        for i in testCases:
            details.subTables = i[4]
            details.tableColors = tableColors
            self.assertEqual(details.createDetailRow(i[0], i[1], i[2]), i[3])
        for i in falseCases:
            details.subTables = i[4]
            details.tableColors = tableColors
            self.assertIsNot(details.createDetailRow(i[0], i[1], i[2]), i[3])

    def testCalculateEvents(self):
        testCases = []
        # Single event
        # chromStart, chromEnd, score, eventXValues, eventWidths, eventBases, eventScores, intervals, maxStack
        caseInput = ('test',0, 10, 0.5, {}, {}, {}, {}, [], 0)
        caseOutput = ({'test' : [4.5]}, {'test' : [10]}, {'test' : [0]}, {'test' : [0.5]}, [((0,10),0)], 1)
        testCases.append((caseInput, caseOutput))
        # Events overlapping
        # chromStart, chromEnd, score, eventXValues, eventWidths, eventBases, eventScores, intervals, maxStack
        caseInput = ('test', 0, 10, 0.5, {'test' : [4.5]}, {'test' : [10]}, {'test' : [0]}, {'test' : [0.5]}, [((0,10),0)], 1)
        caseOutput = ({'test' : [4.5,4.5]}, {'test' : [10,10]}, {'test' : [0,1.5]}, {'test' : [0.5,0.5]}, [((0,10),0),((0,10),1)], 2)
        testCases.append((caseInput, caseOutput))
        # Event bordering on right
        # chromStart, chromEnd, score, eventXValues, eventWidths, eventBases, eventScores, intervals, maxStack
        caseInput = ('test', 10, 15, 0.5, {'test' : [4.5]}, {'test' : [10]}, {'test' : [0]}, {'test' : [0.5]}, [((0,10),0)], 1)
        caseOutput = ({'test' : [4.5,12.0]}, {'test' : [10,5]}, {'test' : [0,0]}, {'test' : [0.5,0.5]}, [((0,10),0),((10,15),0)], 1)
        testCases.append((caseInput, caseOutput))
        # Event bordering on left
        # chromStart, chromEnd, score, eventXValues, eventWidths, eventBases, eventScores, intervals, maxStack
        caseInput = ('test', 0, 5, 0.5, {'test' : [7]}, {'test' : [5]}, {'test' : [0]}, {'test' : [0.5]}, [((5,10),0)], 1)
        caseOutput = ({'test' : [7,2]}, {'test' : [5,5]}, {'test' : [0,0]}, {'test' : [0.5,0.5]}, [((5,10),0),((0,5),0)], 1)
        testCases.append((caseInput, caseOutput))
        for i in testCases:
            xVals = i[0][4]
            widths = i[0][5]
            bases = i[0][6]
            scores = i[0][7]
            intervals = i[0][8]
            overlap = rna.calculateEvents(i[0][0], i[0][1], i[0][2],
                            i[0][3], i[0][4], i[0][5], i[0][6], i[0][7], i[0][8], i[0][9])
            self.assertEqual((xVals, widths, bases, scores, intervals, overlap), i[1])

    def testCalculateEventsScoreColored(self):
        testCases = []
        # Single event
        # chromStart, chromEnd, score, eventXValues, eventWidths, eventBases, eventScores, intervals, maxStack
        caseInput = (0, 10, 0.5, [], [], [], [], [], 0)
        caseOutput = ([4.5], [10], [0], [0.5], [((0,10),0)], 1)
        testCases.append((caseInput, caseOutput))
        # Events overlapping
        # chromStart, chromEnd, score, eventXValues, eventWidths, eventBases, eventScores, intervals, maxStack
        caseInput = (0, 10, 0.5, [4.5], [10], [0], [0.5], [((0,10),0)], 1)
        caseOutput = ([4.5,4.5], [10,10], [0,1.5], [0.5,0.5], [((0,10),0),((0,10),1)], 2)
        testCases.append((caseInput, caseOutput))
        # Event bordering on right
        # chromStart, chromEnd, score, eventXValues, eventWidths, eventBases, eventScores, intervals, maxStack
        caseInput = (10, 15, 0.5, [4.5], [10], [0], [0.5], [((0,10),0)], 1)
        caseOutput = ([4.5,12.0], [10,5], [0,0], [0.5,0.5], [((0,10),0),((10,15),0)], 1)
        testCases.append((caseInput, caseOutput))
        # Event bordering on left
        # chromStart, chromEnd, score, eventXValues, eventWidths, eventBases, eventScores, intervals, maxStack
        caseInput = (0, 5, 0.5, [7], [5], [0], [0.5], [((5,10),0)], 1)
        caseOutput = ([7,2], [5,5], [0,0], [0.5,0.5], [((5,10),0),((0,5),0)], 1)
        testCases.append((caseInput, caseOutput))
        for i in testCases:
            xVals = i[0][3]
            widths = i[0][4]
            bases = i[0][5]
            scores = i[0][6]
            intervals = i[0][7]
            overlap = rna.calculateEventsScoreColored(i[0][0], i[0][1],
                            i[0][2], i[0][3], i[0][4], i[0][5], i[0][6], i[0][7], i[0][8])
            self.assertEqual((xVals, widths, bases, scores, intervals, overlap), i[1])
    
    def testSetUpBlockConstraints(self):
        testCases = []
        # Test overlap on right side of region and with chromStart/end equal xAxisMin/Max
        # chromStart, chromEnd, blockStarts, blockSizes, xAxisMin, xAxisMax
        caseInput = (0, 300, "200,280,340","40,40,40", 0, 300)
        # blockStartsF, blockSizesF
        caseOutput = ([200,280,-1], [40,20,40], 'cont')
        testCases.append((caseInput, caseOutput))
        
        # Test overlap on right side of region
        # chromStart, chromEnd, blockStarts, blockSizes, xAxisMin, xAxisMax
        caseInput = (100, 400, "100,180,240","40,40,40", 100, 300)
        # blockStartsF, blockSizesF
        caseOutput = ([200,280,-1], [40,20,40], 'left')
        testCases.append((caseInput, caseOutput))
        
        # Test overlap on left side of region
        # chromStart, chromEnd, blockStarts, blockSizes, xAxisMin, xAxisMax
        caseInput = (0, 200, "40,80,140","40,40,40", 100, 300)
        # blockStartsF, blockSizesF
        caseOutput = ([-1,100,140], [40,20,40], 'right')
        testCases.append((caseInput, caseOutput))
        
        # Test extension over both sides of the region
        # chromStart, chromEnd, blockStarts, blockSizes, xAxisMin, xAxisMax
        caseInput = (0, 500, "40,80,240,400","40,40,70,40", 100, 300)
        # blockStartsF, blockSizesF
        caseOutput = ([-1,100,240,-1], [40,20,60,40], 'both')
        testCases.append((caseInput, caseOutput))
        
        # Test inclusion in region
        # chromStart, chromEnd, blockStarts, blockSizes, xAxisMin, xAxisMax
        caseInput = (120, 200, "40,80","40,40", 100, 300)
        # blockStartsF, blockSizesF
        caseOutput = ([160, 200], [40, 40], 'cont')
        testCases.append((caseInput, caseOutput))       
        for i in testCases:
            self.assertEqual(iclip.setUpBlockConstraints(i[0][0],i[0][1],i[0][2],i[0][3],i[0][4],i[0][5]), i[1])
    
    def testCalculateBlocks(self):
        testCases = []
        
        # thickStart, thickEnd, blockStart, blockEnd, blockHeight
        caseInput = (10,50, 20, 40, 0.4)
        # blockVals, blockYs, blockWidths
        caseOutput = ([29.5],[0.4],[20])
        testCases.append((caseInput,caseOutput))
        
        # BLock lies completely in coding region
        # thickStart, thickEnd, blockStart, blockEnd, blockHeight
        caseInput = (0,100, 20, 40, 0.4)
        # blockVals, blockYs, blockWidths
        caseOutput = ([29.5],[0.4],[20])
        testCases.append((caseInput,caseOutput))
        # thickStart, thickEnd, blockStart, blockEnd, blockHeight
        caseInput = (0,100, 21, 40, 0.4)
        # blockVals, blockYs, blockWidths
        caseOutput = ([30],[0.4],[19])
        testCases.append((caseInput,caseOutput))
        
        # Block lies right of coding region
        # thickStart, thickEnd, blockStart, blockEnd, blockHeight
        caseInput = (0,10, 20, 40, 0.4)
        # blockVals, blockYs, blockWidths
        caseOutput = ([30],[0.2],[21])
        testCases.append((caseInput,caseOutput))
        # thickStart, thickEnd, blockStart, blockEnd, blockHeight
        caseInput = (0,10, 21, 41, 0.4)
        # blockVals, blockYs, blockWidths
        caseOutput = ([31],[0.2],[21])
        testCases.append((caseInput,caseOutput))
        # thickStart, thickEnd, blockStart, blockEnd, blockHeight
        caseInput = (0,10, 21, 40, 0.4)
        # blockVals, blockYs, blockWidths
        caseOutput = ([30.5],[0.2],[20])
        testCases.append((caseInput,caseOutput))
        
        # Block lies left of coding region
        # thickStart, thickEnd, blockStart, blockEnd, blockHeight
        caseInput = (20,30, 0, 10, 0.4)
        # blockVals, blockYs, blockWidths
        caseOutput = ([4.5],[0.2],[10])
        testCases.append((caseInput,caseOutput))
        # thickStart, thickEnd, blockStart, blockEnd, blockHeight
        caseInput = (20,30, 0, 11, 0.4)
        # blockVals, blockYs, blockWidths
        caseOutput = ([5],[0.2],[11])
        testCases.append((caseInput,caseOutput))
        # thickStart, thickEnd, blockStart, blockEnd, blockHeight
        caseInput = (20,30, 1, 11, 0.4)
        # blockVals, blockYs, blockWidths
        caseOutput = ([5.5],[0.2],[10])
        testCases.append((caseInput,caseOutput))
        
        # Block overlaps coding region on the left
        # thickStart, thickEnd, blockStart, blockEnd, blockHeight
        caseInput = (0,10, 5, 15, 0.4)
        # blockVals, blockYs, blockWidths
        caseOutput = ([7,12],[0.4,0.2],[5,5])
        testCases.append((caseInput,caseOutput))
        # thickStart, thickEnd, blockStart, blockEnd, blockHeight
        caseInput = (0,10, 6, 15, 0.4)
        # blockVals, blockYs, blockWidths
        caseOutput = ([7.5,12],[0.4,0.2],[4,5])
        testCases.append((caseInput,caseOutput))
         # thickStart, thickEnd, blockStart, blockEnd, blockHeight
        caseInput = (0,10, 6, 16, 0.4)
        # blockVals, blockYs, blockWidths
        caseOutput = ([7.5,12.5],[0.4,0.2],[4,6])
        testCases.append((caseInput,caseOutput))
        
        # Block overlaps coding region on the right
        # thickStart, thickEnd, blockStart, blockEnd, blockHeight
        caseInput = (5,15, 0, 10, 0.4)
        # blockVals, blockYs, blockWidths
        caseOutput = ([2,7],[0.2,0.4],[5,5])
        testCases.append((caseInput,caseOutput))
        # thickStart, thickEnd, blockStart, blockEnd, blockHeight
        caseInput = (5,15, 1, 10, 0.4)
        # blockVals, blockYs, blockWidths
        caseOutput = ([2.5,7],[0.2,0.4],[4,5])
        testCases.append((caseInput,caseOutput))
        # thickStart, thickEnd, blockStart, blockEnd, blockHeight
        caseInput = (5,15, 1, 9, 0.4)
        # blockVals, blockYs, blockWidths
        caseOutput = ([2.5,6.5],[0.2,0.4],[4,4])
        testCases.append((caseInput,caseOutput))
        
        # Block completely contains coding region
        # thickStart, thickEnd, blockStart, blockEnd, blockHeight
        caseInput = (5,15, 1, 11, 0.4)
        # blockVals, blockYs, blockWidths
        caseOutput = ([2.5,7.5],[0.2,0.4],[4,6])
        testCases.append((caseInput,caseOutput))
        # thickStart, thickEnd, blockStart, blockEnd, blockHeight
        caseInput = (5,15, 5, 15, 0.4)
        # blockVals, blockYs, blockWidths
        caseOutput = ([9.5],[0.4],[10])
        testCases.append((caseInput,caseOutput))
        # thickStart, thickEnd, blockStart, blockEnd, blockHeight
        caseInput = (5,16, 5, 16, 0.4)
        # blockVals, blockYs, blockWidths
        caseOutput = ([10],[0.4],[11])
        testCases.append((caseInput,caseOutput))
        
        # Block has 0 size
        # thickStart, thickEnd, blockStart, blockEnd, blockHeight
        caseInput = (5,15, 1, 1, 0.4)
        # blockVals, blockYs, blockWidths
        caseOutput = ([],[],[])
        testCases.append((caseInput,caseOutput))
        for i in testCases:
            inputBlockVals = []
            inputBlockYs = []
            inputBlockWidths = []
            iclip.calculateBlocks(i[0][0],i[0][1],i[0][2],i[0][3], inputBlockVals, inputBlockWidths, inputBlockYs, i[0][4])
            output = (inputBlockVals, inputBlockYs, inputBlockWidths)
            self.assertEqual(output, i[1])
            
if __name__ == '__main__':
    unittest.main()
```
