## Additional File 1 for "SEQing: web-based visualization of iCLIP and RNA-seq data in an interactive python framework": validator.py

```
#!/usr/bin/env python
# -*- coding: utf-8 -*-

""" Interactive visualizaton for iCLIP-seq and RNA-seq data"""
import runpy
from argparse import ArgumentParser
from pathlib import Path
from xml.dom import minidom
import pickle
import os
import hashlib
import itertools
import pandas
from Bio import SeqIO
from Bio.Alphabet import generic_dna
import converter
import time
import gzip
import bz2
import zipfile

__author__ = "Yannik Bramkamp"


start = time.time()
geneIndex = pandas.DataFrame()
plotColors = []
geneAnnotations = []
sequences = []
ensembl = False
geneDescriptions = None
descAvail = True
dropList = []
advancedDescriptions = None
subTables = None
dsElements = 0 # number of traces per dataset, i.e Rawdata+ bindingsites = 2
bsRawDFs = {}
rawAvail = False # Raw data available
bsProcDFs = {}
procAvail = False # proc data available
spliceSetNames = [[],[]]
spliceElements = 0
fileDict = {} # This dictionary will holde the file indexes for each dataset
spliceAvail = False # splice data available
spliceEventsAvail = False  # splice events available
spliceEventsDFs = {}
spliceEventsElements = 0
spliceEventNames = [[],[]]
spliceEventTypes = []
dataSetNames = []
# Colors for dna sequence display
colorA = 'rgb(0, 150, 0)'
colorC = 'rgb(15,15,255)'
colorG = 'rgb(209, 113, 5)'
colorT = 'rgb(255, 9, 9)'
# Map for data track colors
colorMap = {}
# Create dictionary for coverage track colors
coverageColors = ['rgb(255,0,0)', 'rgb(255,165,0)','rgb(255,255,0)','rgb(0,0,255)', 'rgb(128,0,128)']
coverageColorDict = {}
eventColors = ['rgb(0,0,255)', 'rgb(255,0,0)', 'rgb(0,255,0)', 'rgb(128,0,128)', 'rgb(255,165,0)']
chunkSize = 10000
spliceEventColors = {} # dictionary for slice event colors
# Headers for the data files, files obviously need to conform to these headers for the visualization to work
bedHeader = ['chrom','chromStart','chromEnd','transID','score','strand','thickStart',
             'thickEnd','itemRGB','blockCount','blockSizes','blockStarts']
bsHeader = ['chrom', 'chromStart','chromEnd','type', 'score', 'strand']
rawHeader = ['chrom','chromStart','chromEnd','count']
gtfheader = ['seqname', 'source', 'feature', 'start', 'end', 'score',
               'strand', 'frame', 'attribute']


print('Loading gene annotation files.')


def validateGTF(df):
    """Validates gtf files. Returns True and an empty String if dataframe is valid,
    else returns false and an error message.
    
    Positional arguments:
    df -- Dataframe to be validated
    """
    try:
        msg = ''
        if df.isnull().values.any() == True:
            msg = 'Missing values' + '\n' + str(df.isnull().sum())
            return [False, msg]
        if (all(x in ['+', '-'] for x in df['strand'].cat.categories.tolist())) != True:
            msg = 'Bad strand symbol(has to be + or -'
            return [False, msg]
        return [True, msg] 
    except (TypeError, AttributeError, KeyError):
        return [False, 'Not a valid dataframe']   

def validateBed12(df):
    """Validates 12 column bed files. Returns True and an empty String if dataframe is valid,
    else returns false and an error message.
    
    Positional arguments:
    df -- Dataframe to be validated
    """
    try:
        msg = ''
        if df.isnull().values.any() == True:        
            msg = 'Missing values' + '\n' + str(df.isnull().sum())
            return [False, msg]
        if (all(x in ['+', '-'] for x in df['strand'].cat.categories.tolist())) != True:
            msg = 'Bad strand symbol(has to be + or -'
            return [False, msg]
        if all(y.isdigit() for z in df['blockSizes'].map(lambda x: x.split(',')[:-1]).tolist()[0] for y in z ) == False:
            msg = 'Column blockSizes contains non int values'
            return [False, msg]    
        if all(y.isdigit() for z in df['blockStarts'].map(lambda x: x.split(',')[:-1]).tolist()[0] for y in z ) == False:
            msg = 'Column blockStarts contains non int values'
            return [False, msg]
        return [True, msg]
    except (TypeError, AttributeError, KeyError):
        return [False, 'Not a valid dataframe']   

def validateBedGraph(df):
    """Validates 4 column bedgraph files. Returns True and an empty String if dataframe is valid,
    else returns false and an error message.
    
    Positional arguments:
    df -- Dataframe to be validated
    """
    try:
        msg = ''
        if df.empty:
            return [False, 'Not a valid dataframe']           
        if df.isnull().values.any() == True:        
            msg = 'Missing values' + '\n' + str(df.isnull().sum())
            return [False, msg]
        return [True, msg]
    except (TypeError, AttributeError, KeyError):
        return [False, 'Not a valid dataframe']   

def validateBed(df):
    """Validates 6 column bed files. Returns True and an empty String if dataframe is valid,
    else returns false and an error message.
    
    Positional arguments:
    df -- Dataframe to be validated
    """
    try:
        msg = ''
        if df.isnull().values.any() == True:        
            msg = 'Missing values' + '\n' + str(df.isnull().sum())
            return [False, msg]
        if (all(x in ['+', '-'] for x in df['strand'].cat.categories.tolist())) != True:
            msg = 'Bad strand symbol(has to be + or -)'
            return [False, msg]
        return [True, msg]
    except (TypeError, AttributeError, KeyError):
        return [False, 'Not a valid dataframe']   
    
def isRGB(color):
    """ Check if the provided strings match the required rgb format

    Positional arguments:
    color -- a single color string
    """
    try:
        if color[0:4] != 'rgb(':
            return False
        if color[-1:] != ')':
            return False
        if len(color[4:-1].split(',')) != 3:
            return False
        for i in color[4:-1].split(','):
            if i.replace(' ', '').isdigit() == False:
                return False
            if int(i.replace(' ', '')) < 0 or int(i.replace(' ', '')) > 255:
                return False
        return True
    except TypeError:
        return False

def md5Gzip(fname):
    hash_md5 = hashlib.md5()

    with gzip.open(fname, 'rb') as f:
        for chunk in iter(lambda: f.read(4096), b""):
            hash_md5.update(chunk)
    return hash_md5

def md5Bz2(fname):
    hash_md5 = hashlib.md5()

    with bz2.open(fname, 'rb') as f:
        for chunk in iter(lambda: f.read(4096), b""):
            hash_md5.update(chunk)
    return hash_md5

def md5Zip(fname):
    hash_md5 = hashlib.md5()
    with zipfile.open(fname, 'rb') as f:
        for chunk in iter(lambda: f.read(4096), b""):
            hash_md5.update(chunk)
    return hash_md5

def loadAnnotations():
    for idx, i in enumerate(geneAnnotationPaths):
        try:
            typeGuess = converter.check_input_file(str(i))
            if typeGuess.header_present == True:
                header = 1  
            else:
                header = None
            print('Loading file ' + str(idx+1) )
            if typeGuess.file_type == 'BED12':           
    #        try:
    #            if i.suffix.lower() =='.bed':
                if typeGuess.zipped == True:
                    if typeGuess.zip_type == 'gzip':
                        checksum = md5Gzip(str(i))
                    elif typeGuess.zio_type == 'bzip2':
                        checksum = md5Bz2(str(i))
                    elif typeGuess.zio_type == 'zip':
                        checksum = md5Zip(str(i))       
                else:
                    checksum = hashlib.md5(open(str(i)).read().encode('utf-8'))
                if checksums.get(str(i.stem), None) != checksum.hexdigest():
                    dtypes = {'chrom' : 'category', 'chromStart' : 'uint32','chromEnd': 'uint32','transID' : 'object','score' : 'int16','strand' : 'category','thickStart' : 'uint64',
                 'thickEnd' : 'uint64', 'blockCount' : 'uint32','blockSizes' : 'object','blockStarts' : 'object'}
                    df = pandas.read_csv(i, sep = '\t', compression='infer', comment = '#', names = bedHeader, dtype = dtypes, header = header)
                    df = df.join(geneIndex.set_index('transID'), on='transID') 
                    validation = validateBed12(df)
                    if validation[0] == True:
                        geneAnnotations.append(df)
                        out = open(binFilePath + str(i.stem)+'.bin', 'wb')
                        pickle.dump(df, out)
                        out.close()
                    else:
                        print('Error in file ' + str(i) + ':')
                        print(validation[1])
                    checksums[str(i.stem)] = checksum.hexdigest()
                else:
                    try:
                        df = pickle.load(open(binFilePath + str(i.stem)+'.bin', 'rb'))
                        geneAnnotations.append(df)
                        print('Loaded from pickle')
                    except IOError:
                        print('pickle not  found, loading from raw file')
                        dtypes = {'chrom' : 'category', 'chromStart' : 'uint32','chromEnd': 'uint32','transID' : 'object','score' : 'int16','strand' : 'category','thickStart' : 'uint64',
                                  'thickEnd' : 'uint64', 'blockCount' : 'uint32','blockSizes' : 'object','blockStarts' : 'object'}
                        df = pandas.read_csv(i, sep = '\t', compression='infer', comment = '#', names = bedHeader, dtype = dtypes, header = header)
                        df = df.join(geneIndex.set_index('transID'), on='transID') 
                        validation = validateBed12(df)
                        if validation[0] == True:
                            geneAnnotations.append(df)
                            out = open(binFilePath + str(i.stem)+'.bin', 'wb')
                            pickle.dump(df, out)
                            out.close()
                        else:
                            print('Error in file ' + str(i) + ':')
                            print(validation[1])
                    except UnicodeDecodeError:
                        print('Error decoding pickle binary file, will load from raw file instead')
                        dtypes = {'chrom' : 'category', 'chromStart' : 'uint32','chromEnd': 'uint32','transID' : 'object','score' : 'int16','strand' : 'category','thickStart' : 'uint64',
                                  'thickEnd' : 'uint64', 'blockCount' : 'uint32','blockSizes' : 'object','blockStarts' : 'object'}
                        df = pandas.read_csv(i, sep = '\t', compression='infer', comment = '#', names = bedHeader, dtype = dtypes, header = header)
                        df = df.join(geneIndex.set_index('transID'), on='transID')                     
                        validation = validateBed12(df)
                        if validation[0] == True:
                            geneAnnotations.append(df)
                            out = open(binFilePath + str(i.stem)+'.bin', 'wb')
                            pickle.dump(df, out)
                            out.close()
                        else:
                            print('Error in file ' + str(i) + ':')
                            print(validation[1])
                    except ModuleNotFoundError:
                        print('Pickle was created using different package versions, will load from raw file instead')
                        dtypes = {'chrom' : 'category', 'chromStart' : 'uint32','chromEnd': 'uint32','transID' : 'object','score' : 'int16','strand' : 'category','thickStart' : 'uint64',
                                  'thickEnd' : 'uint64', 'blockCount' : 'uint32','blockSizes' : 'object','blockStarts' : 'object'}
                        df = pandas.read_csv(i, sep = '\t', compression='infer', comment = '#', names = bedHeader, dtype = dtypes, header = header)
                        df = df.join(geneIndex.set_index('transID'), on='transID')                          
                        validation = validateBed12(df)
                        if validation[0] == True:
                            geneAnnotations.append(df)
                            out = open(binFilePath + str(i.stem)+'.bin', 'wb')
                            pickle.dump(df, out)
                            out.close()
                        else:
                            print('Error in file ' + str(i) + ':')
                            print(validation[1])
            elif typeGuess.file_type == 'GTF':
    #            if i.suffix.lower() == '.gtf':
                if typeGuess.zipped == True:
                    if typeGuess.zip_type == 'gzip':
                        checksum = md5Gzip(str(i))
                    elif typeGuess.zio_type == 'bzip2':
                        checksum = md5Bz2(str(i))
                    elif typeGuess.zio_type == 'zip':
                        checksum = md5Zip(str(i))       
                else:
                    checksum = hashlib.md5(open(str(i)).read().encode('utf-8'))
                if checksums.get(str(i.stem), None) != checksum.hexdigest():
                    dtypes = {'seqname' : 'object', 'source' : 'object', 'feature' : 'object', 'start' : 'uint32', 'end': 'uint32', 'score' : 'object',
                              'strand' : 'category', 'frame' : 'object', 'attribute' : 'object'}
                    df = pandas.read_csv(i, sep = '\t', compression='infer', comment = '#', names = gtfheader, dtype = dtypes, header = header)
                    validation = validateGTF(df)
                    if validation[0] == True:
                        df = converter.convertGTFToBed(df)
                        out = open(binFilePath + str(i.stem)+'.bin', 'wb')
                        pickle.dump(df, out)
                        out.close()
                        geneAnnotations.append(df)
                    else:
                        print('Error in file ' + str(i) + ':')
                        print(validation[1])
                    checksums[str(i.stem)] = checksum.hexdigest()                        
                else:
                    try:
                        df = pickle.load(open(binFilePath + str(i.stem)+'.bin', 'rb'))
                        geneAnnotations.append(df)
                        print('Loaded from pickle')
                    except IOError:
                        print('pickle not  found, loading from raw file')
                        dtypes = {'seqname' : 'object', 'source' : 'object', 'feature' : 'object', 'start' : 'uint32', 'end': 'uint32', 'score' : 'category',
                              'strand' : 'category', 'frame' : 'object', 'attribute' : 'object'}
                        df = pandas.read_csv(i, sep = '\t', compression='infer', comment = '#', names = gtfheader, dtype = dtypes, header = header)
                        validation = validateGTF(df)
                        if validation[0] == True:
                            df = converter.convertGTFToBed(df)
                            out = open(binFilePath + str(i.stem)+'.bin', 'wb')
                            pickle.dump(df, out)
                            out.close()
                            geneAnnotations.append(df)
                        else:
                            print('Error in file ' + str(i) + ':')
                            print(validation[1])
                    except UnicodeDecodeError:
                        print('Error decoding pickle binary file, will load from raw file instead')
                        dtypes = {'seqname' : 'object', 'source' : 'object', 'feature' : 'object', 'start' : 'uint32', 'end': 'uint32', 'score' : 'category',
                              'strand' : 'category', 'frame' : 'object', 'attribute' : 'object'}
                        df = pandas.read_csv(i, sep = '\t', comment = '#', compression='infer', names = gtfheader, dtype = dtypes, header = header)
                        validation = validateGTF(df)
                        if validation[0] == True:
                            df = converter.convertGTFToBed(df)
                            out = open(binFilePath + str(i.stem)+'.bin', 'wb')
                            pickle.dump(df, out)
                            out.close()
                            geneAnnotations.append(df)
                        else:
                            print('Error in file ' + str(i) + ':')
                            print(validation[1])
                    except ModuleNotFoundError:
                        print('Pickle was created using different package versions, will load from raw file instead')
                        dtypes = {'seqname' : 'object', 'source' : 'object', 'feature' : 'object', 'start' : 'uint32', 'end': 'uint32', 'score' : 'category',
                              'strand' : 'category', 'frame' : 'object', 'attribute' : 'object'}
                        df = pandas.read_csv(i, sep = '\t', comment = '#', compression='infer', names = gtfheader, dtype = dtypes, header = header)
                        validation = validateGTF(df)
                        if validation[0] == True:
                            df = converter.convertGTFToBed(df)
                            out = open(binFilePath + str(i.stem)+'.bin', 'wb')
                            pickle.dump(df, out)
                            out.close()
                            geneAnnotations.append(df)
                        else:
                            print('Error in file ' + str(i) + ':')
                            print(validation[1])
            elif typeGuess.file_type == 'unsupported':
                print('Invalid file format, please use only .bed or .gtf files')              
        except FileNotFoundError:
            print('File ' + str(i) + ' not found, skipping')
        except ValueError as e:
            print('File ' + str(i) + ' had errornous datatypes or missing values, skipping: ' + str(e))                
    if len(geneAnnotations) == 0:
        print('No valid gene annotation files found, terminating.')
        exit()
    
    # Write new checksums file
    try:
        out = open(binFilePath + 'checksums', 'wb')
        pickle.dump(checksums, out)
        out.close()
    except FileNotFoundError:
        pass

def loadSequences():
    try:
        for i in fastaPaths:
            try:
                seq = SeqIO.parse(str(i), 'fasta', alphabet = generic_dna)
                for record in seq:
                    if record.description == record.name:
                        ensembl = False
                    else:
                        ensembl = True
                    seq = itertools.chain([record], seq)
                    break
                if ensembl != True:
                    sequences.append(SeqIO.to_dict(seq, 
                        key_function = lambda rec : rec.description.split(':')[0]
                        )
                    )
                else:
                    sequences.append(SeqIO.to_dict(seq, 
                        key_function = lambda rec : rec.name
                        )
                    )
            except FileNotFoundError:
                print('Sequence annotations for coding genes not found, proceeding without')
    except TypeError:
        pass
    
def loadBasicDescriptions():
    global descAvail, geneDescriptions
    try:
        geneDescriptions = pandas.read_csv(descriptionPath, compression='infer',
                            names = ['ensembl_gene_id', 'description', 'external_gene_name'], sep = '\t', usecols=[0,1,2])        
        # Filter for genes that are actually in the dataset
        geneDescriptions = geneDescriptions[geneDescriptions['ensembl_gene_id'].isin(geneNames)]
        geneDescriptions.fillna(':',inplace = True)
    except FileNotFoundError:
        print('Description file not found, proceeding without')
        descAvail = False
    except ValueError:
        descAvail = False

def loadAdvancedDescriptions():
    global advancedDescriptions
    try:
        advancedDescriptions = pandas.read_csv(advancedDescPath, compression='infer', sep = '\t')
        if len(advancedDescriptions.index.values) > 0:
            if 'gene_ids' not in list(advancedDescriptions.columns.values):
                print('Advanced description file does not contain "gene_ids" column, ignoring file.')
                advancedDescriptions = None
            elif list(advancedDescriptions.index.values) != list(range(0, len(advancedDescriptions.index.values))):
                print('You might be missing a column name in your advanced descriptions. Ignoring file')
                advancedDescriptions = None
    except FileNotFoundError:
        print('Adanced description file could not be found, ignoring.')
        advancedDescriptions = None
    except ValueError:
        advancedDescriptions = None

def loadSubTables():
    global subTables
    try:
        subTables = pandas.read_csv(subTablePath, compression='infer', sep = '\t', names = ['column_id', 'columns'])
    except FileNotFoundError:
        print('Sub table file could not be found, ignoring.')
        subTables = None
    except ValueError:
        subTables = None
        
def loadICLIPData():
    global rawAvail, dsElements
    if len(bindingSiteRawPaths) > 0:
        print('Loading iCLIP data.')
        for i in bindingSiteRawPaths:
            try:
                dtypes = {'chrom' : 'category' ,'chromStart' : 'uint64','chromEnd' : 'uint64', 'count' : 'uint32'}
                df = pandas.read_csv(i, compression='infer', sep = '\t', names = rawHeader, dtype = dtypes)
                validation = validateBedGraph(df)
                if validation[0] == True:
                    if i.stem.split('_')[0] not in dataSetNames:
                        dataSetNames.append(i.stem.split('_')[0])
                        bsRawDFs.update({str(dataSetNames[-1]) : df})
                    else:
                        print('Warning, you are using the same prefix for multiple iCLIP files, file ' + str(i) + ' will be ignored')
                else:
                    print('Error in file ' + str(i) + ':')
                    print(validation[1])
            except FileNotFoundError:
                print('File '+str(i) + ' was not found')
            except ValueError as e:
                print('File ' + str(i.stem) + ' had errornous data types or missing values: ' + str(e))        
        print('Done.')
    if len(bsRawDFs) > 0:
        rawAvail = True
        dsElements += 1
        
def loadBSData():
    global dsElements, procAvail
    if len(bindingSitePaths) > 0:
        print('Loading bindings site data.')
    for i in bindingSitePaths:
        if i.stem.split('_')[0] in dataSetNames:
            try:
                dtypes = {'chrom' : 'category', 'chromStart' : 'uint64','chromEnd' : 'uint64','type' : 'category', 'score' : 'float32', 'strand' : 'category'}
                df = pandas.read_csv(i, compression='infer', sep = '\t', names = bsHeader, dtype = dtypes)
                validation = validateBed(df)
                if validation[0] == True:
                    if i.stem.split('_')[0] in bsProcDFs:
                        print('Warning, you are using the same prefix for multiple binding site files, file ' + str(i) + ' will be ignored')
                    else:
                        bsProcDFs.update({i.stem.split('_')[0] : df})
                else:
                    print('Error in file ' + str(i) + ':')
                    print(validation[1])                   
            except FileNotFoundError:
                print('File '+str(i) + ' was not found')  
            except ValueError as e:
                print('File ' + str(i.stem) + ' had errornous datatypes or missing values, skipping: ' + str(e))    
        else:
            print('No corresponding raw data found for data set ' + i.stem.split('_')[0])
    if len(bindingSitePaths) > 0:
        print('Done.')
    if len(bsProcDFs) > 0:
        dsElements +=1
        procAvail = True

def loadCoverageData():
    global spliceElements, spliceAvail
    if len(spliceSitePaths) > 0:
        print('Loading RNA-seq data')
    for path in spliceSitePaths:
        try:
            checksum = hashlib.md5(open(str(path)).read().encode('utf-8'))
        except FileNotFoundError:
            print('Error loading file ' + str(path) + ', skipping.')
            continue
        try:
            file_name = path.stem.split('_')[0]+'_'+path.stem.split('_')[1]
        except IndexError:
            file_name = path.stem.split('_')[0] 
        print(file_name)
        if coverageChecksums.get(str(path.stem), None) != checksum.hexdigest():
            try: 
                dtypes = {'chrom' : 'category', 'chromStart' : 'uint64','chromEnd' : 'uint64','type' : 'category', 'score' : 'float32', 'strand' : 'category'}
                df = pandas.read_csv(path, compression='infer', sep= '\t', names= rawHeader, dtype = dtypes)
                validation = validateBedGraph(df)
                coverageChecksums[str(path.stem)] = checksum.hexdigest()
            except FileNotFoundError:
                validation = [False]
            except ValueError as e:
                print('File ' + str(path.stem) + ' had errornous datatypes or missing values, skipping: ' + str(e))                       
            if validation[0]:
                df.sort_values(by=['chromStart'])
                # Split dataframe into small parts, these will be pickled and loaded on demand.
                # Store covered region as minimum starting point and maximum ending point in the file name.
                dfList = [df.iloc[i:i+chunkSize,] for i in range(0, len(df),chunkSize)]
                # This index will be used to filter out relevant files during runtime
                fileIndex = pandas.DataFrame(columns = ['start', 'end', 'fileName'])
                for i in dfList: 
                    end = i['chromEnd'].max()
                    start = i['chromStart'].min()
                    fileName = binFilePath + 'coverage/' + str(file_name) + "_" + str(start) + "_" + str(end) + '.bin'
                    fileIndex.loc[len(fileIndex)] = [start, end, fileName]
                    out = open(fileName, 'wb')
                    pickle.dump(i, out)
                    out.close()
                indexOut = open(binFilePath + 'coverage/' + str(file_name) + '_' + 'index.bin', 'wb')
                pickle.dump(fileIndex, indexOut)
                indexOut.close()
                fileDict.update({file_name : fileIndex})
                dfList = []
                # Add the dataset to the list of datasets, check  for number of underscores
                if path.stem.split('_')[0] not in spliceSetNames[1]:
                    try:
                        spliceSetNames[0].append(path.stem.split('_')[1])
                        spliceSetNames[1].append(path.stem.split('_')[0])
                    except IndexError:
                        spliceSetNames[1].append(path.stem.split('_')[0])
                        spliceSetNames[0].append(path.stem.split('_')[0])
                out = open(binFilePath + str(path.stem)+'.bin', 'wb')
                pickle.dump(df, out)
                out.close()
        else: # Checksum matches, try to load old index from pickle
            try:
                fileIndex = pickle.load(open(binFilePath + 'coverage/' + str(file_name) + '_' + 'index.bin', 'rb'))
                fileDict.update({file_name : fileIndex})
                if path.stem.split('_')[0] not in spliceSetNames[1]:
                    try:
                        spliceSetNames[0].append(path.stem.split('_')[1])
                        spliceSetNames[1].append(path.stem.split('_')[0])
                    except IndexError:
                        spliceSetNames[1].append(path.stem.split('_')[0])
                        spliceSetNames[0].append(path.stem.split('_')[0])
            except (FileNotFoundError, UnicodeDecodeError, IOError, ImportError):
                try:
                    df = pandas.read_csv(path, compression='infer', sep= '\t', names= rawHeader, dtype = dtypes)
                    validation = validateBedGraph(df)
                except FileNotFoundError:
                    validation = [False]
                except ValueError as e:
                    print('File ' + str(path.stem) + ' had errornous datatypes or missing values, skipping: ' + str(e))    
                if validation[0]:
                    df.sort_values(by=['chromStart'])
                    dfList = [df.iloc[i:i+chunkSize,] for i in range(0, len(df),chunkSize)]
                    fileIndex = pandas.DataFrame(columns = ['start', 'end', 'fileName'])
                    for i in dfList:
                        end = i['chromEnd'].max()    
                        start = i['chromStart'].min()
                        fileName = binFilePath + 'coverage/' + str(file_name) + "_" + str(start) + "_" + str(end) + '.bin'
                        fileIndex.loc[len(fileIndex)] = [start, end, fileName]
                        out = open(fileName, 'wb')
                        pickle.dump(i, out)
                        out.close()
                    indexOut = open(binFilePath + 'coverage/' + str(file_name) + '_' + 'index.bin', 'wb')
                    print(binFilePath + 'coverage/' + str(file_name) + '_' + 'index.bin')
                    pickle.dump(fileIndex, indexOut)
                    indexOut.close()
                    fileDict.update({file_name : fileIndex})
                    dfList = []
                    if path.stem.split('_')[0] not in spliceSetNames[1]:
                        try:
                            spliceSetNames[0].append(path.stem.split('_')[1])
                            spliceSetNames[1].append(path.stem.split('_')[0])
                        except IndexError:
                            spliceSetNames[1].append(path.stem.split('_')[0])
                            spliceSetNames[0].append(path.stem.split('_')[0])
                else:
                    print('Error loading file ' + str(path))
                        
    if len(fileDict.keys()) > 0:
        spliceElements += 1
        spliceAvail = True
    print('Done.')   

def loadSpliceEvents():
    global spliceEventsAvail, spliceEventsElements
    if len(spliceEventsPaths) > 0:
        print('Loading splice event data')
    for i in spliceEventsPaths:
        try:
            dtypes = {'chrom' : 'category', 'chromStart' : 'uint64','chromEnd' : 'uint64','type' : 'category', 'score' : 'float32', 'strand' : 'category'}
            df = pandas.read_csv(i, compression='infer', sep= '\t', names= bsHeader, dtype = dtypes)
            validation = validateBed(df)
            try:
                file_name = i.stem.split('_')[0]+'_'+i.stem.split('_')[1]
            except IndexError:
                file_name = i.stem.split('_')[0]       
            if validation[0]:
                if file_name in spliceEventsDFs:
                    print('Warning, you are using the same prefix for multiple bed files, file ' + str(
                        i) + ' will be ignored')
                else:
                    spliceEventsDFs.update({file_name: df})
                if i.stem.split('_')[0] not in spliceEventNames[1]:
                    try:
                        spliceEventNames[0].append(i.stem.split('_')[1])
                        spliceEventNames[1].append(i.stem.split('_')[0])
                    except:
                        spliceEventNames[1].append(i.stem.split('_')[0])
                        spliceEventNames[0].append(i.stem.split('_')[0])
                for i in df['type'].cat.categories.tolist():
                    if i not in spliceEventTypes:
                        spliceEventTypes.append(i)
            else:
                print('Error in file ' + str(i) + ':')
                print(validation[1])
            validation = None
        except FileNotFoundError:
            print('File ' + str(i) + ' was not found')
        except ValueError as e:
            print('File ' + str(i.stem) + ' had errornous datatypes or missing values, skipping: ' + str(e))                
    if len(spliceEventsPaths) > 0:
        print('Done.')
    if len(spliceEventsDFs) > 0:
        spliceEventsElements += 1
        spliceEventsAvail = True

parser = ArgumentParser(description = '''Interactive, web based visualization for iCLIP and rna-seq data.
                        Atleast one gene annotaiton file in bed12 or gtf format is required for execution.
                        These files need to be unzipped and have the proper file extension.
                        The -bsraw and -bsdata options can be used to provide iCLIP and binding site data
                        to be shown in the visualization. The program will use prefixes to match files 
                        from both options in a one to one relationship. 
                        If multiple files from the same category share a prefix,
                        only the first will be taken into account.
                        For more details on the prefixes consult the help text for the two options.
                        An optional file containing gene descriptions can be provided with -desc,
                        this file should be a tab seperated 4 column csv.
                        The -seqs option can be used to provide fasta files containing dna sequence information.''')
parser.add_argument(dest='geneAnno', help = '''files containing gene annotations in bed12 or gtf format,
                    atleast one such file is required for execution. These files should not include a header.''',
                    nargs = '+', type = Path,metavar = 'GENE ANNOTATION FILE')
parser.add_argument('-bsdata', dest = 'bsdata', help = '''files containing binding site data in 6 column bed format. 
                    Everything before the first underscore _ in the file name will be treated as prefix 
                    and used to match iCLIP files to files containing binding sites.
                    These files should not include a header.''',
                    nargs = '+', type = Path, default = [], metavar = 'FILE')
parser.add_argument('-bsraw', dest = 'bsraw', help = '''files containing iCLIP data, 
                    in .bedgraph format. Everything before the first underscore _ 
                    in the file name will be treated as prefix and used to match 
                    iCLIP files to files containing binding sites. These files should not include a header.''',
                    nargs = '+', type = Path, default = [], metavar = 'FILE')
parser.add_argument('-port', dest = 'port', help = 'port for the dashboard to run on, defaults to 8060',
                    default = 8060, metavar = 'INTEGER', type = int)
parser.add_argument('-desc', dest = 'desc', help = '''file containing gene descriptions,
                    tab seperated csv with 3 columns. This file should not include include a header line,
                    but the column order has to match:
                        gene_id description gene_name''',
                    type = Path, metavar = 'FILE')
parser.add_argument('-seqs', dest = 'fastas', help = '''Fasta files containing genomic sequences,
                    please consult readme for important details''',
                    type = Path, nargs = '+', metavar = 'FILE')
parser.add_argument('-colors', dest = 'colors', 
                    help = '''Colors for the different data sets. Default is a set of 4 colors. 
                    Format(in quotes): rgb(111, 111, 111) Multiple color strings have to be seperated by a sapce.''',
                    nargs = '+', type = str, 
                    default = ['rgb( 88, 24, 69 )', 'rgb( 199, 0, 57 )', 'rgb(46, 214, 26)', 'rgb(255, 87, 51)' ],
                    metavar = 'STRING')
parser.add_argument('-k', dest = 'keys',
                    help = '''Enables you to provide the \'key\' and \'reverse\' arguments
                    for the python standard sort() function. You can use this argment multiple times. 
                    Each key should be provided like this example: -k \'lambda x : x[1:]\' \'False\' ''',
                    nargs = '+',
                    action = 'append')
parser.add_argument('-config' , dest = 'cfg',
                    help = 'An xml config file can be used instead of command lin arguments to specify parameters',
                    type = Path, metavar = 'FILE')
parser.add_argument('-splice_data',
                    dest='splice_data',
                    help = 'files containing splice site data in 4 column bed format.',
                    nargs = '+',
                    default = [],
                    type = Path,
                    metavar = 'FILE')
parser.add_argument('-splice_events',
                    dest='splice_events',
                    help = 'files containing splice events in 6 column bed format.',
                    nargs = '+',
                    default = [],
                    type = Path,
                    metavar = 'FILE')
parser.add_argument('-adv_desc',
                    dest = 'advancedDesc',
                    help = '''Tab seperated file containing additional information
                    on genes. It needs to have a header line and a column named
                    "gene_ids", other columns can be of your choice.
                    For multi value attributes, such as synonyms you can 
                    seperate them by semicolon. Additionally, if you have complex
                    attributes like pulbications, you cam seperate the sub-values by
                    comma and use the -sub_tables argument to create a sub table
                    with specified heaser. Example: author1,year1,title1;author2,
                    year2,title2''',
                    type = Path,
                    metavar = 'FILE')
parser.add_argument('-sub_tables',
                    dest = 'subTables',
                    help = ''''Tab seperated file containing header information to 
                    create sub tables in the advanced description tab. Has two columns,
                    one containing applicable column names from your advanced description file,
                    the other containing semicolon seperated Strings to be used as 
                    header for the sub table. Application example: Publications with author,
                    year, etc. Also see the help for _adv_desc.''',
                    type = Path,
                    metavar = 'FILE')
parser.add_argument('-pswd',
                    dest = 'auth',
                    help = '''Password to access the dashboard''',
                    type = str,
                    default = '',
                    metavar = 'String')
parser.add_argument('-chunksize',
                    dest = 'chunkSize',
                    help = '''Allows specification of the chunk size usedfor subdividing coverage data in lines. Default is 10000''',
                    type = int,
                    default = 10000,
                    metavar = 'Integer')
parser.add_argument('-name',
                    dest = 'name',
                    help = '''Name to create subfolder for binary files''',
                    type = str,
                    default = '',
                    metavar = 'String')
parser.add_argument('-geneindex',
                    dest = 'geneIndex',
                    help = '''Index mapping gene identifiers to transcript identifiers. For use with Bed12 annotation files. Should be two-column
                    tsv file, with the first column containing gene ids and the second column containing transcript ids. No header row.''',
                    type = Path,
                    metavar = 'FILE',
                    )
if __name__ == '__main__':
    args=parser.parse_args()
    
    # Check if xml config file was provided
    useCfg = False
    if args.cfg != None:
        try:
            configFile = minidom.parse(str(args.cfg))
            useCfg = True
        except FileNotFoundError:
            print('Could not open config file, aborting.')
            exit()
        
    if useCfg == False: # Use command line arguments for setup  
        port = args.port
        geneAnnotationPaths = args.geneAnno
        bindingSitePaths = args.bsdata
        bindingSiteRawPaths = args.bsraw
        fastaPaths = args.fastas
        sortKeys = args.keys
        spliceSitePaths = args.splice_data
        spliceEventsPaths = args.splice_events
        chunkSize = args.chunkSize
        try:
            advancedDescPath = Path(args.advancedDesc)
        except TypeError:
            advancedDescPath = None
        try:
            indexPath = Path(args.geneIndex)
        except TypeError:
            indexPath = None
        try:
            subTablePath = Path(args.subTables)
        except TypeError:
            subTablePath = None
        try:
            descriptionPath = Path(args.desc)
        except TypeError:
            descriptionPath = None
        colors = args.colors
        for i in colors:
            if isRGB(i) == True:
                plotColors.append(i)
            else:
                print('Color string ' + str(i) + ' is not valid')
        password = args.auth
        subDir = args.name
    else: # Use xml document for setup
        geneAnnotationPaths = args.geneAnno
        try:
            port = configFile.getElementsByTagName('port')[0].firstChild.data
        except (AttributeError, IndexError):
            print('No port specified, using 8060')
            port = 8060
        try:
            fastaPaths = [Path(i.firstChild.data) for i in configFile.getElementsByTagName('seq')]
        except (AttributeError, IndexError):
            fastaPaths = None
        try:
            descriptionPath = [Path(i.firstChild.data) for i in configFile.getElementsByTagName('desc')][0]
            print(descriptionPath)
        except (AttributeError, IndexError):
            descriptionPath = None
        try:
            sortKeys = []
            keyList = [i for i in configFile.getElementsByTagName('key')]
            for  i in keyList:
                sortKeys.append([i.getElementsByTagName('lambda')[0].firstChild.data,
                                 i.getElementsByTagName('reverse')[0].firstChild.data])
        except (AttributeError, IndexError):
            sortKeys = None
        try:
            dataSetList = [i for i in configFile.getElementsByTagName('set')]
        except (AttributeError, IndexError):
            dataSetList = []
        bindingSitePaths = []
        bindingSiteRawPaths = []
        if len(dataSetList) >= 1:
            for  i in dataSetList:
                try:
                    bindingSiteRawPaths.append(Path(i.getElementsByTagName('clip')[0].firstChild.data))
                except (AttributeError, IndexError):
                    pass
                try:
                    bindingSitePaths.append(Path(i.getElementsByTagName('binding')[0].firstChild.data))
                except (AttributeError, IndexError):
                    pass
                try:
                    color = i.getElementsByTagName('color')[0].firstChild.data
                    if isRGB(color) == True:
                        plotColors.append(color)
                except (AttributeError, IndexError):
                    pass
        try:
            advancedDescPath = Path(configFile.getElementsByTagName('advDesc')[0].firstChild.data)
        except (AttributeError, IndexError):
            advancedDescPath = None
        try:
            subTablePath = Path(configFile.getElementsByTagName('subtables')[0].firstChild.data)
        except (AttributeError, IndexError):
            subTablePath = None
        try:
            spliceSitePaths = [Path(i.firstChild.data) for i in configFile.getElementsByTagName('rnaData')]
        except (AttributeError, IndexError):
            spliceSitePaths = []
        try:
            spliceEventsPaths = [Path(i.firstChild.data) for i in configFile.getElementsByTagName('spliceEvents')]
        except (AttributeError, IndexError):
            spliceEventsPaths = []
        try:
            password = configFile.getElementsByTagName('password')[0].firstChild.data
        except (AttributeError, IndexError):
            password = ''
    
    # Setup directories to store pickles
    if subDir == '':
        binFilePath = os.path.join(os.path.dirname(__file__),'bin_data/')
    else:
        binFilePath = os.path.join(os.path.dirname(__file__),'bin_data/' + subDir +'/')
    if not os.path.exists(os.path.join(os.path.dirname(__file__),'bin_data/')):
        os.mkdir(os.path.join(os.path.dirname(__file__),'bin_data/'))
    if not os.path.exists(binFilePath):
        os.mkdir(binFilePath)
    coveragePath = os.path.join(binFilePath, 'coverage/')
    if not os.path.exists(coveragePath):
        os.mkdir(coveragePath)
        
    # Dict containing checksums for gene annotation files, files loaded once will
    # be serialized to speed up future loading
    try:
        sums = pickle.load(open(binFilePath+'checksums', 'rb'))
    except IOError:
        sums = []
    checksums = dict(sums)
    
    if len(plotColors) == 0:
        print('No valid color strings provided, using defaults')
        plotColors = ['rgb( 88, 24, 69 )', 'rgb( 199, 0, 57 )', 'rgb(46, 214, 26)', 'rgb(255, 87, 51)']        
    
    try:
        geneIndex = pandas.read_csv(indexPath, sep = '\t', compression='infer', comment = '#', names = ['geneID', 'transID'])
    except FileNotFoundError:
        print('Gene index file not found.')
        geneIndex = pandas.DataFrame(columns = ['geneID', 'transID'])
    except ValueError:
        print('geneIndex not set')
        geneIndex = pandas.DataFrame(columns = ['geneID', 'transID'])
    # Load gene annotations from either bed or gtf files. also handle pickling
    loadAnnotations()
    test = [k for k in [i['geneID'].tolist() for i in geneAnnotations]]
    geneNames = list(set().union(*test))
    print('Done.')
    print('Loading description and sequence data if provided.')
    # Read dna sequences from fasta
    loadSequences()
    
    # Read gene descriptions from csv
    loadBasicDescriptions()
        
    # Advanced descriptions for the Details tab
    loadAdvancedDescriptions()
    
    # Subtatbles for the Details tab
    
    loadSubTables
    
    # Setup dropdown with gene descriptions if available
    if descAvail == True:
        geneDict = geneDescriptions.to_dict(orient = 'records')
        # Builds list of gene names and descriptions.
        # Note, in order to be added to the dashboard a gene has to have an entry in the descriptions file
        for i in geneDict:
            dropList.append(
                [i['ensembl_gene_id'] + ' - ' + i['description'][:30] + ' - ' + i['external_gene_name'],
                i['ensembl_gene_id']]
            )
    if descAvail == False or len(dropList) == 0:
        dropList = [[i,i] for i in geneNames]
    dropList.sort(key = lambda x : x[1])
    
    print('Done.')
    # Setup iCLIP data
    loadICLIPData()
    # Setup data for binding sites
    loadBSData()
    
    try:
        coverageSums = pickle.load(open(binFilePath + 'coverage_checksums', 'rb'))
    except IOError:
        coverageSums = []
    coverageChecksums = dict(coverageSums)
        
    
    # Setup data for splice sites
    loadCoverageData()           
    # Write new checksums file
    out = open(binFilePath + 'coverage_checksums', 'wb')
    pickle.dump(coverageChecksums, out)
    out.close()
    
    loadSpliceEvents()
    
    # Keys for sorting of dataset names in iCLIP tab
    if sortKeys == None:
        sortKeys = [['lambda x : x[:1]', 'False']]
    else:
        if len(sortKeys) == 0:
            sortKeys.append(['lambda x : x[:1]', 'False'])
    
    for i in range(len(dataSetNames)):
        colorMap.update({dataSetNames[i] : plotColors[i%len(plotColors)]})
    
    for index, ds in enumerate(sorted(spliceSetNames[1])):
        coverageColorDict.update({ds : coverageColors[index%len(coverageColors)]})
    
    for index, elem in enumerate(sorted(spliceEventTypes)):
        spliceEventColors.update({elem : eventColors[index%len(eventColors)]})
    
    print('preparing to start dashboard on port ' + str(port) + '.')
    
    # Setup gloabl variables for the dashboard
    globalDict = {
        'colorMap' : colorMap, # Contains colors for the data traces
        'descAvail' : descAvail, # Description file present yes/no
        'colorA' : colorA, # Colors for sequence display
        'colorC' : colorC,
        'colorG' : colorG,
        'colorT' : colorT,
        'port' : port, # Port the dashboard runs on
        'procAvail' : procAvail, # Binding site data available True/False
        'dsElements' : dsElements, # Number of elements per dataset, can be 0,1,2
        'spliceElements': spliceElements, # Number of elements per rna dataset, can be 0, 1
        'bsProcDFs' : bsProcDFs, # Dataframes with binding site data
        'bsRawDFs' : bsRawDFs, # Dataframes with iCLIP data
        'dataSetNames' : dataSetNames, # Names for the data sets
        'spliceSetNames' : spliceSetNames, # Names for the rnaSeq data sets
        'rawAvail' : rawAvail, # iCLIP data available True/False
        'spliceAvail' : spliceAvail, # rnaSeq data available True/False
        'dropList' : dropList, # list of entries for the gene selection dropdown
        'geneDescriptions' : geneDescriptions, # dataframe with gene descriptions
        'sequences' : sequences, # list containing sequence files
        'geneAnnotations' : geneAnnotations, # dataframes containing gene annotation data
        'ensembl' : ensembl, # ensembl style fasta format True/False
        'sortKeys' : sortKeys, # arguments for the list.sort function
        'advancedDesc' : advancedDescriptions, # advanced descriptions for Details tab
        'subTables' : subTables, # subtable information for Details tab
        'spliceEventElements' : spliceEventsElements, # Number of elements per rna dataset
        'spliceEventDFs' : spliceEventsDFs,  # Dataframes with splice event data
        'spliceEventNames' : spliceEventNames, # Names for the splice event data sets
        'spliceEventAvail' : spliceEventsAvail, # splice event data available True/False,
        'eventColors' : spliceEventColors, # Colorsfor the splice event types
        'coverageColors' : coverageColorDict, # Colors for the coverage plots
        'eventTypes' : sorted(spliceEventTypes), # List containing types of splice events
        'authentication': password, # Password for authentication
        'coverageData' : fileDict} # Types of splice events
    end = time.time()
    runpy.run_module('app_layout', init_globals = globalDict, run_name = '__main__')
```
