## Supplementary figures and images for "SEQing: web-based visualization of iCLIP and RNA-seq data in an interactive python framework"

### Seqing.png

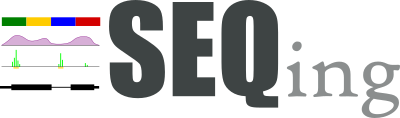

### SEQing_iCLIP_sample.PNG

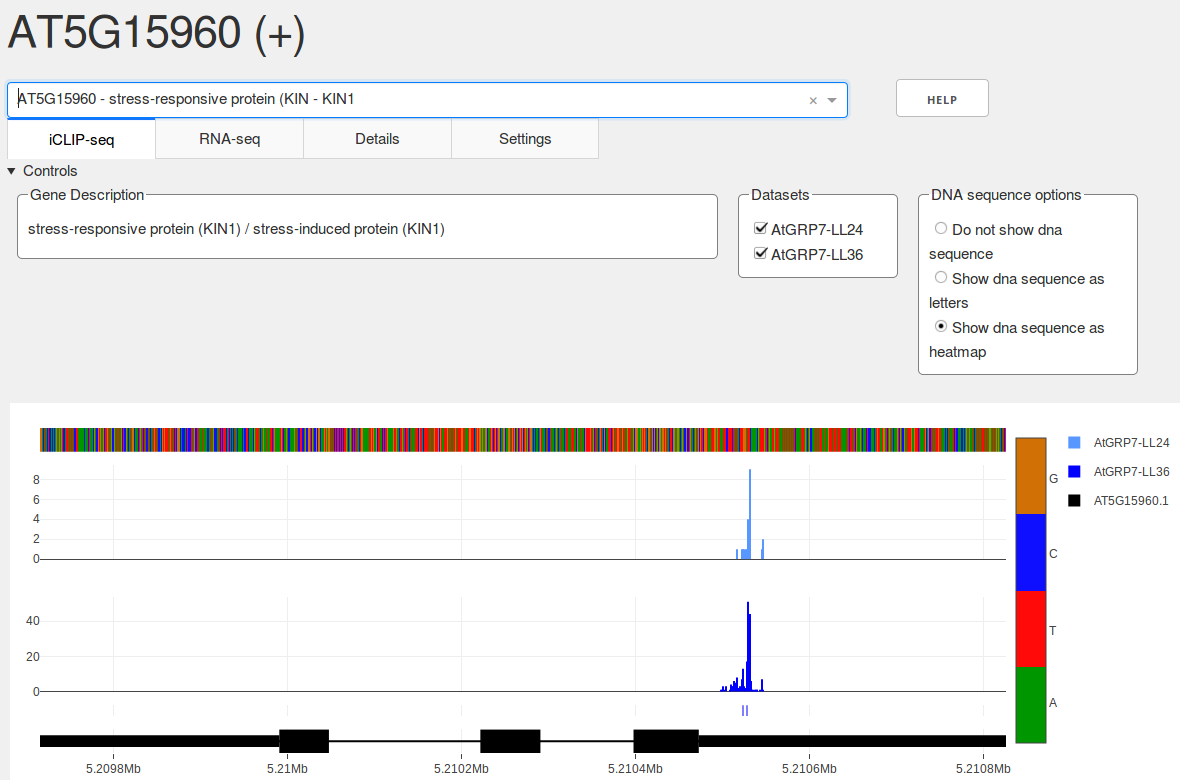

### SEQing_RNA_sample.png

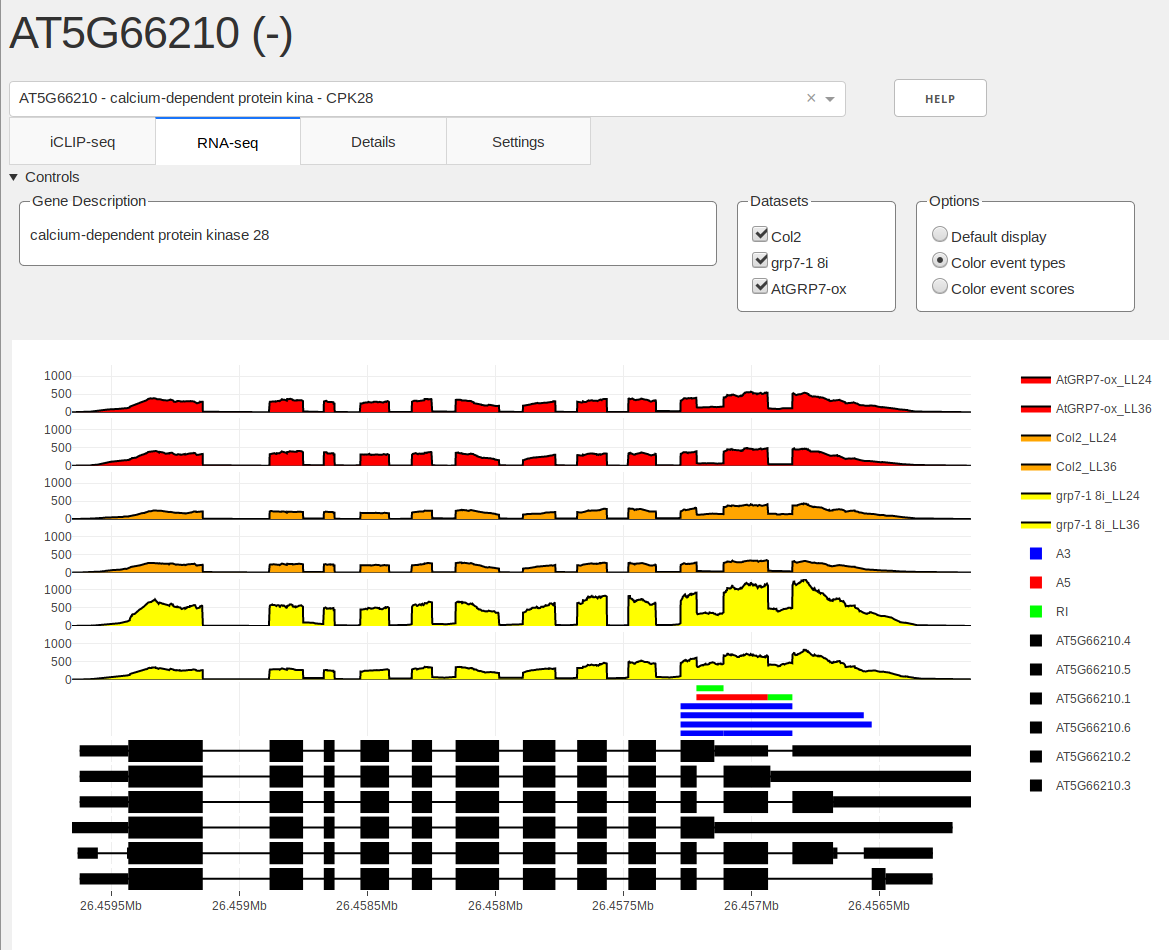
