## Additional File 2 for "SEQing: web-based visualization of iCLIP and RNA-seq data in an interactive python framework"

### SEQing manual

24th February 2020

#### SEQing [seeking]

The goal of this project is to develop a generalized, web-based, interactive visualization and exploration tool for iCLP and RNA-seq data. The application case is a local machine inside the users network, allowing members a web-based (browser) access to explore their experimental data omitting any local installations.

#### Getting Started

These instructions will get you a copy of the project up and running on your local machine.

##### Prerequisites

The source code can be cloned to your local directory using:

```
git clone https://github.com/malewins/SEQing.git
```

or downloaded and extracted from the github page: github.com/malewins/SEQing

The file `requirements.txt` can be used to install all necessary dependencies. Python 3.5 or higher is required and we recommend to setup a virtual environment for this project. If your current python points to a python2 version, please put `python3` instead of just `python` before running SEQing. The same applies to the package installer `pip`.

Once you have setup your virtual environment run the following code to install the dependencies:

```
pip install -r requirements.txt
```

or

```
pip3 install -r requirements.txt
```

if your pip points to an existing Python2 environment.

#### Running SEQing with sample data

After cloning or extraction enter the `example_set` subfolder and run the present bash script:

```
cd example_set
./start_sample.sh
```

This should create the following output:

```
Python3 is installed, starting dashboard ...
Loading gene annotation files.
Loading file 1
Done.
Loading description and sequence data if provided.
Done.
Loading iCLIP data.
Done.
Loading bindings site data.
Done.
Loading RNA-seq data
Col2_LL24
Col2_LL36
grp7-1 8i_LL24
grp7-1 8i_LL36
AtGRP7-ox_LL24
AtGRP7-ox_LL36
Done.
Loading splice event data
Done.
preparing to start dashboard on port 8066.
Running on http://0.0.0.0:8066/
Debugger PIN: XXX-XXX-XXX
```

Now the dashboard with the sample data should be accessible by entering `http://0.0.0.0:8066/` in your browser on the machine or `http://ip-adress:8066/` (e.g. `http://192.168.0.1:8066`) for remote users.

#### Running SEQing

##### Minimal startup

After activation of a virtual environment (env), a minimal startup of the tool can be initiated with the following command:

```
python3 validator.py gene_annotation_file
```

Note that `gene_annotation_file` has to be either a BED12 or GTF file containing gene annotations for the organism. In case of a provided BED12, another file with tab separated gene-id to transcript-id mapping must be provided using the `-geneindex` parameter. It is possible to provide multiple annotation files. In this case the annotation track will be showing every gene. Please ensure that annotation files contain NO header lines (GTF comment lines beginning with '#' are allowed).

On successful initiation the dashboard is accessible via a browser. Genes can be selected from a dropdown on the top to display their associated gene models below.

#### Setting a costomized port

As displayed in the example, the dashboard will be available on a specific port (`preparing to start dashboard on port 8066.`). This port can be set manually using the `-port` parameter:

```
python3 validator.py gene_annotation_file -port 8066
```

Please make sure that you use a valid port number. Otherwise the dashboard will return an error and exit.

#### Loading iCLIP and binding site data

One of the key features of SEQing is the interactive visualization of raw iCLIP crosslink and binding sites. This data should be passed in the form of 4 column bedGraph files using the `-bsraw` parameter:

```
python3 validator.py gene_annotation_file -bsraw WT_iCLIP.bedgraph
```

SEQing will treat everything before the first underscore in the filename as a prefix. This prefix will be used to match a raw iCLIP file to a corresponding binding site file provided with `-bsdata`):

```
python3 validator.py gene_annotation_file -bsraw WT_iCLIP.bedgraph -bsdata WT
_bsites.bed
```

These files must be 6 column BED files. Please note that an iCLIP data sample consists of maximal two files: A mandatory raw iCLIP file (crosslinks) and a binding site file. Multiple datasets can be imported into SEQing by passing them to the corresponding command line options separated by spaces.

#### Loading RNA-seq coverage and splice event data

Besides iCLIP data, SEQing can alsosupports data from RNA-seq, in the form of coverage tracks and bars underneath showing splice events or other regions of interest (events). Coverage data is accepted as 4 column bedGraph format, while events can be provided as 6 column BED files. The following code will load in the files `sample01.bedgraph` and `sample01.bed`:

```
python3 validator.py gene_annotation_file  -splice_data sample01.bedgraph -sp
lice_events sample01.bed
```

The name prefix here works slightly different than it does for the iCLIP data: Files have to share the filename (except file extensions) in order to be associated. An underscore can be used to define a prefix, which can be shared by multiple files and allows them to be (de)selected with one click by the user:

```
python3 validator.py gene_annotation_file  -splice_data sa_sample01.bedgraph
sa_sample02.bedgraph \
-splice_events sa_sample01.bed sa_sample02.bed
```

This will create the category **sa**, which will contain two datasets, **sample01** and **sample02**. Coverage and splice event files still have a one-to-one relationship, i.e. associating multiple event files with a single coverage file is not possible.

#### Displaying gene descriptions and genomic sequences

Apart from visualising iCLIP and binding sites, SEQing is able to provide the user with gene descriptions (`-desc` parameter):

```
python3 validator.py gene_annotation_file -desc description_file
```

The `description_file` needs to be in Tab-separated values format without a header line and consist of 3 columns which are expected in the following order:

```
gene_id description gene_name
```

For this feature SEQing only accepts a single file.

It is possible to provide DNA sequences in FASTA format for the visualisation of a genomic sequence track:

```
python3 validator.py gene_annotation_file -seqs sequence_file
```

Although, there are a few limitations: The fields in the descriptions of the FASTA entries should either be separated by a space or colon, with the first field being the transcript ID:

```
>ENST00000631435.1 cds chromosome:GRCh38:CHR_HSCHR7_2_CTG6:142847306:14284731
7:1
ACGTACGTACGTACGTACGTACGTACGTACGTACGTACGTACGTACGTACGTACGTACGTACGTACGTACGTA
C

>AT1G03993.1::Chr1:23311-24099
ACGTACGTACGTACGTACGTACGTACGTACGTACGTACGTACGTACGTACGTACGTACGTACGTACGTACGTA
C
```

The sequences provided need to cover the whole region of the transcripts as defined in the gene annotations, i.e. the whole genomic sequence in forward direction. SEQing will automatically construct a master sequence if multiple isoforms of the gene are provided, but the individual isoform sequences have to be continuous and not spliced. SEQing will adjust the sequence according to the strand given in the gene annotation to always display in 5’ to 3’ direction.

#### Setting graph colors

The current default color palette consists of four colors. Colors will be reused should more than four datasets be provided. Users can provide customized colors using the parameter -colors:

```
python3 validator.py gene_annotation_file -colors 'rgb(46, 214, 26)' 'rgb(255
, 87, 51)'
```

The colors will be associated to datasets based on the order they are provided.

#### Advanced Description

Using the -adv_desc option it is possible to provide additional information for the genes in your dataset, which will be displayed in the "Details" tab. The parameter takes a single tab separated file which needs to have a header line and a column named gene_ids. Another constraint is that the file should only contain one row of data per gene, thus multi value attributes such as synonyms need to be combined into one, semicolon delimited field. All other columns can have names of your choosing. Example:

```
gene_ids    synonyms
AT1G01010   ANAC001;NAC domain containing protein 1
```

As an additional feature it is possible to create subtables for complex attributes like publications. To provide author, year and title information for a gene, it is necessary to use additional comma based separations and the -sub_tables option to create a tabular view of your publication data. An example for the field in your advanced description file:

```
gene_ids    publications
AT1G10670   author1,year1,title1;author2,year2,title2
```

As demonstrated, comma and semicolon delimiters can be mixed in one field. The -sub_tables option takes an additional file which describes the table for the specific attribute. The structure is consists of two tab delimited columns, one containing the attribute names of the subtables for and the other containing the semicolon delimited column headers for the table:

```
publications    Author;Year;Title
```

This file does not need a header line. Should the number of columns not match the number of comma separated entries in your advanced description file the data will be displayed as the comma separated string without a table.

Finally you can use ? before a value to mark it as a hyperlink:

```
gene_ids    ext_link
AT1G01010   ?http://araport.org
```

This will also work for semicolon separated values as well as in subtables, but not in comma separated Strings.

#### Restricting access to the dashboard

In the case of data that is not meant to be openly accessible, even over a local network, it is possible to specify a password using the `-pswd` parameter. Users will then have to enter this password before they can access the dashboard. The application will use a hardcoded username, which is “u”, in conjunction with the password provided.

#### Setting the name of your instance

SEQing can be started multiple times (instance) from the same folder. Each instance is set to use the standard `bin_data` directory to store binary files. To omit overwriting of other instances, a name for the current instance can be defined on start using the `-name` parameter:

```
python3 validator.py gene_annotation_file -name ath_iclip
```

This is only relevant, if you re-use files in separate instances of SEQing.

#### Screenshots

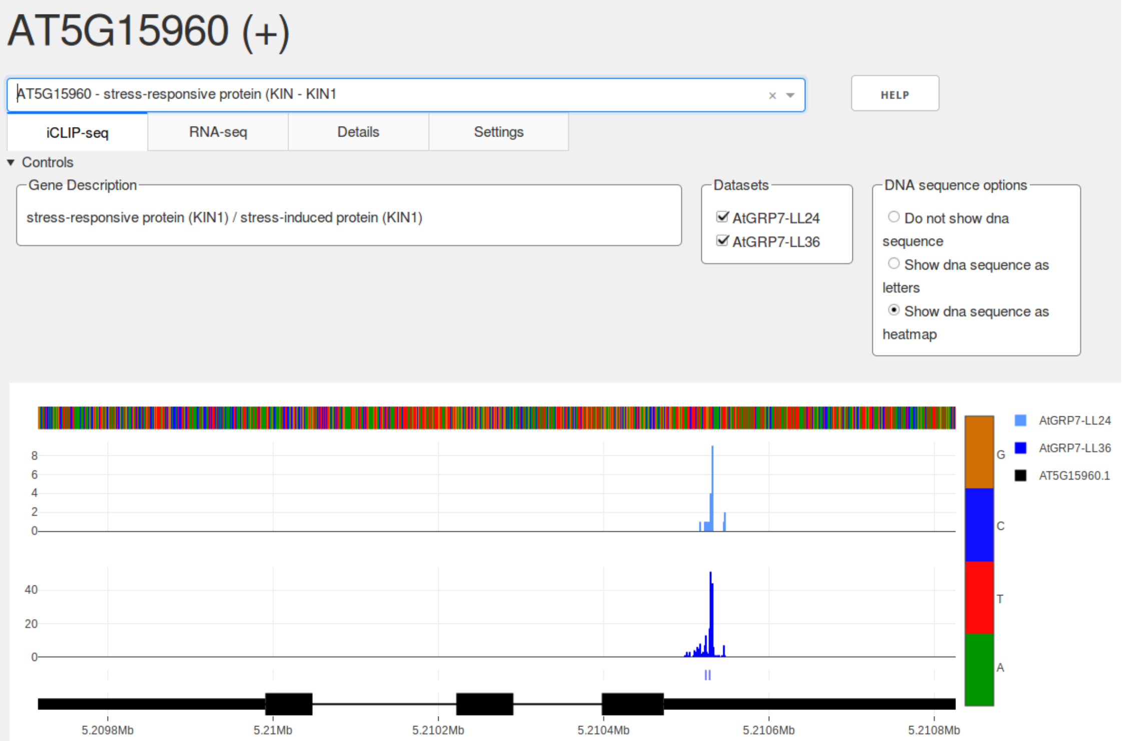


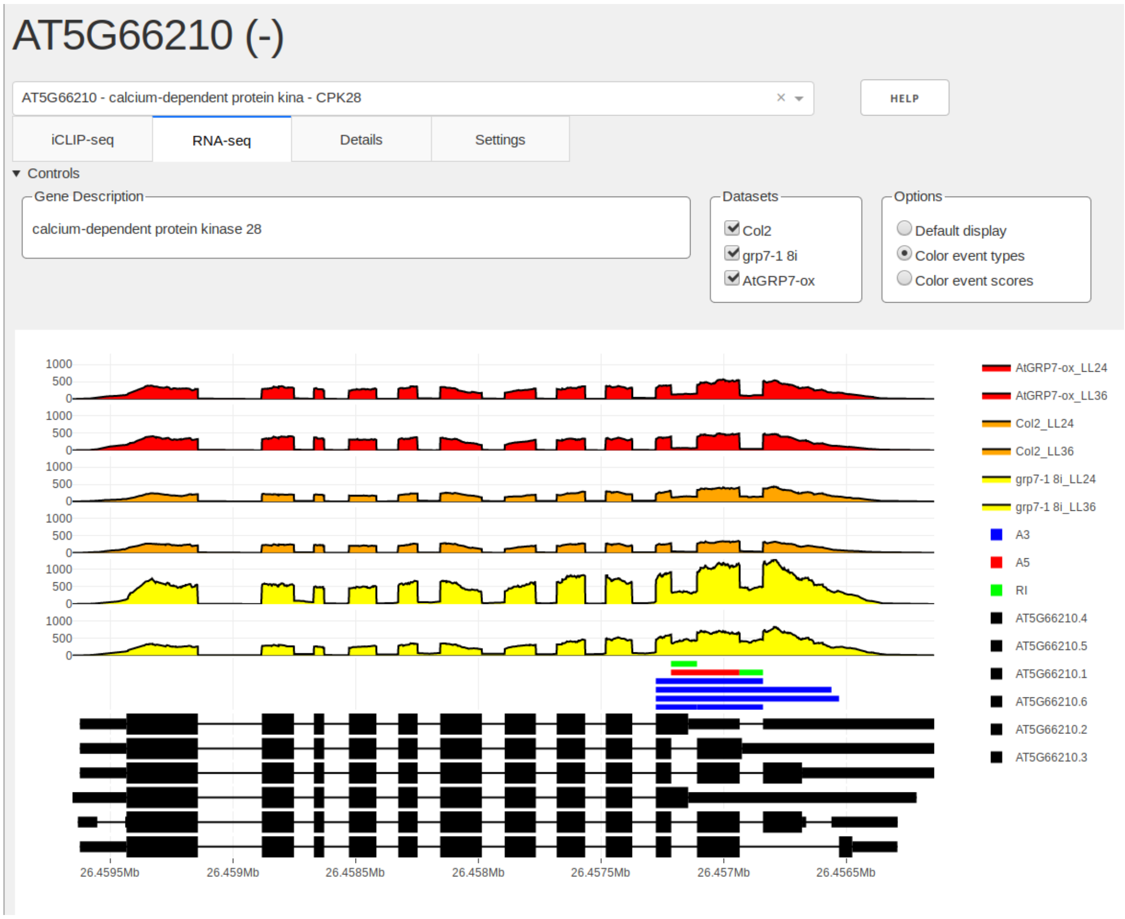

#### Input file format overview

The following is a quick overview over all files types used as inputs and their respective requirements:

| Input | Description |
|---|---|
| Gene annotations | BED12 or GTF files that conform to the corresponding standards. Do not include header rows |
| iCLIP raw data | BEDGRAPH files conforming to the standard. No header rows |
| Binding Site data | BED6 files conforming to the standard. No header rows |
| Coverage data | BEDGRAPH files conforming to the standard. No header rows |
| Splice event data | BED6 files conforming to the standard. No header rows |
| Basic description file | Tab separated file with 3 columns. No header. Columns are expected in order: gene_id, description, gene_name |
| Advanced description file | Tab separated file. Has to contain a header row and a column named "gene_ids". All other columns can be custom |
| Subtable file | 2 column tab separated file, first column contains column names from advanced descriptions, second column contains subtable column names separated by semicolon. No header |

#### Sorting

Arranging data tracks of multiple datasets is done by providing a `-k` parameter to provide SEQing with regular expressions for sorting. Graphs for datasets can be toggled on and off in the visualisation. Depending on the way the user does this, the order in which datasets are displayed may change. Therefore the graphs are sorted by default in ascending order using the prefix. However you might have a need for more complex sorting, like the following example:

```
python3 validator.py gene_annotation_file -bsraw 7pref26_iCLIP 7pref30_iCLIP \
-k 'lambda x : x[-2:]' 'True' -k 'lambda x : x[:1]' 'False'
```

The `-k` parameter allows users to directly provide arguments for the `list.sort` function of python. In this case we provide 2 different sets of arguments, the first one sorts the prefixes by the last two characters (descending order) and the second sorts them by the first character in ascending order. Each -k has to be followed by a string containing the desired lambda expression and a boolean telling the program whether to revert the order (the default is *ascending*).

#### License

MIT
