## Additional File 3 for "SEQing: web-based visualization of iCLIP and RNA-seq data in an interactive python framework"

### Supplemental Figures

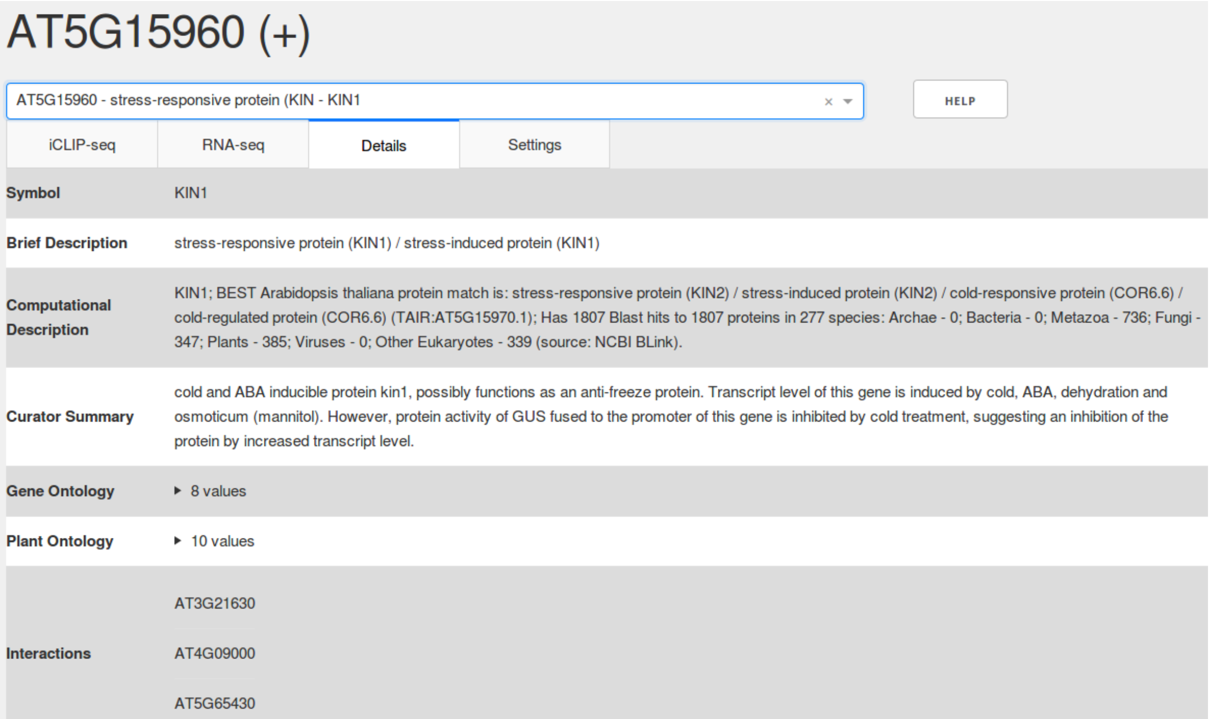


**Supplemental Figure 1: Screenshot of the SEQing Details tab.** This separate tab displays extended information about the selected gene in tabular format and delivers background information directly through SEQing. These data can be loaded into SEQing by specifying a Tab-separated values file using the parameter -adv_desc.

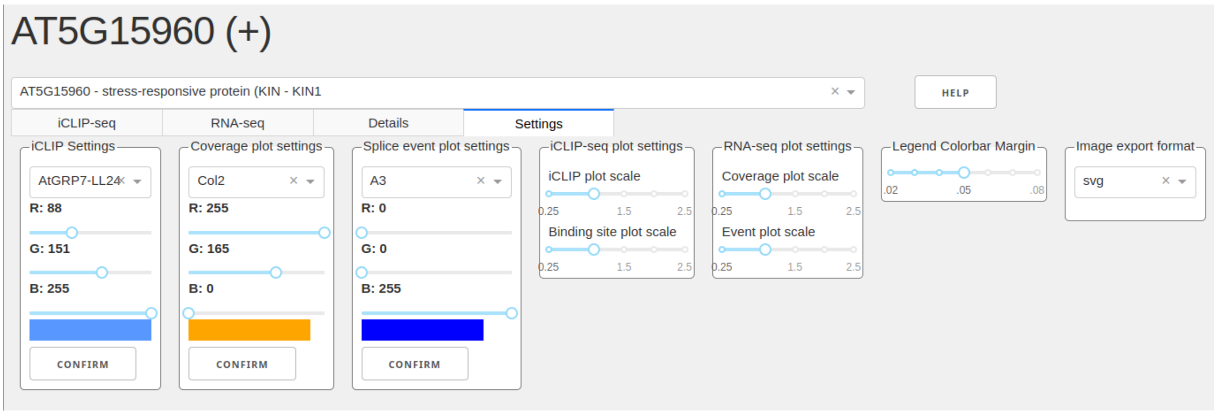


**Supplemental Figure 2: Screenshot of the SEQing Settings tab.** The display options for each data track can be modified manually in this tab. Track colors can be set individually for each data track, whereas track heights are adjustable globally for the iCLIP and RNA-seq tab. The file format in which the images should be exported is switchable between .png and .svg. The .svg export is the default setting.

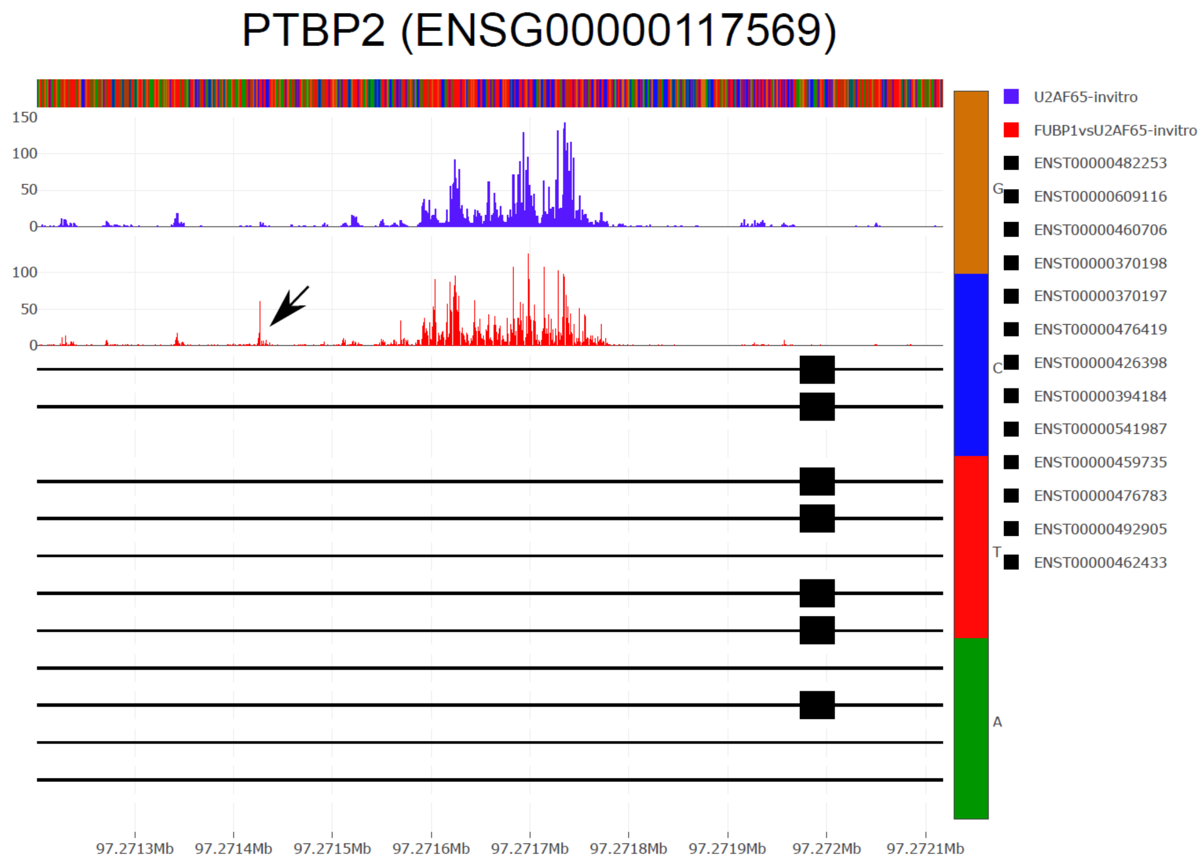


**Supplemental Figure 3: Visualization of U2 Auxiliary Factor 2 (U2AF2) crosslink sites on the Polypyrimidine Tract Binding Protein 2 (PTBP2) gene using SEQing.** The exported image, created by SEQing, depicts a genomic window from one of the U2AF2 targets, Polypyrimidine Tract Binding Protein 2 (PTBP2). The data tracks show *in vitro* crosslink sites from U2AF2 alone and from U2AF2 together with the cofactor Far Upstream Element Binding Protein 1 (FUBP1). It is demonstrated that the presence of the FUBP1 protein influences U2AF2 binding. The area marked by the arrow points to a U2AF2 binding site in the presence of FUBP1 protein. Below the data tracks reside known transcript annotations of the PTBP2 gene. A track legend, color coding of the bases and transcript isoform accessions are featured on the right side of the exported image.
